# On the complex interplay between spectral harmonicity and different types of cross frequency couplings in non linear oscillators and biologically plausible neural network models

**DOI:** 10.1101/2020.10.15.341800

**Authors:** Damián Dellavale, Osvaldo Matías Velarde, Germán Mato, Eugenio Urdapilleta

## Abstract

**Background:** Cross-frequency coupling (CFC) refers to the non linear interaction between oscillations in different frequency bands, and it is a rather ubiquitous phenomenon that has been observed in a variety of physical and biophysical systems. In particular, the coupling between the phase of slow oscillations and the amplitude of fast oscillations, referred as phase-amplitude coupling (PAC), has been intensively explored in the brain activity recorded from animals and humans. However, the interpretation of these CFC patterns remains challenging since harmonic spectral correlations characterizing non sinusoidal oscillatory dynamics can act as a confounding factor.

**Methods:** Specialized signal processing techniques are proposed to address the complex interplay between spectral harmonicity and different types of CFC, not restricted only to PAC. For this, we provide an in-depth characterization of the Time Locked Index (TLI) as a novel tool aimed to efficiently quantify the harmonic content of noisy time series. It is shown that the proposed TLI measure is more robust and outperform traditional phase coherence metrics (e.g. Phase Locking Value, Pairwise Phase Consistency) in several aspects.

**Results:** We found that a non linear oscillator under the effect of additive noise can produce spurious CFC with low spectral harmonic content. On the other hand, two coupled oscillatory dynamics with independent fundamental frequencies can produce true CFC with high spectral harmonic content via a rectification mechanism or other post-interaction nonlinear processing mechanisms. These results reveal a complex interplay between CFC and harmonicity emerging in the dynamics of biologically plausible neural network models and more generic non linear and parametric oscillators.

**Conclusions:** We show that, contrary to what is usually assumed in the literature, the high harmonic content observed in non sinusoidal oscillatory dynamics, is neither sufficient nor necessary condition to interpret the associated CFC patterns as epiphenomenal. There is mounting evidence suggesting that the combination of multimodal recordings, specialized signal processing techniques and theoretical modeling is becoming a required step to completely understand CFC patterns observed in oscillatory rich dynamics of physical and biophysical systems.

**Highlights:**

- Time locked index efficiently quantifies the harmonic content of noisy time series.
- A non linear oscillator under the effect of additive noise can produce spurious cross frequency couplings (CFC) with low spectral harmonic content.
- Two coupled oscillatory dynamics with independent fundamental frequencies can produce true CFC with high spectral harmonic content via rectification mechanisms or other post-interaction nonlinear processing mechanisms.
- A non sinusoidal oscillatory dynamics with high harmonic content is neither sufficient nor necessary condition for spurious CFC.
- A complex interplay between CFC and harmonicity emerges from the dynamics of nonlinear, parametric and biologically plausible oscillators.

## 1. INTRODUCTION

One of the most challenging and active topics in signal processing research refers to tackling the inverse problem associated to infer the underlying mechanisms producing the time series observed from a given physical system. This is particularly true in electrophysiologically based Systems Neuroscience, in which a long standing goal is to infer the underlying multidimensional neural dynamics from spatially sparse low dimensional recordings [1]. Cross frequency coupling (CFC) is a signature observed at the signal level informative on the mechanisms underlying the oscillatory dynamics. From the signal processing point of view, a CFC pattern emerges when certain characteristics (e.g. amplitude, phase) of a frequency band interact with others in a different band, either in the same signal or in another related one. CFC is a rather ubiquitous phenomenon observed in the oscillatory dynamics of a variety of physical and biophysical systems (see [1] and references therein). In particular, the phase-amplitude cross frequency coupling (PAC) observed in the electrical oscillatory activity of animal and human brains, has been proposed to be functionally involved in neuronal communication, memory formation and learning. This has motivated the development of specialized signal processing algorithms to robustly detect and quantify PAC patterns from noisy neural recordings [2, 3, 4, 5, 6, 7]. However, the interpretation of these PAC patterns remains challenging due to the fact that harmonic spectral correlations characterizing non sinusoidal oscillatory dynamics can act as a confounding factor.

The concept of harmonicity refers to the degree of commensurability between the periods of the rhythms constituting the analyzed oscillatory dynamics. More precisely, two frequencies f_2_ and f_1_ are commensurable if they satisfy f_2_/f_1_ ∈ ℚ, while harmonic frequencies are related by an integer ratio, i.e. for f_2_ > f_1_ they satisfy f_2_/f_1_∈ ℤ. In terms of the Fourier analysis, harmonicity can be thought as the amount of spectral power concentrated at harmonic frequencies (i.e. spectral harmonicity). In the context of CFC, harmonicity is measured as the degree of phase synchronization between rhythms pertaining to different frequency bands (i.e. PPC: phase-phase cross frequency coupling). Thus, the proposed harmonicity analysis would be worthy in many fields involving the study of physical and biophysical systems through low dimensional time series. For instance, the quantification of spectral harmonicity associated to non sinusoidal neural oscillations can serve as a measure of spatial synchronization in non invasive brain recordings like scalp electroencephalography (EEG) and magnetoencephalography (MEG) [8]. In addition, the on and off medication states of patients with Parkinson’s disease can be distinguished by means of the non sinusoidal waveform shape of the exaggerated beta band oscillations observed in invasive (intracerebral EEG) and non invasive (scalp EEG) recordings [9, 10]. Moreover, the harmonicity observed in intracerebral EEG recorded from epilepsy patients has been used to effectively distinguish between harmonic and non harmonic PAC patterns putatively linked to two essentially different mechanisms of seizure propagation [11]. Furthermore, when two oscillatory inputs converge in a nonlinear integrator (e.g. a neuron), new harmonic and non harmonic (a.k.a. emergent) oscillations are generated via the frequency mixing mechanism [12]. Importantly, oscillations emerging from this mechanism entrain unit activity [12], suggesting that frequency mixing is intrinsic to the structure of spontaneous neural activity and contributes significantly to neural dynamics. Recently, it has been reported that frequency mixing is widely expressed in a state and region-dependent manner in cortical and subcortical structures in rats [12]. In this context, spectral harmonicity could be used as a surrogate of the frequency mixing mechanism and emergent components entraining unit activity, thus, our proposed method for harmonicity quantification complements the tools described in [12]. The evidence discussed above suggest that the quantification of harmonicity of invasive and non invasive neural recordings from humans and animal models can be used as a biomarker to characterize physiological and also pathological brain states like those observed in Parkinson’s disease and epilepsy. In this work we provide an in-depth characterization of the Time Locked Index (TLI) as a novel tool aimed to efficiently quantify the harmonic content of noisy time series. In addition, the TLI is used together with other proposed signal processing techniques to quantitatively analyze the complex interplay between spectral harmonicity and different types of CFC patterns, not restricted only to PAC.

## 2. METHODS

### 2.1. Synthetic and simulated dynamics

The spectral harmonicity and CFC patterns were analyzed in a variety of synthetic and simulated oscillatory dynamics in presence of intrinsic and extrinsic additive noise. In Appendix A.1 it is described the formulation used to synthesize amplitude-modulated time series. Appendix A.2 and Appendix A.3, provide the equations for simulating the dynamics associated to the Van der Pol oscillator and a 2nd order parametric oscillator, respectively.

In what follows we define an analytically tractable model capable to produce unidirectional PAC with external drive. In this model the slow rhythm represents an external sensory input modulating the fast oscillations in sensory circuits (see discussion in [13]). The characteristics of the oscillatory dynamics and the PAC patterns elicited by the proposed biologically plausible neural network architecture have been extensively analyzed in our previous works [1, 14]. In brief, the model consists of a single excitatory and a single inhibitory population that are reciprocally connected (Figure 1). This representation follows the model introduced in [15], it is a minimal version of a system capable of generating oscillations [15, 16]. The dynamics of the two populations can be written as,

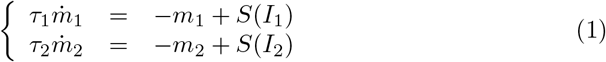

where *m_i_* and *τ_i_* represent the output of the population *i* ∈ {1,2} and the time constant, respectively. The output of this representation is constituted by the currents *I*_1_ = *G*_2_*m*_2_(*t*-Δ_2_) + *H*_1_+*η*_1_ and *I*_2_ = *G*_1_*m*_1_(*t*-Δ_1_) +*H*_2_+*η*_2_, where *G_i_* indicates the efficacy of the interactions, *H_i_* is a external input and Δ*_i_* are delays in the transmission of the interaction. The terms *η_i_* represent additive white Gaussian noise (AWGN) to the inputs *I_i_*. More precisely, 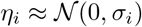.

**Figure 1:**
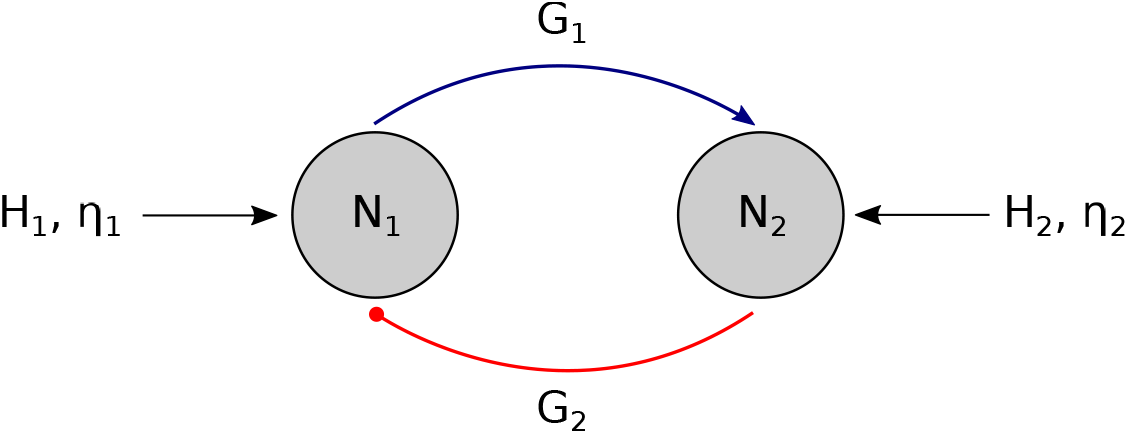
Biologically plausible network for unidirectional PAC with external drive representing a slow sensory input entraining fast oscillations underpinning local neural processing in a cortical oscillator (Sensory entrainment).

Regarding the instantaneous activity *A_i_ = S(I_i_)* of both populations, in Eqs. 1 we consider threshold linear *S(I_i_)* and softplus *S_c_(I_i_)* transfer functions defined as,

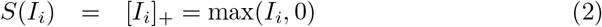

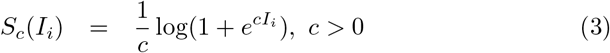

The softplus transfer function in Eq. 3 results *S_c_(I_i_)* > 0 and converges toward the threshold linear transfer in the limit *c* → ∞. However, these two transfer functions are essentially different regarding their order of continuity, being *S(I_i_)* of class *C^0^* (continuous but not differentiable) and *S_c_(I_i_)* of class *C*^∞^ since it is infinitely differentiable. This has rather profound implications in the resulting dynamics. For instance, the stability of the stationary state depends on the activation function as well as of its derivative (see Eq. 4 in [1]). As a consequence, *S(I_i_)* and *S_c_(I_i_)* can produce very different stability conditions even when the latter converge to the former in the limit c → ∞. A discussion on how the activation functions constituting the biologically plausible model affect the CFC patterns emerging in the resulting oscillatory dynamics is presented in Section 3.4.

Synaptic efficacies *G_i_* were imposed so that the system was in the oscillatory state. The resulting oscillatory activity at 50 Hz belongs to the gamma band. All the parameters for network are summarized in Table 1.

**Table 1:** Values of the coupling parameters, time constants and delays for the model shown in Figure 1.

|  | Synaptic efficacy | Delay | Time constant |
| --- | --- | --- | --- |
| | $G$ | $\Delta$ [ms] | $\tau$ [ms] |
| 1→2 | 1.4 | 5 | 0.1 |
| 2→1 | -1 | 5 | 0.1 |

### 2.2. Power Spectral Density

Power spectral density (PSD) estimates were computed using the modified periodogram method with a Hann window in the time domain [17].

### 2.3. Cross Frequency Coupling

To quantify cross frequency coupling (CFC) patterns observed in the explored oscillatory dynamics, non parametric methods were used: Phase Locking Value (PLV), the Mean Vector Length (MVL) and the Modulation Index based on the Kullback-Leibler distance (KLMI) (see [6] and references therein). In the particular case of PAC, Figure 3 shows for two synthetic oscillatory dynamics the raw time series (*x*(*t*)) together with the band-pass filtered signals (*x_LF_*(*t*), *x_HF_*(*t*)), the phase of the low frequency signal (*ϕ_LF_*(*t*)) and, the amplitude envelope of the high frequency signal (*a_HF_*(*t*)), as well as its phase evolution (*ϕ_a_*_HF_(*t*)) from which the PLV, MVL and KLMI metrics can be computed as follows [2, 4, 6],

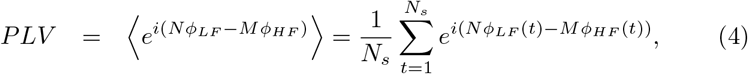

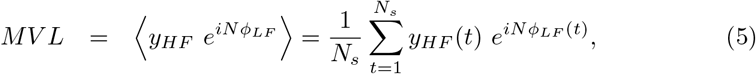

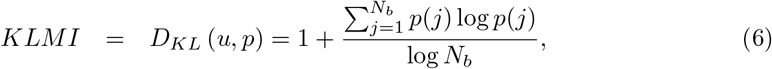

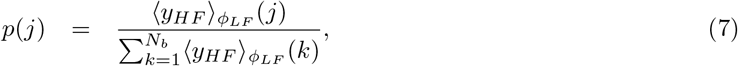

where *t*∈ℤ is the discrete time index, *i* is the imaginary unit, *N* and *M* are some integers, *N_s_* is the number of samples of the time series, *p*(*j*) denotes the mean *y_HF_(t)* value at the *ϕ_LF_(t)* phase bin 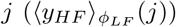 normalized by the sum over the bins (see histograms in Figures 3C1,C2), *Nb* is the number of bins for the phase histogram and *D_KL_* represents the Kullback-Leibler distance between *p* and the uniform distribution *u*. In the case of PAC (see Figure 3), Eqs. 4 to 7 are computed using *y_HF_*(*t*) = *a_HF_*(*t*) and *ϕ_LF_*(*t*) = *ϕ_aHF_*(*t*). It is worth noting that PLV, MVL and KLMI metrics have been extensively used to quantify PPC and PAC, however, they can also be used to quantify other CFC types like AAC and PFC after replacing *ϕ_LF_*(*t*), *ϕ_HF_*(*t*) and *y_HF_*(*t*) with the appropriate time series. A detailed discussion regarding the proper configuration and processing of the time series involved in the quantification of several CFC types including those explored in this work is given in Appendix A.4.

One of the main confounds when assessing PAC is related to the nonuniform distribution of phase angles of the modulating component *x_LF_*(*t*), which can produce spurious PAC levels [18]. To detect the occurrence of this confound we computed the phase clustering (PC) as shown in Eq. 8 (see Chapter 30, p. 414 in [17]),

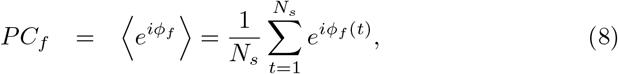

where *ϕ_f_(t)* is computed as described in Appendix A.4 and the subscript for the frequency band of interest is defined as *f ∈ {LF, HF}*. When *x_LF_(t)* has a periodic sinusoidal-like waveform shape, we obtain a rather uniform phase angle distribution *ϕ_LF_(t)* resulting in |*PC_LF_*| ≈ 0 for a sufficiently large number of samples Ns. On the other hand, if the time series *x_LF_(t)* is highly non sinusoidal, we obtain a skewed distribution of phase angles producing |*PC_LF_*| ≈ 1. Worthy to note, the spurious PAC associated to high PC values can be mitigated by using narrow enough band-pass filter (BPF) to obtain the modulating low frequency oscillations *x_LF_(t)* (see the time series *ϕ_LF_(t)* in Figures 3C1,C2), or by the method described in [18]. In contrast, to effectively assess PAC, the BPF aimed to obtain the modulated high frequency oscillations *x_HF_(t)* must satisfy the restriction related to the minimum bandwidth determined by the low frequency band: 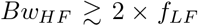, where *f_LF_* is the center frequency of the BPF for *x_LF_(t)* [19].

### 2.4. Time Locked Index

A specialized tool was developed to characterize the spectral harmonicity associated to the CFC patterns observed in the explored oscillatory dynamics [1, 11]. Specifically, the Time Locked Index (TLI) was implemented to efficiently quantify the presence of spectral harmonics associated to the emergence of CFC in noisy signals. The quantitative characterization of the harmonicity of the oscillatory dynamics is important given that coupled oscillatory dynamics characterized by independent frequencies or non sinusoidal repetitive waveform shapes can both elicit a similar signature in the Fourier spectrum. In particular, the traditional algorithms aimed to assess CFC based on linear filtering (e.g. PLV, MVL, KLMI) are confounded by harmonically related spectral components associated to non sinusoidal pseudoperiodic waveform shapes, reporting significant CFC levels in absence of independent frequency bands [1, 20, 21]. In the TLI algorithm, time-locked averages are implemented in the time domain to exploit the phase synchronization between harmonically related spectral components constituting the non sinusoidal oscillatory dynamics. The following steps describe the procedure to compute TLI (see Figures 3B1,B2),

1. The input signal x is band-pass filtered at the low (LF) and high (HF) frequency bands under analysis, producing the time series *x_LF_* and xHF, respectively. Z-score normalization is applied on the time series *x_LF_*and *x_HF_* to ensure the TLI metric is independent of the signals amplitude.
2. The time instants corresponding to the maximum amplitude (or any other particular phase) of both time series, *x_LF_*and *x_HF_*, are identified in each period of the low frequency band (*T_LF_*). These time values for the slow and fast oscillation peaks are recorded in the time vectors *t_LF_* (red down-pointing triangles in Figures 3B1,B2) and *t_HF_* (green up-pointing triangles in Figures 3B1,B2), respectively.
3. Epochs 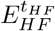 with a length equal to one period of the low frequency band (*T_LF_*) centered at the fast oscillation peaks (*t_HF_*) are extracted form the time series *x_HF_*. Averaging over these epochs is computed to produce a mean epoch 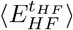. Note that the latter is a time-locked averaging due to the fact that every single epoch 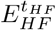 is centered at the corresponding time instant *t_HF_*.
4. Epochs 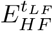 with a length equal to one period of the low frequency band (*T_LF_*) centered at slow oscillation peaks (*T_LF_*) are extracted form the time series *x_HF_*. Averaging over these epochs is computed to produce a mean epoch 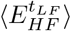. Note that the latter is also a time-locked averaging, now with epochs centered at the corresponding time instants *T_LF_*.
5. Finally, the TLI is computed as follows,

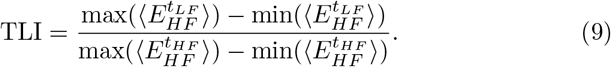

In the case that the time series *x* were predominantly constituted by harmonic spectral components, the fast (*x_HF_*) and slow (*x_LF_*) oscillatory dynamics are characterized by a high degree of synchronization in time domain (i.e. phaselocking). As a consequence, the amplitude of 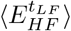 results comparable to that of the 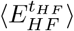 and so we obtain TLI ≈ 1 (see Figures 3A1,B1). On the other hand, if the spectral energy of the time series *x* is not concentrated in narrow harmonically related frequency bands, the fast (*x_HF_*) and slow (*x_LF_*) rhythms will be not, in general, phase-locked. Therefore, the amplitude of 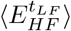 is averaged out to zero and TLI ≈ 0 is obtained for a sufficiently large number of samples *N_s_* (see Figures 3A2,B2).

It is worth noting that the phase synchronization between the band-pass filtered time series (*x_LF_* and *x_HF_*) can be quantified using the PLV metric, however, the TLI algorithm has two significant advantages: 1) The computation of the TLI measure does not require to know the harmonic ratio between the frequency bands of interest. In contrast, to compute the PLV one needs to know this harmonic ratio (i.e. the values of the integers *N* and *M* in Eq. 4), a priori, in order to be able to evaluate the phase-phase cross frequency coupling characterizing the harmonic spectral components [3]. 2) The TLI metric can be effectively computed using slightly selective BPF to obtain the HF component *x_HF_*(*t*), i.e. filters having wide bandwidths or low steepness of the transition bands. That is, by operating in the time domain the TLI reliably assesses the degree of time-locking, even in the case in which several (harmonic) spectral components are included within the bandwidth of the filter used to obtain the fast rhythm (*x_HF_*). This specific capability of the TLI metric is illustrated and further discussed below in connection with Figures 5, 8, 13 and 14.

Even though the TLI is a measure bounded in the range [0,1] (see Section 2.4) and independent of the processed oscillations amplitude, the absolute value of the TLI does depend on the noise level present in the processed time series and on the epoch length, i.e. the number of periods of the low frequency oscillation taken to implement the time-locked average involved in the TLI computation (this is further discussed below in connection with Figures 5 to 8 and B.1). As a consequence, the TLI is not a bias-free measure and this issue must be taken into account to implement a quantitative analysis of harmonicity. Fortunately, the surrogate control analysis [6] based on sample shuffled *x_HF_*(*t*) time series described in Section 2.7 below, overcome this limitation. Besides, in the case of time series corresponding to multiple channels (e.g. multi-site recordings) or trials, a commonly used method to remove the bias is to implement a Z-score normalization across channels/trials (i.e. spatial whitening) [11].

In contrast to the PLV and TLI which are biased measures [22], the pairwise phase consistency is a bias-free metric suitable for quantifying phase-phase coupling [23, 24]. However, it should be noted that the number of arithmetic operations involved in the computation of the PLV and TLI increase linearly with the number of samples *N_s_* (i.e. computational complexity of *O*(*N_s_*)), while the pairwise phase consistency measure presents a significantly higher computational complexity of *O*(*N_s_*^2^). Another measure commonly used to assess phase synchronization is the spectral coherence (see [17], Section 26.7, p. 342). Importantly, although the definition of spectral coherence includes a normalization by the total power to produce a bounded metric in the range [0, 1], in the expression of spectral coherence individual phase angle vectors are weighted by power values. Therefore, results from spectral coherence are likely to be influenced by strong increases or decreases in power ([17, 25]). In other words, the spectral coherence is sensitive to phase-phase and also to amplitude-amplitude and phase-amplitude correlations between the input signals. On the other hand, the TLI measure is defined as the ratio of time-locked averages computed on the same signal (HF oscillations *x_HF_*), as such, it results an amplitude independent quantity only depending on the degree of synchronization between the sequence of time instants used to compute these time-locked averages (*t_LF_* and *t_HF_*). As a result, the TLI metric is sensitive only to PPC between the input rhythms.

The source code for the computation of TLI together with test script examples implemented in Matlab® and Python are freely available at, https://github.com/damian-dellavale/Time-Locked-Index/.

We are willing to provide technical support to investigators who express an interest in implementing the TLI metric in other programming languages, integrate it in open-source software toolboxes or use it for non-profit research activities. The potential of the TLI metric to improve the characterization and aid the interpretation of PAC patterns observed in invasive neural recordings obtained from epileptic patients and in simulated dynamics of biologically plausible networks, has been demonstrated in our previous works [1, 11]. In this paper we extent the harmonicity analysis to four types of CFC patterns including an indepth characterization of the TLI performance using simulated and synthetic oscillatory dynamics under controlled levels of intrinsic noise (AWGN: additive white Gaussian noise).

Figure 2 shows two essentially different CFC scenarios in terms of the spectral harmonicity, however, they are indistinguishable by traditional metrics aimed to assess CFC based on band-pass linear filtering (e.g. PLV, MVL, KLMI). Figure 2A shows phase-amplitude coupling via harmonic content. In terms of telecoms engineering, the harmonic content constituting the spectrum of a quasi-periodic non sinusoidal waveform with fundamental frequency *f*_0_ can be though as a ‘carrier’ given by the harmonic *N f_0_*, being *N*∈ℤ the harmonic number, and ‘sidebands’ (*N* - 1)*f*_0_, (*N* + 1)*f*_0_. This spectral profile is known to produce CFC patterns in the time domain (e.g, an amplitude-modulated signal). This kind of CFC patterns will be referred as ‘harmonic’ CFC. Figure 2B shows phaseamplitude coupling in absence of phase-phase cross frequency coupling between the ‘sidebands’ and the ‘carrier’. That is, the ‘sidebands’ are not harmonics of the ‘carrier’ (non harmonic frequencies). This kind of CFC patterns will be referred as ‘non harmonic’ CFC. Importantly, traditional algorithms aimed to assess CFC (e.g. PLV, MVL, KLMI) are confounded by harmonically related spectral components associated to a single (quasi)periodic non sinusoidal dynamics, reporting significant CFC levels even in absence of underlying coupled dynamics (i.e. spurious CFC). This is due to the fact that coupled oscillatory dynamics characterized by independent frequencies (i.e. true CFC) and a single non sinusoidal oscillatory dynamics (i.e. spurious CFC) produce similar signatures in the Fourier spectrum that are hardly distinguishable by using band-pass linear filtering.

**Figure 2:**
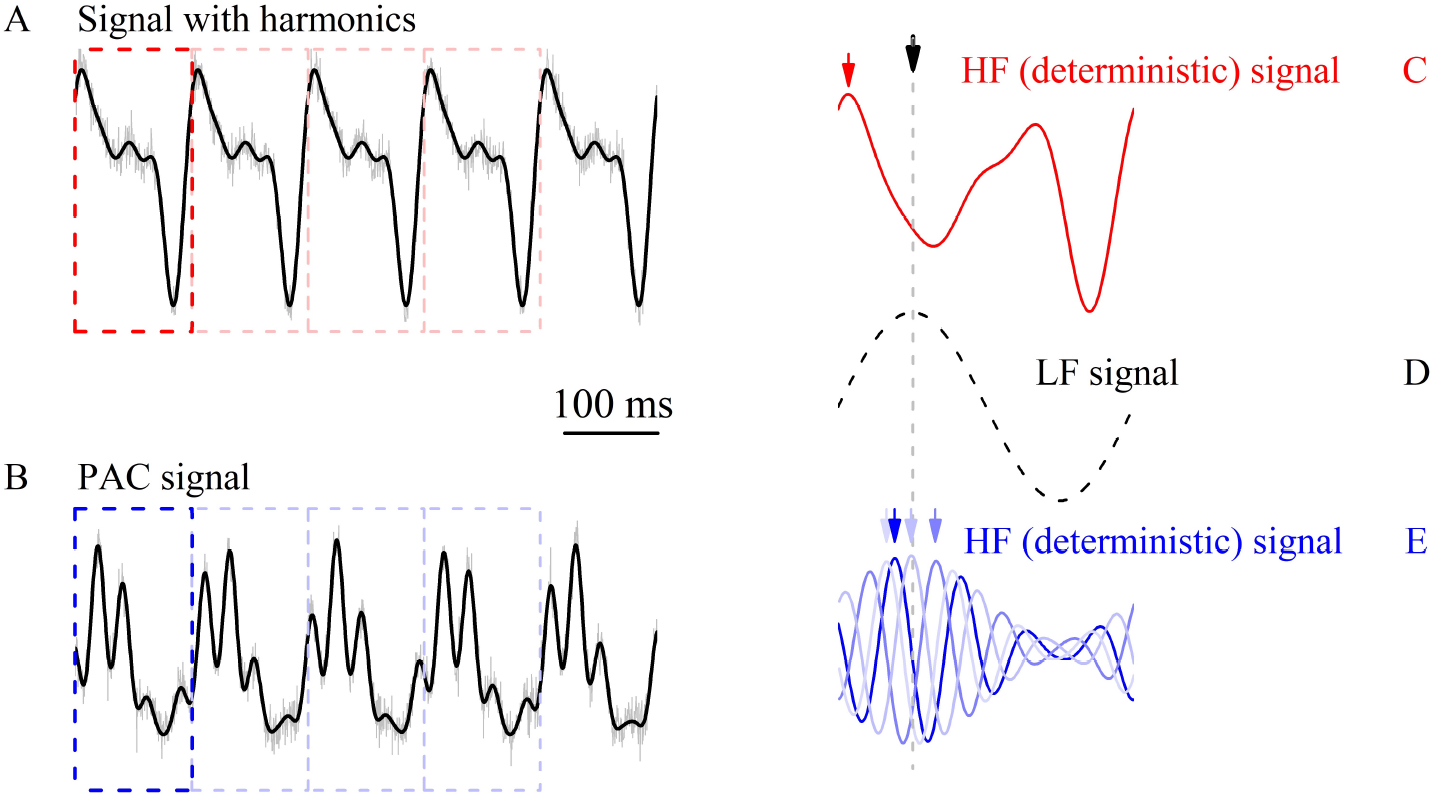
Coupling between low and high frequency signals. (A, C) Upper panels correspond to phase-amplitude coupling via harmonic content: A periodical nonsinusoidal signal, composed by a fundamental sinusoid at 8 Hz plus the first three harmonics with decreasing power, is stereotypically repeated over time (black line in panel A). By chopping the signal in consecutive segments (red boxes, one prototypical in dark red, others in light red), whose length is the period of the fundamental rhythm, the very same signal is obtained (see panel C). In panel C, the true high frequency signal, i.e. the deterministic (noiseless) signal minus the fundamental oscillatory component (thus avoiding filtering artifacts), is shown as well as the arrow corresponding to each maxima in the chopped high frequency signals (red arrow). These maxima always lie in the very same position compared to the low frequency maxima (black arrow). (D) For comparison purposes, the deterministic fundamental low frequency signal can be observed in panel D. (B, E) Lower panels correspond to a phase-amplitude modulated signal: The phase of a fundamental sinusoid at 8 Hz modulates the intensity of a high frequency signal at 65 Hz. Here, high frequency signals corresponding to chopped segments (blue boxes) does not result in a single trace (see panel E). Since the modulation is developed only through amplitude, low and high frequency signals are not tightly coupled regarding phase relationships, and each maxima of the high frequency chopped signal (blue arrows) has a distribution (over phases, or relative time) with respect to the low frequency maxima (black arrow). A small level of additive white Gaussian noise (see noisy signals in gray in panels A and B) does not change conclusions and a concomitant dispersion in the location of high frequency maxima may be observed.

To examine how the TLI metric distinguishes the harmonic CFC from the non harmonic CFC patterns, we can focus on Figure 3. Figures 3A1 and 3A2 show two synthetic signals which are constituted by two coupled oscillatory dynamics plus a small level of extrinsic additive white Gaussian noise (AWGN). Figure 3A1 shows that the amplitude of the fast oscillation *x_HF_* (*t*) is modulated by the phase of the slow rhythm *x_LF_* (*t*) and these two oscillatory dynamics are also phase-locked, as evidenced by the superposition of the individual LF cycles shown at the top of the raw time series *x*(*t*). A similar phase-amplitude coupling is observed between the slow and fast oscillations constituting the signal shown in Figure 3A2, however, the superposition of the individual LF cycles shows no evidence of phase-locked between the slow and fast rhythms in this case. To differentiate these scenarios in a quantitative manner we introduce the TLI metric which exploits the fact that, for a repetitive pattern with a fixed waveform in each cycle, harmonically related frequency bands are intrinsically linked to phase locking oscillations in time domain. The computation of the TLI metric is illustrated in Figures 3B1 and 3B2 for the synthetic signals shown in Figures 3A1 and 3A2, respectively.

**Figure 3:**
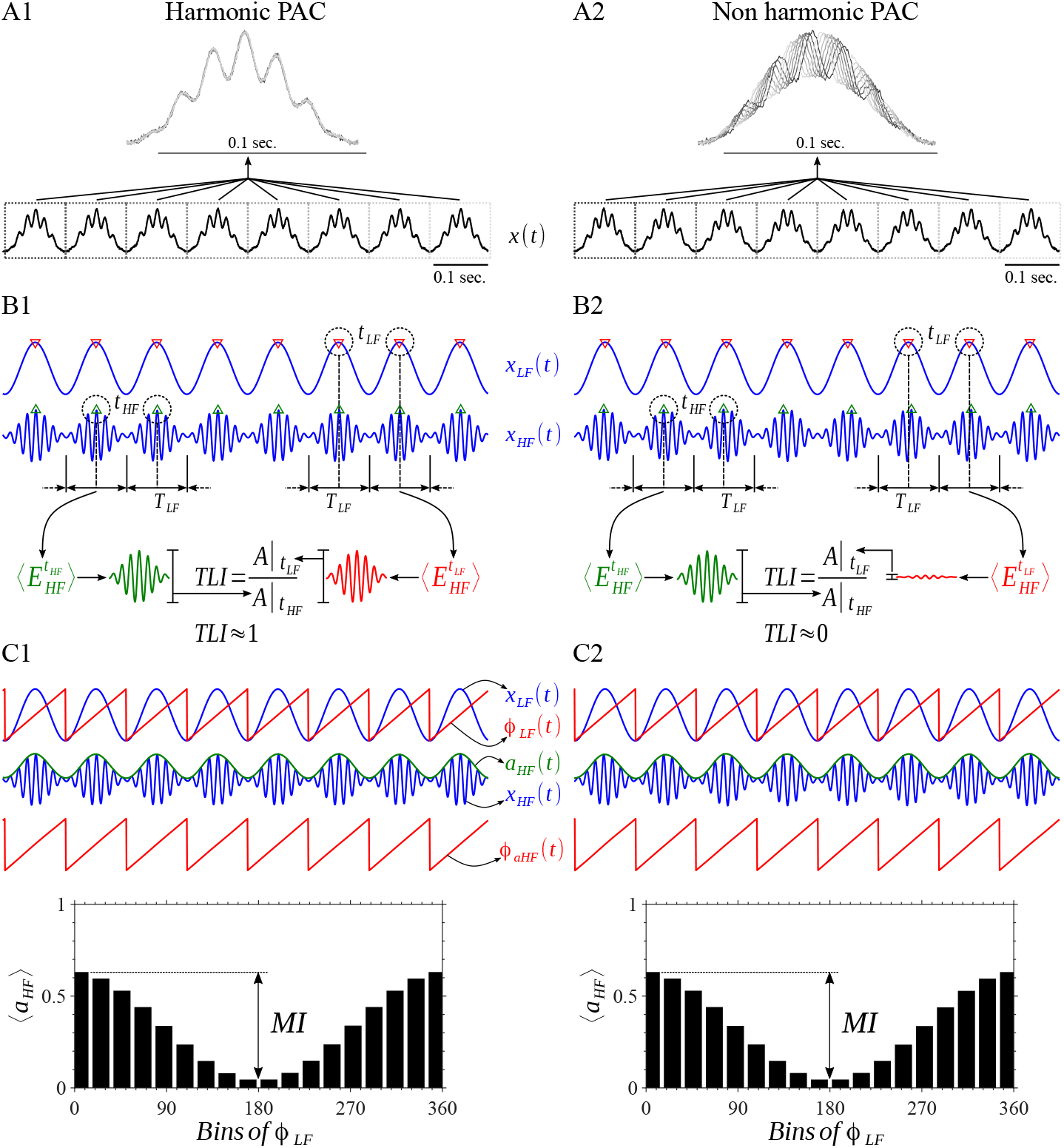
Synthetic amplitude-modulated signals and derived time series involved in the algorithms for quantification of PAC and harmonicity. Amplitude-modulated signals were computed as described in Section Appendix A.1 using a sinusoidal modulating at *f_LF_*= 9 Hz and modulated oscillations at *f_HF_* = 7 × *f_LF_* = 63 Hz and *f_HF_* = 7.1×*f_LF_* = 63.9 Hz for the harmonic PAC and non harmonic PAC, respectively. The LF signals (*x_LF_(t)*) were obtained by filtering the raw signal *_x(t)_* using a band-pass filter centered at 9 Hz and a null-to-null bandwidth of 9 Hz. The HF signals *(x_HF_ (t))* were obtained by filtering the raw signal *x(t)* using a band-pass filter centered at 63 Hz and a null-to-null bandwidth of 81 Hz (see Section Appendix A.5). (A1, A2) Synthetic amplitude-modulated signals. (B1, B2) Time series used to compute the TLI metric to quantify spectral harmonicity. Note that the synthetic harmonic and non harmonic PAC patterns are characterized by *TLI* ≈ 1 and *TLI* ≈ 0 values, respectively. (C1, C2) Time series used to compute the PLV and KLMI metrics to quantify PAC. The histograms show the MI from which the KLMI can be computed. The histograms show the distribution of amplitude of HF as a function of the phase of LF.

In the case of the signal constituted by time-locked oscillations *x*_LF_ and *x_HF_* (Figure 3B1), we obtain similar amplitudes for the time-locked averages 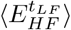 and 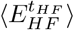 resulting in TLI ≈ 1. On the other hand, in the case of non time-locked oscillations *x_LF_* and *x_HF_*(Figure 3B2), 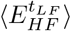 averages out resulting in *TLI* ≈ 0. Importantly, these two essentially different scenarios in terms of spectral harmonicity both present the same level of PAC as evidenced by the phase-amplitude histograms shown in Figures 3C1 and 3C2.

### 2.5. Hilbert-Filter method for instantaneous frequency estimation

Frequency-modulated patterns like phase-frequency (PFC), amplitude-frequency (AFC) and frequency-frequency (FFC) have been the least explored CFC types in neuroscience and biophysics in general, with the remarkable exception of the respiratory sinus arrhythmia associated to the PFC between the respiratory and cardiac rhythms. A possible reason for this may lie in the fact that detection methods to assess frequency-modulated patterns have been poorly described in the specialized literature [26]. Importantly, the conventional CFC metrics (PLV, MVL, KLMI from Eqs. 4 to 7) in combination with equations A.22, A.23 and A.24 constitute a complete formulation to effectively assess PFC, AFC and FFC patterns, provided that a method to compute the instantaneous frequency is given. In this section we briefly discuss the conventional method used to estimate the instantaneous frequency in time and frequency domains [27, 28]. In addition, we provide an alternative approach based on the Hilbert-Filter transformation of the phase time series. The proposed Hilbert-Filter transformation which operates on phase time series to produce an instantaneous frequency time series, should not be confused with the traditional Filter-Hilbert method which operates on raw time series to compute instantaneous phase and amplitude envelopes time series (see Chapter 14, p. 175 in [17]).

Let *ϕ_f_ (τ)* be an *unwrapped* phase time series, corresponding to the band-limited signal *x_f_(τ): f ∈ {LF, HF}*, not constrained to its principal value in the interval (*-π, π*] or [0, 2π), i.e. *ϕ_f_ (τ)* is a continuous function of argument τ ∈ℝ. We also consider that the *ϕ_f_ (τ)* time series has been detrended through a linear fit to remove a trendline with slope 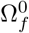. Then, the instantaneous frequency *Ω_f_ (τ)* of the undetrended phase time series is defined as follows [27],

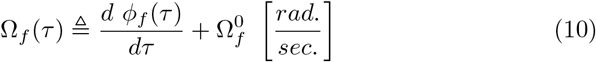

Worthy to note, Eq. 10 implies that a bounded frequency *Ω_f_(τ)* requires a band-limited phase time series *ϕ_f_ (τ)*. This condition can be imposed by bandpass filtering *ϕf (τ)* to restrict it to a finite frequency band of interest (*f ∈ {LF,HF}*). Inthe discrete time domain (*t ∈ ℤ*), Eq. 10 is usually approximated by a low-pass filtered version of the numerical derivative of the phase time series [28],

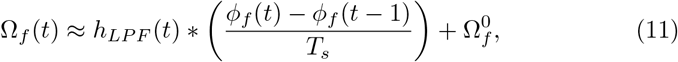

where *T_s_* = 1/*f_s_* is the sampling time interval corresponding to the sampling rate *f_s_*, and * denotes linear convolution. Due to the fact that *ϕ_f_(t)* is not well defined at low amplitude values of the signal *x_f_(t)* (see Eq. A.27), very small or large artifactual values of *ϕ_f_ (t)* sometimes occur which are amplified by the numerical derivative in Eq. 11. To mitigate these artifacts, the numerical derivative is in general smoothed by applying the low-pass filter kernel *h_LPF_ (t)*. Besides, the discrete Fourier transform 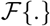 of the difference equation in Eq. 11 can be well described by a first order approximation in the non dimensional angular frequency *ω*, provided that the oversampling condition *(f_s_* ≫ *f : f* ∈ {*LF, HF*}) is satisfied. Under this condition, Eq. 11 can be written as (see Appendix A.6),

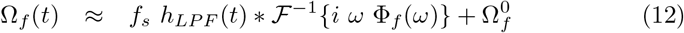

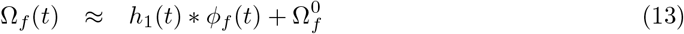

In Eqs. 12 and 13, 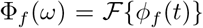 is the discrete Fourier transform of the phase time series, 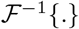 stands for the inverse discrete Fourier transform, *h_1_(t)* is a filter with frequency response equivalent to the cascade connection of the low-pass filter *h_LPF(t)_* and the ideal derivator *iωf_s_*. Importantly, *h_1_(t)* can be implemented as a high-pass or band-pass filter provided that it satisfies two main requirements: within the frequency band of interest (*f ∈ {LF,HF}*), the frequency response of *h_1_(t)* must approximate the magnitude response of the ideal derivator *|ω| f_s_* with the following phase response,

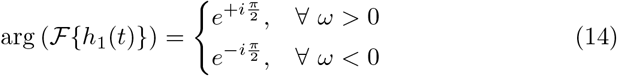

In what follows, we shall obtain an expression equivalent to Eq. 13 by introducing the Hilbert transform with the aim to relax the requirement on the phase response of the filter *h_1_(t)*. The Fourier representation of the Hilbert transformed phase time series is (see Chapter 11, p. 790, Eq. 11.63b in [29]),

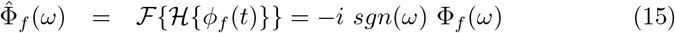

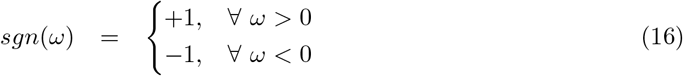

From Eqs. 15 and 16, the Eq. 12 can be written as follows,

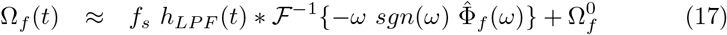

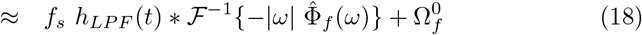

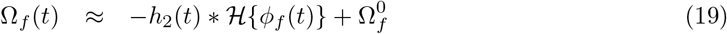

In this case, the frequency response of *h_2_(t)*, within the frequency band of interest *(f ∈ {LF, HF})*, must approximate the magnitude response of the ideal derivator |ω| fs with a zero-phase response (note that the constant phase of π given by the negative sign in equation 19 can be easily introduced as an external gain of −1). As a result, the Hilbert transformed phase time series 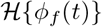 in Eq. 19 accounts for the phase response given by Eq. 14, hence, relaxing this phase requirement on the filter *h_2_(t)*. Besides, for offline data processing applications we can obtain a zero-phase-shift (i.e. non causal) filter by applying a magnitude mask in the frequency domain or by using a linear filter with an arbitrary phase response and reversing the phase delays. In the later case, after filtering the data in the forward direction, the filtered sequence is reversed and passed back through the filter again. Hence, we obtain zero-phase frequency response avoiding phase distortion and delays in the resulting filtered time series [17].

As a conclusion, in the proposed method (Eq. 19), the phase time series ϕ_f_(t) is Hilbert transformed via Eq. A.26 and then passed through a zero-phase (high-pass or band-bass) filter to produce the instantaneous frequency estimation. Note that the order of the Hilbert transformation and filtering indicated in Eq. 19 can be interchanged since they are linear processes. The series of steps involved in the proposed method for the computation of the instantaneous frequency of a frequency-modulated signal (e.g. PFC) can be summarized as follows,

1. The frequency-modulated raw signal *x(t)* is band-pass filtered around the modulated high frequency band (HF) to obtain the band-limited time series *x_HF_(t)*.
2. The Filter-Hilbert method is applied on the signal *x_HF_(t)* to obtain its phase time series (see Eq. A.27 and Chapter 14 in [17]).
3. Unwrap the phase time series.
4. Detrend the phase time series through a linear fit to remove a trendline with slope *Ω^0^_HF_*.
5. The unwrapped and detrended phase time series is band-pass filtered around the modulating low frequency band (*LF*) to obtain the band-limited phase time series *ϕ_HF_(t)*. Note that in the case of a frequency-modulated signal *x(t)*, the oscillatory components of *ϕ_HF_(t)* pertain to the modulating low frequency band *LF*.
6. The Hilbert-Filter method (Eq. 19) is applied on the phase time series *ϕ_HF_(t)* to obtain the instantaneous frequency time series *Ω_HF_(t)*. Note that the zero-phase filter *h_2_(t)* can be of type high-pass or band-pass since the main requirement is that it must approximate the magnitude of the frequency response of the ideal derivator *(|ω| f_s_)* within the modulating low frequency band LF.

Figures 4A and 4D show a frequency-modulated signal *x(t)* (solid black line) and its power spectrum, respectively. The simulated dynamics was obtained from a forced 2nd order parametric oscillator (see Section 3.3). In Figure 4A is possible to distinguish a PFC pattern in which a high frequency oscillation *x_HF_(t)* (*HF*: 10.4-190 Hz, solid red line) is frequency-modulated by the phase of another oscillatory dynamics *x_LF_(t)* with lower frequency (LF: 6.3 - 10.4 Hz, solid green line). The *x_LF_(t)* and xHF(t) time series were obtained bandpass filtering the raw signal *x(t)* (solid black line in Figure 4A) with the filters LF BPF (dotted green line) and HF BPF (dotted red line) shown in Figure 4D, respectively. Figure 4B shows the frequency time series *Ω_HF_*(t) estimated using Eqs. 11 (dotted black line) and 19 (solid black line). Figure 4C shows the phase of the frequency time series required to assess PFC (see Eq. A.22). The low-pass filter *h_LPF(t)_* indicated in Eq. 11 was implemented using a moving average filter with a cutoff frequency (first sidelobe null) equal to the center frequency of the HF BPF (*f_HF_*≈ 90 Hz). The input signal was filtered in forward and reverse direction to obtain zero-phase response (no phase delays). In the time domain, this implies taking the averages over surroundings of 2*f_s_*/*f_HF_* points, where *f*_s_ = 1*/T*_s_ is the sampling rate. Figure 4E shows the frequency response of the moving average filter *h_LPF_*(*t*) (dotted black line). The solid black line in Figures 4E and 4F represents the frequency response of the filter *h*_2_(*t*) used to compute the Eq. 19. Note that the resulting zero-phase band-pass filter (LF BPF) approximate the magnitude response of the ideal derivator in the modulating low frequency band (*LF*: 6.3 - 10.4 Hz, LF BPF has central frequency *f_LF_*≈ 2 Hz and bandwidth *Bw_LF_*≈ 4 Hz). The filter *h*_2_(*t*) was implemented using a Tukey window in the frequency domain (see Appendix A.5).

**Figure 4:**
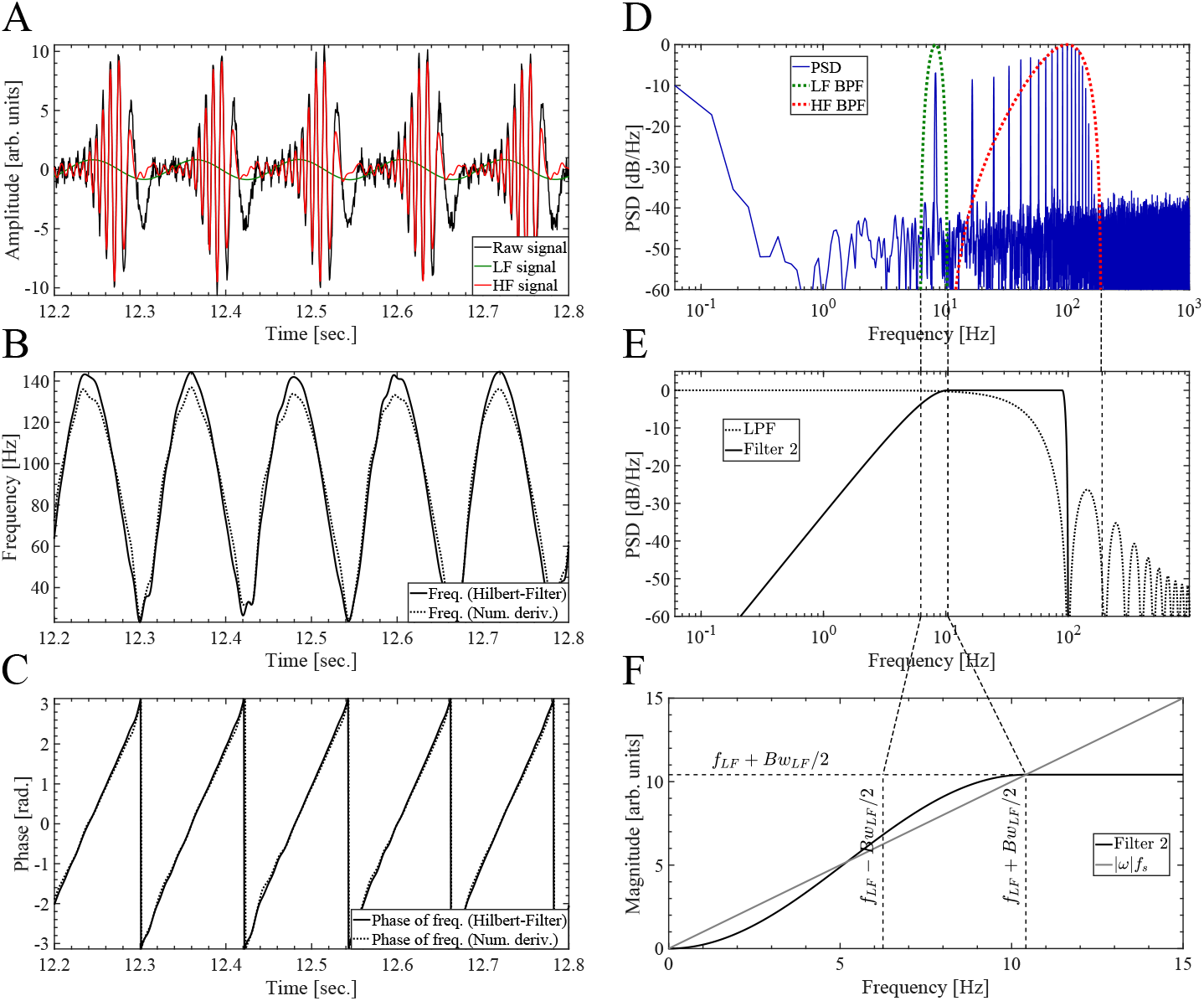
Hilbert-Filter method for instantaneous frequency estimation. (A) Dynamics of the parametric oscillator (solid black line) generated by simultaneously applying an off-resonance external driving *F_e_* and a parametric driving *W_p_* tuned at the same frequency *f_e_* = *f_p_* = *f*_0_/12 ≈ 8.33 Hz and *θ_e_* = 0 (see Eqs. A.16 and A.17 in Appendix A.3). The LF (solid green line) and HF (solid red line) signals where obtained band-pass filtering the raw signal (solid black line) using the BPF whose power responses (i.e. square magnitude) are shown in graph D as dotted green and red lines, respectively. The configuration for the parametric oscillator used in this plots is identical to that used in graphs D and E of Figure 17. (B) Instantaneous frequency time series computed for the raw signal (solid black line) shown in graph A. Solid and dotted black lines correspond to the instantaneous frequency computed using the Hilbert-Filter methos (Eq. 19) and the numerical derivative (Eq. 11), respectively. (C) Instantaneous phase of the frequency time series shown in graph B, computed via Hilbert transformation. (D) Power spectrum (solid blue line) of the dynamics of the parametric oscillator (solid black line in graph A). The power responses (i.e. square magnitude) of the BPF used to compute the LF and HF signals are shown as dotted green and red lines, respectively. (E) The solid black line represents the power response (i.e. square magnitude) of the band-pass filter h2 (t) used to compute the instantaneous frequency time series by means of the Hilbert-Filter method (Eq. 19). The band-pass filter h2 (t) was implemented as described in Appendix A.5 using a Tukey window in the frequency domain. The dotted black line represents the power response of the low-pass filter *h_LPF_(t)* used to compute the instantaneous frequency time series by means of the numerical derivative (Eq. 11). (F) Magnitude responses of the band-pass filter *h_2_(t)* and the ideal derivator *|ω|f_s_*. Regarding the oversampling condition discussed in Appendix A.6, in this case the oversampling ratio is *OSR = f_s_ /f_LF_* = 2000/8.33 ≈ 240.

### 2.6. Harmonicity-CFC plots

To characterize in a quantitative manner the harmonic content of CFC patterns emerging from the oscillatory dynamics explored in this work, we compute harmonicity vs. CFC plots aimed to identify correlations between these two metrics. The harmonicity (TLI) and CFC (PLV, MVL, KLMI) metrics were computed from epochs of 5 sec. or 10 sec. in length corresponding to the synthesized or simulated oscillatory dynamics (see Section 2.1) obtained for a subset of values of a parameter of interest (e.g. modulation depth, amplitude of the external driving, non linear parameter of the oscillator). In the case of simulated data, epochs of twice the required length were computed and then the first half of the time series were discarded to remove the transient period of the numerical simulation. In all the harmonicity-CFC plots shown in this work, the analyzed epochs include between 15 and 90 cycles of the slowest oscillation present in the synthetic or simulated dynamics. We verified that these results hold even in the case of using shorter epoch lengths of ≈ 7 cycles of the slowest oscillation present in the synthetic or simulated dynamics. Then, the scatter plot between the harmonicity and CFC metrics was constructed, in which each data point corresponds to a given value of the parameter of interest. The frequency bands used to compute the harmonicity and CFC metrics were configured accordingly to the time scales of each analyzed oscillatory dynamics.

### 2.7. Comodulograms and harmonicity maps

Comodulograms for the CFC metrics (PLV, MVL, KLMI) were computed following [2, 30] and using the band-pass filters described in Appendix A.5. In all the comodulograms and harmonicity maps show in this work, the analyzed epochs include approx. 60 cycles of the slowest oscillation present in the synthetic or simulated dynamics. Each harmonicity map was constructed by computing the TLI metric for the same modulating (comodulogram *x* axis) and modulated (comodulogram *y* axis) frequency band combinations used to construct the corresponding CFC comodulogram. To assess the statistical significance of the CFC comodulograms, we compute a distribution of 1 ×10^3^ surrogate CFC values achieved by applying the CFC measure (PLV, MVL, KLMI) to sample shuffled *ϕ*_HF_(*t*) or *y*_HF_(*t*) time series (see Eqs. 4 to 7)[6]. Then, assuming a normal distribution of the surrogate CFC values, a significance threshold is then calculated by using P < 0.001 after Bonferroni correction for multiple comparisons [31]. A similar procedure was used to assess the statistical significance of the TLI harmonicity maps using sample shuffled *x_HF_* (*t*) time series (see Section 2.4).

### 2.8. Time series of CFC and harmonicity metrics

Time series were constructed for the TLI, PLV, MVL, KLMI and PC metrics to analyze their temporal evolution during synthetic CFC patterns. The time series were constructed by computing all metrics in a sliding epoch of 20 sec. in length with 90% overlap to include several periods for the slowest modulating rhythms explored. This epoch length was an acceptable trade-off between statistical significance and temporal resolution capable to capture the CFC and harmonicity transients occurring in the synthetic dynamics (see the discussion about Eqs. A.1, A.2 and A.3 in Appendix A.1). Unless otherwise specified, the time series for the CFC metrics were constructed using the Algorithm 2 of Table 2.

**Table 2:** Algorithms to compute the CFC time series.

| Algorithm 1 | Algorithm 2 |
| --- | --- |
| 1. The input time series is Z-score normalized. | 1. The input time series is Z-score normalized. |
| 2. The whole time series is band-pass filtered. | 2. The whole time series is band-pass filtered. |
| **3. The feature (e.g. phase, amplitude) is computed for the whole time series.** | 3. The band-pass filtered time series is then subdivided in sliding epochs. |
| 4. The feature time series is then subdivided in sliding epochs. | **4. Each sliding epoch is Z-score normalized.** |
| 5. The CFC metrics are computed for each sliding epoch. | **5. The feature (e.g. phase, amplitude) is computed for each sliding epoch.** |
|  | 6. The CFC metrics are computed for each sliding epoch. |

## 3. RESULTS

### 3.1. Spectral harmonicity: Characterization of the TLI metric

In this section we discuss the dependence of the TLI on the relevant parameters to quantify the spectral harmonicity in experimental recordings. In addition, we compare the performance of the proposed TLI metric with the conventional method to assess PPC based on the PLV measure (*PLV_PPC_*). For this, we compute the *PLV_PPC_* using Eq. 4 with the configuration given by Eq. A.19. Importantly, both harmonicity metrics are bounded in the range [0,1] which is particularly convenient for the sake of comparison purposes. Due to the fact that to compute the *PLV*_PPC_ measure using Eq. 4 one needs to know a priori the harmonic ratio between the two frequency bands of interest, i.e. the value of *M* and *N*, the characterization presented here is based on synthetic time series in which we have precise control on these parameters. Figure 5 shows, for the case of the linear superposition of two harmonic oscillations (*f*_HF_ */f*_LF_ = 7), the dependence of the TLI and *PLV_PPC_* (*M* = 1, *N* = 7) metrics on the epoch length and the bandwidth of the BPF used used to obtain the fast rhythm (*Bw*_HF_), and taking the noise level as a parameter (AWGN in the range [0%, 200%] of the slow oscillation amplitude). Figure 5C and 5F show the signals and power spectrum for a given set of parameter values, respectively.

**Figure 5:**
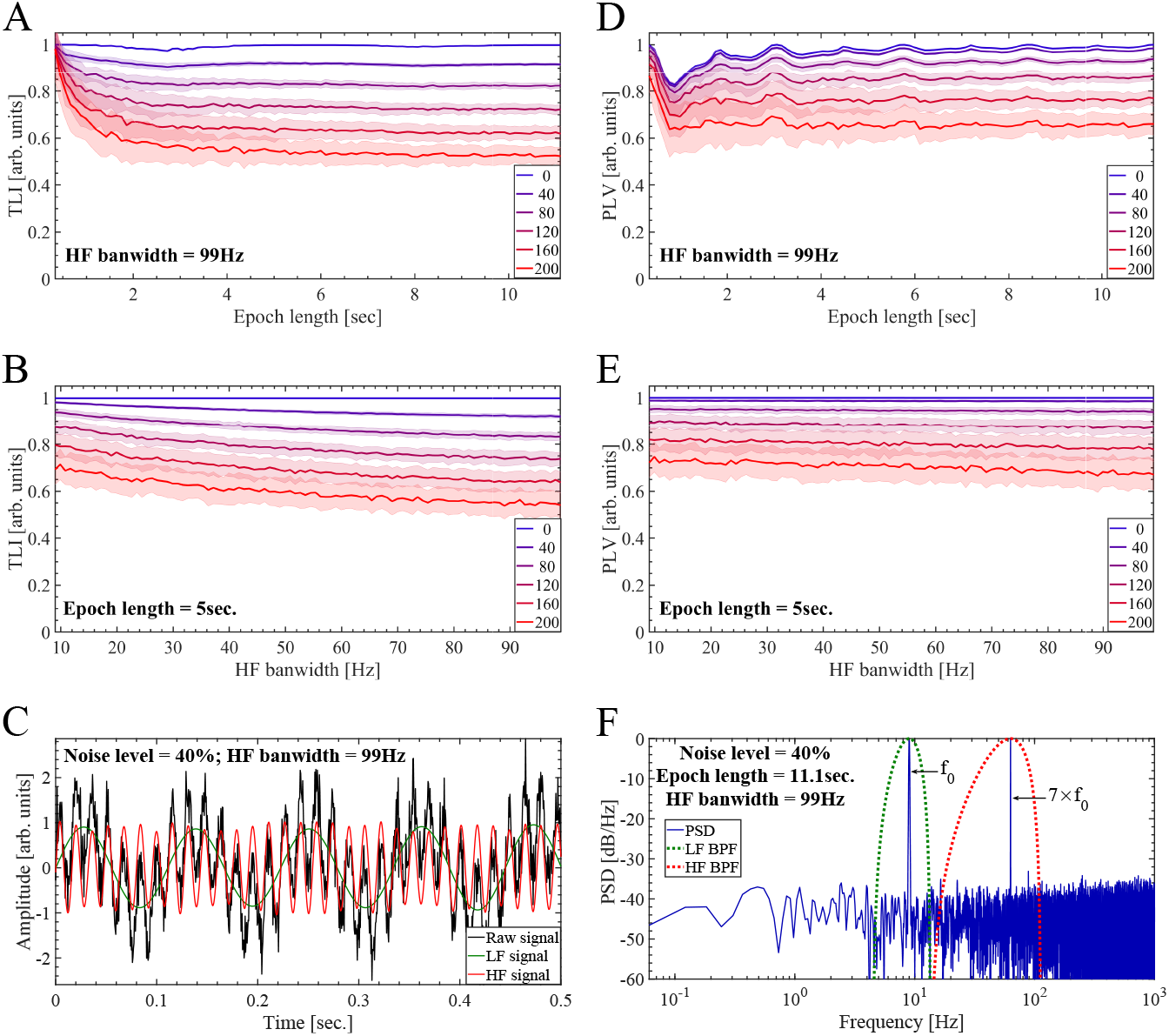
Performance of the TLI and *PLV_PPC_* in quantifying the harmonicity of a synthetic dynamics constituted by the linear superposition of two sinusoidal oscillations at *f_0_* = *f_LF_* = 9 Hz and *f_HF_* = 7 × *f_LF_* = 63 Hz. In all the cases shown in this figure, we used a sampling rate of *f_s_* = 2000 Hz and the frequency and amplitude of the LF and HF oscillations were kept unchanged. To obtain all the band-pass filtered signals shown in this figure we use the BPF as described in Appendix A.5. The bandwidth of the BPF for the LF component (LF BPF) was kept fixed at *Bw_LF_* = 9 Hz. The *PLV_PPC_* was computed using Eq. 4 with the configuration given by Eq. A.19 and *M* = 1, *N* = 7. (A, D) TLI and *PLV_PPC_* metrics as a function of the epoch length and taking the level of additive white Gaussian noise (AWGN) as a parameter. The noise level is expressed as the percent of the amplitude of the LF component at *f_LF_* = 9 Hz scaling the standard deviation σ of the additive white Gaussian noise 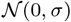. To compute graphs A and D, the bandwidth of the HF BPF was kept unchanged in *Bw_HF_* = 99 Hz. Our implementation of the TLI algorithm (Section 2.4) requires at least 3 cycles of the low frequency oscillation (*f_LF_* = 9 Hz), which determines the minimum epoch length shown in graphs A and D (3/*f_LF_*≈ 0.3 sec.). The maximum epoch length used to compute graphs A and D was 100/*f_LF_* ≈ 11.1 sec. (B, E) TLI and *PLV_PPC_* metrics as a function of the HF bandwidth (*Bw_HF_*) corresponding to the BPF used to obtain the HF signal (*x_HF_(t)*), and taking the level AWGN as a parameter. The minimum and maximum *Bw_HF_* values used to compute the graphs B and E were 9 Hz and 99 Hz, respectively. To compute the graphs B and E, the epoch length was kept unchanged in 45/*f_LF_* ≈ 5 sec. In the panels A, B, D and E, the solid lines represent the mean values and the shaded error bars correspond to the standard deviation of 100 realizations at each point. (C) Synthetic dynamics (solid black line) together with the HF and LF signals shown as solid red and green lines, respectively. The synthetic dynamics includes additive white Gaussian noise 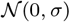 with the standard deviation σ corresponding to the 40% of the amplitude of the LF component at *f_LF_* = 9 Hz. The LF and HF signals where obtained by filtering the raw signal with the band-pass filters whose power responses are shown as dotted green (*Bw_LF_* = 9 Hz) and red (*Bw_HF_* = 99 Hz) lines in graph F, respectively. (F) Power spectrum (solid blue line) of the synthetic dynamics (solid black line in graph C) computed using an epoch length of 100/*f_LF_* ≈ 11.1 sec. The power responses (i.e. square magnitude) of the BPF used to compute the LF and HF signals are shown as dotted green and red lines, respectively.

Our implementation of the TLI algorithm (Section 2.4) requires at least 3 cycles of the low frequency oscillation (*x_LF_*(*t*)), which determines the minimum epoch length used to compute Figure 5 (3*/f_LF_* ≈ 0.33 sec.). As expected, Figures 5B and 5E show that in presence of harmonic oscillations, the value of the harmonicity metrics decay as the AWGN level is increased (*TLI* = 1 without noise and *TLI* ≈ 0.5 for a AWGN level equal to 200% of the slow oscillation amplitude). On the other hand, Figures 5A and 5D show that in case of epoch length including sufficiently large number of slow oscillation cycles, the value of the harmonicity metrics converges to a constant value which depends on the noise level. Importantly, it was found that for short epoch length, comprising less than ≈ 10 cycles of the slow oscillatory component, the TLI and *PLV_PPC_* metrics present a significant bias. This bias produces the high values (≈ 1) of the harmonicity metrics in Figures 5A and 5D for epoch length less than ≈ 1 sec. The bias of the TLI and *PLV_PPC_* metrics was also investigated in presence of non harmonic oscillations. This analysis is discussed in Appendix B.1 and the obtained results support the conclusion drawn from Figures 5A and 5D. Figures 6A and 7A show the *PLV_PPC_* and TLI metrics as a function of the frequency ratio of the two oscillations constituting the synthetic dynamics, and taking the noise level as a parameter. The frequency ratio *f_HF_/f_LF_* was explored for a slow oscillation with *f_LF_* = 3 Hz pertaining the High-Delta band (1-4 Hz) and the fast rhythm with *f_LF_* ranging from the Theta band (4- 8 Hz) to beyond the HFO (High Frequency Oscillations) band (100 - 500 Hz). Note that this cover the conventional frequency bands for the human brain activity which have been defined on the basis of certain cognitive significance and neurobiological mechanisms of brain oscillations [17]. Figures 6A and 7A were computed using epochs of 5 sec. in length and BPF with constant bandwidths ( *Bw*_HF_ = *Bw*_LF_ = 3 Hz), and show that both harmonicity metrics present more dispersion and lower values compared to unity indicating a detriment of their performance for increasing values of AWGN level and frequency ratio between the harmonic oscillations. Besides, we found that the small drop of the TLI metric for high frequency ratios in the case without noise shown in Figure 7A, was due to the effect of the finite sampling rate of the processed time series. In this case the oversampling rate was *f*_s_*/*( 1 80*f_LF_*) = 3.7, where *f*_s_ = 2000 Hz is the sampling rate and 180*f_LF_*= 540 Hz is the maximum frequency explored in Figure 7A. We investigate this finite sampling rate effect on the TLI and *P LV*P P C metrics. In the case of the TLI measure this effect diminished exponentially with the oversampling ratio, producing a drop of the TLI value less than ≈ 5% for oversampling ratios above ≈ 5 (data not shown). The behavior of the *PLV_PPC_* and TLI metrics in between the harmonic frequency ratios is shown in Figures 6B, E, H and 7B, C, D. Figures 6C, F, I and 6D, G, J show the PSD and time series for three representative cases within the explored range of frequency ratios.

**Figure 6:**
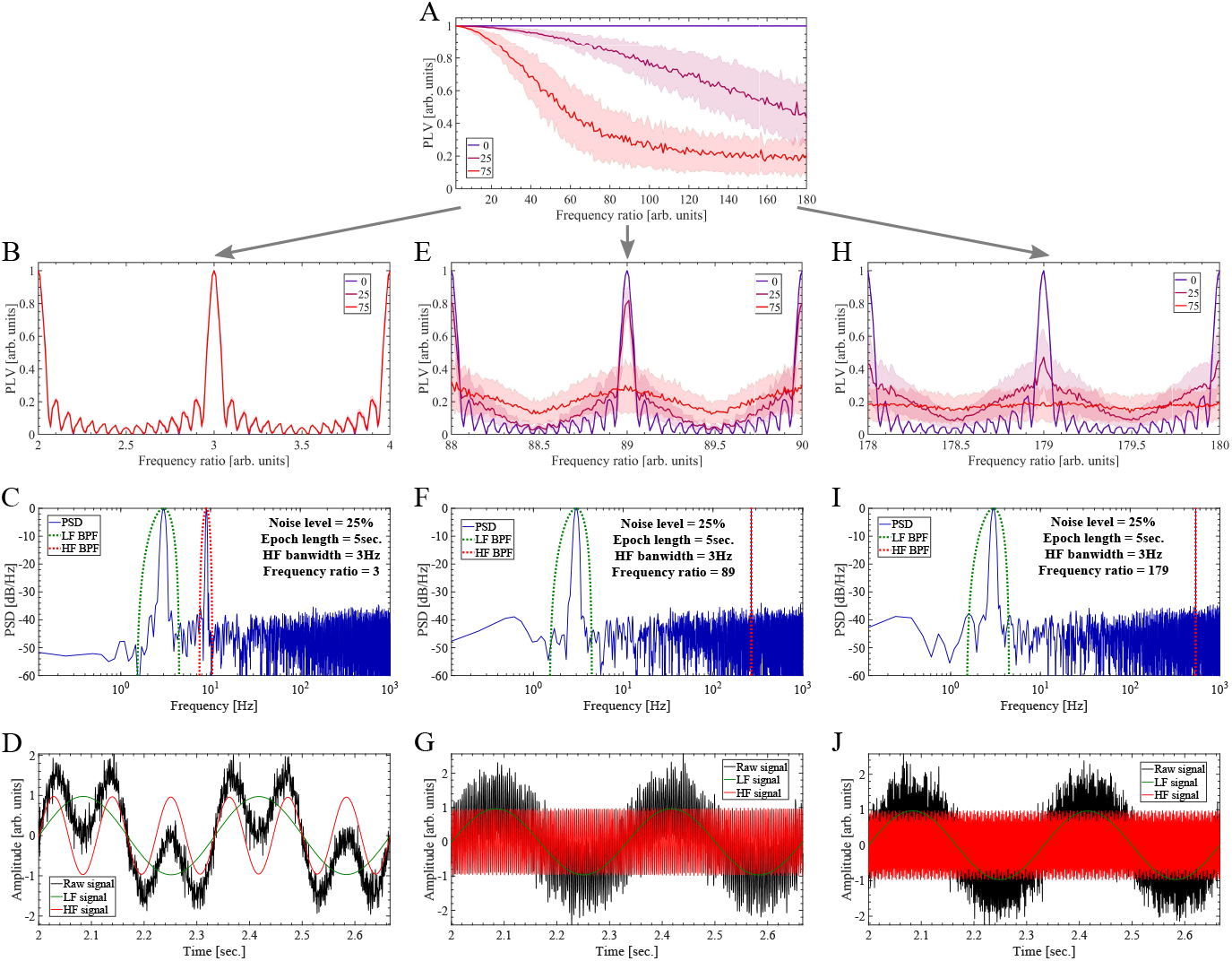
Performance of the *PLV_PPC_* in quantifying the harmonicity of a synthetic dynamics constituted by the linear superposition of two sinusoidal oscillations. In all the cases shown in this figure, we used a sampling rate of *f_s_* = 2000 Hz and the amplitude of the LF and HF oscillations were kept unchanged. To obtain all the band-pass filtered signals shown in this figure we use the BPF as described in Appendix A.5. The bandwidth of the BPF for the LF (LF BPF) and HF (HF BPF) components were kept fixed at *Bw_LF_* = *Bw_HF_* = 3 Hz. The P LV P P C was computed using Eq. 4 with the configuration given by Eq. A.19. (A) P LV P P C intensity as a function of the frequency ratio *f_HF_* /*f_LF_* and taking the level of additive white Gaussian noise (AWGN) as a parameter. The LF component was kept fixed at *f_LF_* = 3 Hz and the frequency of the HF oscillation was varied in the range 2 × *f_LF_* ≤ *f_HF_* ≥ 180 × *f_LF_*. The noise level is expressed as the percent of the amplitude of the LF component at *f_LF_* = 3 Hz scaling the standard deviation σ of the additive white Gaussian noise 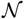(0, σ). The *PLV_PPC_* was computed using an epoch length of 45/*f_LF_* ≈ 5 sec. (B, E, H) Evolution of the *PLV_PPC_* intensity in between the harmonic frequency ratios *f_H F_*/*f_LF_* for three AWGN levels. In the panels A, B, E and H, the solid lines represent the mean values and the shaded error bars correspond to the standard deviation of 100 realizations at each point. (C, F, I) Power spectrum (solid blue line) of the synthetic dynamics (solid black line in graphs D, G and J) corresponding to three frequency ratio values (*f_HF_* /*f_LF_* = 3, 89, 179). The power spectra were computed using an epoch length of 45/*f_LF_* ≈ 5 sec. The power responses (i.e. square magnitude) of the BPF used to compute the LF and HF signals are shown as dotted green and red lines, respectively. (D, G, J) Synthetic dynamics (solid black line) together with the HF and LF signals shown as solid red and green lines, respectively, corresponding to three frequency ratio values (*f_HF_* /*f_LF_* = 3, 89, 179). The synthetic dynamics includes additive white Gaussian noise 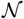(0, σ) with the standard deviation σ corresponding to the 25% of the amplitude of the LF component at *f_LF_* = 9 Hz. The LF and HF signals where obtained by filtering the raw signal with the band-pass filters whose power responses are shown as dotted green (*Bw_LF_* = 3 Hz) and red (*Bw_HF_* = 3 Hz) lines in graphs C, F and I, respectively.

**Figure 7:**
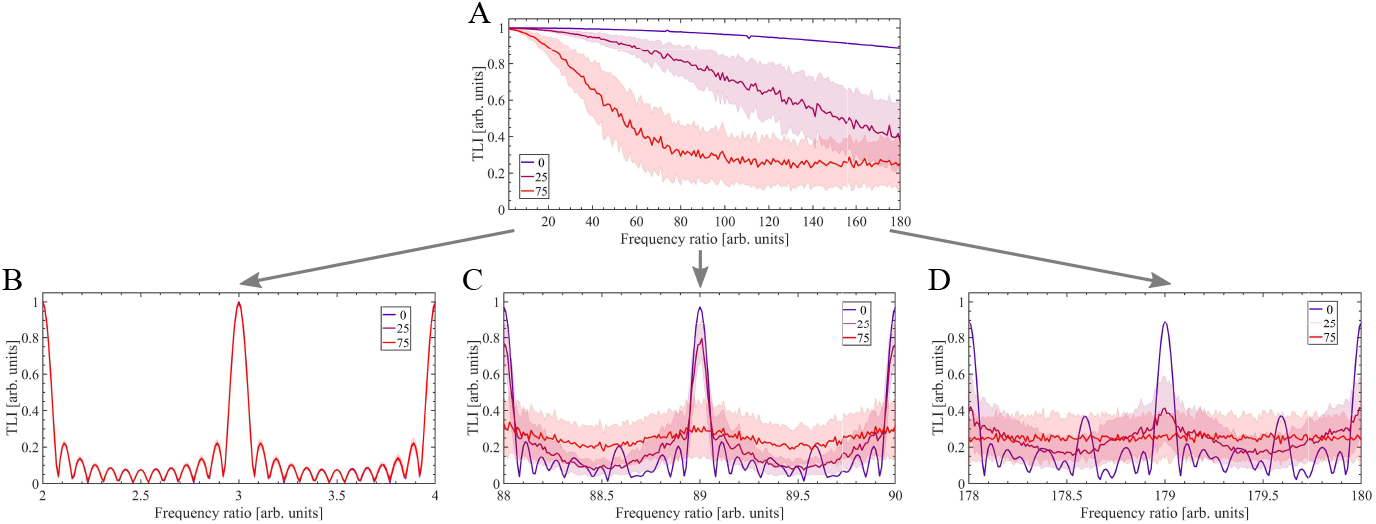
Performance of the TLI metric in quantifying the harmonicity of a synthetic dynamics constituted by the linear superposition of two sinusoidal oscillations. For the sake of comparison purposes this figure was computed using the same hyperparameters than those used to compute and process the synthetic dynamics shown in Figure 6.

Figure 8 show the performance of the harmonicity (TLI and *PLV_PPC_*) and PAC (*KLMI_PAC_*) metrics in a simple multi-harmonic oscillatory dynamics capable to generate PAC. For this, we compute the *KLMI_PAC_* using Eqs. 6 and 7 with the configuration given by Eq. A.20. The dynamics was synthesized using Eqs. A.1, A.4 and A.6 configured for the DSB-C case with a sinusoidal modulating *a*(*t*) and maximum modulation depth (see the caption of Figure 8 for the complete list of parameter values). Figures 8A and 8C for the multi-harmonic dynamics should be compared with their counterparts in the case of a single HF harmonic component shown in Figures 5A and 5D, respectively. While no significant differences are observed in the *PLV_PPC_* metric between these two scenarios (see Figures 8C and 5D), the TLI metric present higher values (close to unity) and less dispersion when multiple harmonics are included in the HF bandwidth (see Figures 8A and 5A). This result is consistent with the behavior observed in Figures 8B and 8D showing an opposite trend between the two harmonicity metrics, that is, as the HF bandwidth increases a concomitant increase in the dispersion and drop of the values occurs in the *PLV_PPC_* metric and the opposite is observed for the TLI measure. Figure 8E shows the PAC metric as a function of the epoch length and taking the AWGN as a parameter. Importantly, Figure 8F shows that the *KLMI_PAC_* metric increases only after the HF bandwidth is wide enough to include the two sidebands (6 × *f_LF_* and 8 × *f_LF_*) around the carrier *(f_HF_* = 7 × *f_LF_*), that is *Bw_HF_*≳2 × *f_LF_* = 18 Hz. Worthy to note, the increase rate of the *KLMI_PAC_* curves in Figure 8F is related to the steepness (i.e. transition-band width) of the BPF used to obtain the fast (amplitude-modulated) rhythm. That is, the steeper the roll-offs of the BPF the higher the increase rate of the *KLMI_PAC_* curves in Figure 8F. Note that we do not use BPF with very steep roll-offs to prevent creating artificial narrow-band oscillations [31, 32] (see Appendix A.5). Figures 8G and 8H show the PSD and time series for a representative case within the explored parameters.

**Figure 8:**
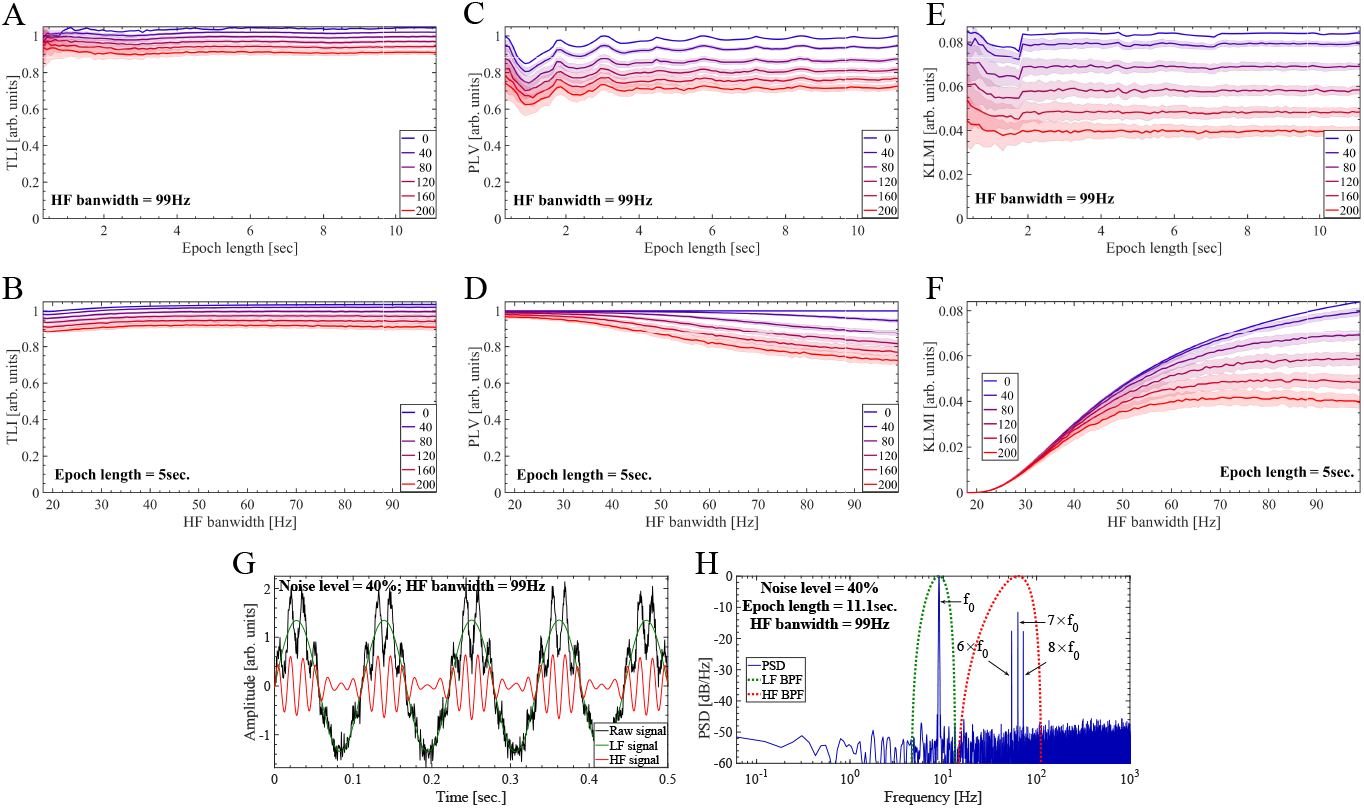
Performance of harmonicity (TLI, *PLV_PPC_*) and PAC (*KLMI_PAC_*) metrics to characterize an amplitude modulated synthetic signal. In all the cases shown in this figure, we used a sampling rate of *f_s_* = 2000 Hz and the frequency and amplitude of the modulating (LF) and the amplitude-modulated (HF) oscillations were kept unchanged. To obtain all the band-pass filtered signals shown in this figure we use the BPF as described in Appendix A.5. The bandwidth of the BPF for the LF component (LF BPF) was kept fixed at *Bw_LF_* = 9 Hz. The *PLV_PPC_* was computed using Eq. 4 with the configuration given by Eq. A.19 and *M* = 1, *N* = 7. The amplitude-modulated signal was synthesized as described in Appendix A.1 using the following hyperpameter values: *c* = 1 (i.e. DSB-C), maximum modulation depth *m* = 0, η_m_ = 0, we used a sinusoidal modulating *a(t)* at *f_0_* = *f_LF_* = 9 Hz as given by Eq. A.6, *A_m_* = 1, ϕ*_m_* = 0, *z_DBS_* was set with *f_H F_* = 7 × *f_LF_*= 63 Hz, ϕ*_c_* = 0, *z_HF_* = 0, for *z*_h_ we use *A*_1_ = 4, *A_k_* = 0 ∀ k > 1 and ϕ_k_ = 0 ∀ *k*. In Eq. A.1, we configured a constant amplitude envelope *ε(t)* = 1. The extrinsic noise level shown in the graphs corresponds to *η(t)* in Eq. A.1, and is expressed as the percent of the modulating signal *a(t)* maximum amplitude (*A_m_*) scaling the standard deviation σ of the additive white Gaussian noise (AWGN) 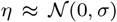. (A, C, E) Harmonicity (TLI, *PLV_PPC_*) and PAC (*KLMI_PAC_*) metrics as a function of the epoch length and taking the level of AWGN as a parameter. To compute graphs A, C and E, the bandwidth of the HF BPF was kept unchanged in *Bw_HF_* = 99 Hz. Our implementation of the TLI algorithm (Section 2.4) requires at least 3 cycles of the low frequency oscillation (*f_LF_* = 9 Hz), which determines the minimum epoch length shown in graphs A and D (3/*f_LF_* ≈ 0.3 sec.). The maximum epoch length used to compute graphs A and D was 100/*f_LF_* ≈ 11.1 sec. (B, D, F) Harmonicity (TLI, *PLV_PPC_*) and PAC (*KLMI_PAC_*) metrics as a function of the HF bandwidth (*Bw_HF_*) corresponding to the BPF used to obtain the HF signal (*x_HF_(t)*), and taking the level AWGN as a parameter. The minimum and maximum *Bw_HF_* values used to compute the graphs B and E were 18 Hz and 99 Hz, respectively. To compute the graphs B, D and F, the epoch length was kept unchanged in 45/*f_LF_* ≈ 5 sec. In the panels A, B, C, D, E and F, the solid lines represent the mean values and the shaded error bars correspond to the standard deviation of 100 realizations at each point. (G) Synthetic dynamics (solid black line) together with the HF and LF signals shown as solid red and green lines, respectively. The synthetic dynamics includes additive white Gaussian noise 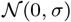 with the standard deviation σ corresponding to the 40% of the modulating signal *a(t)* maximum amplitude (*A_m_*). The LF and HF signals where obtained by filtering the raw signal with the band-pass filters whose power responses are shown as dotted green (*Bw_LF_*= 9 Hz) and red (*Bw_HF_*= 99 Hz) lines in graph H, respectively. (H) Power spectrum (solid blue line) of the synthetic dynamics (solid black line in graph G) computed using an epoch length of 100/*f_LF_* ≈ 11.1 sec. The power responses (i.e. square magnitude) of the BPF used to compute the LF and HF signals are shown as dotted green and red lines, respectively.

As a conclusion, it was found that for dynamics with two harmonic (Figures 5 to 8) or two non harmonic (Figure B.1) oscillatory components, the TLI and *PLV_PPC_* metrics have a comparable performance in terms of the explored parameters. On the other hand, for oscillatory dynamics containing multi-harmonic HF components the TLI present a better performance when compared with the *PLV_PPC_* metric under similar conditions (compare panels A and C of Figure 8). This aspect will be further discussed below in connection with the simulated dynamics of the Van der Pol oscillator.

The TLI metric was tailored designed to be combined with the PC metric for improving the characterization and interpretation of the CFC patterns observed at the signal level. To illustrate this point, the temporal evolution of the relevant metrics were analyzed during a variety of synthetic oscillatory dynamics. For this, time series for the PLV, KLMI, TLI and PC metrics were constructed as it was described in Section 2.8. It is essential to note a key point regarding the TLI temporal evolution as a complementary tool to interpret the estimators aimed to quantify CFC (e.g. PLV, MVL, KLMI). Even though the TLI is a measure bounded in the range [0, 1] (see Section 2.4) and independent of the processed oscillations amplitude, the absolute value of the TLI does depend on the noise level present in the processed time series and on the epoch length, i.e. the number of periods of the low frequency oscillation taken to implement the time-locked average involved in the TLI computation (see Figures 5 to 8 and B.1). A similar behavior was observed for the bounded (PLV, KLMI ∈[0, 1]) and unbounded (MVL) CFC metrics. As a consequence, a robust indicator of the occurrence of transient harmonic CFC patterns is given by the fact that the TLI increases concurrently with the CFC metrics, rather than by the absolute TLI value at a particular time instant. In this regard, Figure 9A shows a synthetic dynamics presenting a transient harmonic PAC pattern. This type of transient dynamics is relevant since it is commonly observed during the ictal activity recorded invasively in patients candidates to epilepsy surgery and animal models of epilepsy (see [11] and references therein). The dynamics was synthesized using Eqs. A.1, A.4 configured for the DSB-C case with a Gaussian modulating *a(t)* (Eqs. A.7 and A.8) and maximum modulation depth (see the caption of Figure 9 for the complete list of parameter values). The transient harmonic PAC pattern was implemented through the time series envelope *E(t)* as defined in Eqs. A.2 and A.3. Figure 9B shows that the PAC (P LVPAC) and harmonicity (TLI) metrics increase almost concurrently from their baseline value previous to the transient activation to close the unity. Note that while the amplitude-modulated dynamics remains stable (80sec. ≲ *Time* ≲ 120sec. in Figure 9A) so the PAC and harmonicity metrics indicating the occurrence of a PAC pattern (*PLV_PAC_* ≈ 1) constituted by harmonic spectral components (*TLI* ≈ 1) which is not an ephiphenomenon due to the presence of phase clustering (*PC_LF_* ≈ 0). Figures 9D and 9E show the signals and power spectrum representative of this time interval in which the dynamics remains stable. In particular, 9D shows the modulating signal with *f_LF_*= 3 Hz (solid green line, *f_LF_*= 3 Hz) and modulated (*f*_HF_= 89 × *f_LF_*= 267 Hz, red solid line) signals obtained band-pass filtering the raw dynamics (solid black line). The modulating and modulated signals were computed using the LF BPF (dotted green line) and HF BPF (dotted red line) shown in Figure 9E, respectively. In addition, the harmonicity map and comodulogram computed for an epoch during the time interval in which the dynamics remains stable are shown in Figures 9C and 9F, respectively, revealing the modulating (*f_LF_*) and modulated (*f_HF_*) frequency bands involved in the harmonic PAC pattern.

**Figure 9:**
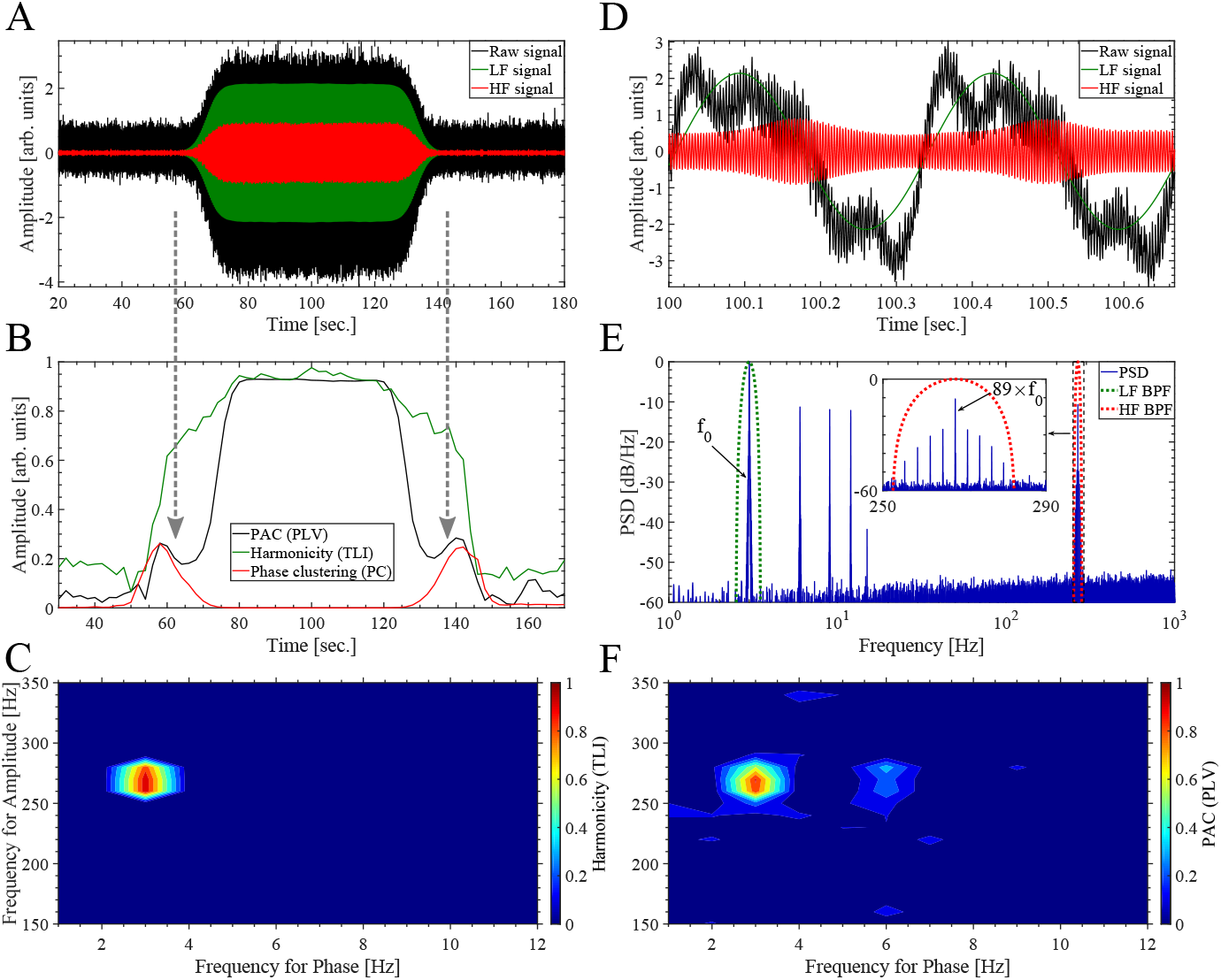
Temporal evolution of the PAC (*PLV_PAC_*), harmonicity (TLI) and phase clustering (*PC_LF_*) metrics during a synthetic dynamics presenting a transient harmonic PAC pattern. To obtain all the band-pass filtered signals shown in this figure we use the BPF as described in Appendix A.5. (A) Synthetic dynamics (solid black line) together with the HF and LF signals shown as solid red and green lines, respectively. The dynamics (solid black line) was synthesized using Eqs. A.1 and A.4 with the following hyperpameter values: sampling rate *f_s_* = 2000 Hz, c = 1 (i.e. DSB-C), half modulation depth *m* = 0.5, η_*m*_ = 0, we used a Gaussian modulating *a(t)* with the fundamental frequency at *f_0_* = *f_LF_*= 3 Hz as given by Eqs. A.7 and A.8 with σ ≈ 55 and *A_m_* = 1, *z_DBS_* was set with *f_HF_* = 89 × *f_LF_* = 267 Hz, ϕc = 0, z_H F_= 0, for z_h_ we use A1 = 4, A_k_= 1 ∀2 ≤ k ≤ 4, A_k_= 0 ∀k ≥ 5 and ϕ_k_= 0 ∀k. The transient harmonic PAC pattern was implemented through the time series envelope *E(t)* as defined in Eqs. A.2 and A.3, with α = 0.5 and β equals to one third of the time series length. Extrinsic noise η (t) was added as shown in Eq. A.1. In this case the noise level corresponds to the 10 percent of the maximum amplitude of the deterministic part of signal *x(t)* (i.e first term of the right-hand member of the Eq. A.1), scaling the standard deviation σ of the additive white Gaussian noise (AWGN) 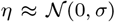. The LF (solid green line) and HF (solid red line) signals where obtained by filtering the raw signal (solid black line) with the band-pass filters whose power responses are shown as dotted green (*Bw_LF_*= 1 Hz) and red (*Bw_HF_*= 30 Hz) lines in graph E, respectively. (B) Time series showing the temporal evolution of the *PLV_PAC_*, TLI and *PC_LF_* metrics. These time series were computed as described in Section 2.8 using the algorithm 2 summarized in Table 2, with a sliding window of 20 sec. in length, i.e. 60 cycles of the slowest oscillatory component at *f_0_* = *f_LF_* = 3 Hz. (C) TLI harmonicity map computed as described in Section 2.7 using a 20 sec. epoch extracted from the center (*Time* ≈ 100 sec.) of the synthetic dynamics shown in panel A. In computing the map, all the TLI values below the significance threshold were set to zero (see Section 2.7). The pseudocolor scale represents the TLI values ranging from 0 (blue) to 1 (red). (D) Zoom showing two cycles of the synthetic dynamics (solid black line) together with the HF and LF signals shown as solid red and green lines, respectively. The two cycle epoch corresponds to the center (T ime ≈ 100 sec.) of the synthetic dynamics shown in panel A. (E) Power spectrum (solid blue line) computed from the synthetic dynamics (solid black line in graph A). The power responses (i.e. square magnitude) of the BPF used to compute the LF and HF signals are shown as dotted green and red lines, respectively. (F) Comodulogram computed as described in Section 2.7 computed from the same epoch used to obtain the harmonicity map (panel C). In computing the comodulogram, all the |*PLV_PAC_*| values below the significance threshold were set to zero (see Section 2.7). The pseudocolor scale represents the |*PLV_PAC_*| values ranging from 0 (blue) to 1 (red). The harmonicity map (panel C) and comodulogram (panel F) were computed using the same BPF (see Appendix A.5).

Due to the fact that to compute the metrics shown in Figure 9 we used sufficiently narrow LF BPF to obtain an almost sinusoidal low frequency component (dotted green line in Figure 9E and solid green line in Figure 9D), i.e. uniform distribution of ϕLF (t) values in Eq. 8, the |*PC_LF_* | time series is close to zero along the transient dynamics (red solid line in Figure 9B). This indicates that the observed PAC pattern is not a spurious artifacts related to the presence of phase clustering in the modulating LF component [17, 18]. On the other hand, Figure 10 corresponds to the very same synthetic dynamics of that shown in Figure 9A, but in this case we use a wide LF BPF including several spectral components, and thus, resulting in a highly non sinusoidal low frequency component (dotted green line in Figure 10E and solid green line in Figure 10D). As a consequence, we obtain a skewed distribution of phase angles producing |PC_LF_| ≈ 0.5. It is crucial to note that this finite phase clustering associated to a non sinusoidal low frequency component (*PC_LF_*) produces a bias in both the PAC and harmonicity metrics, which in this case becomes evident by comparing Figures 9B and 10B. Note that a wider LF BPF (dotted green line in Figure 10E) imposes a limit on the minimum value of the frequency for phase (abscissa) that is possible to compute in the harmonicity map and comodulogram as it is shown in Figures 10C and 10F, respectively.

**Figure 10:**
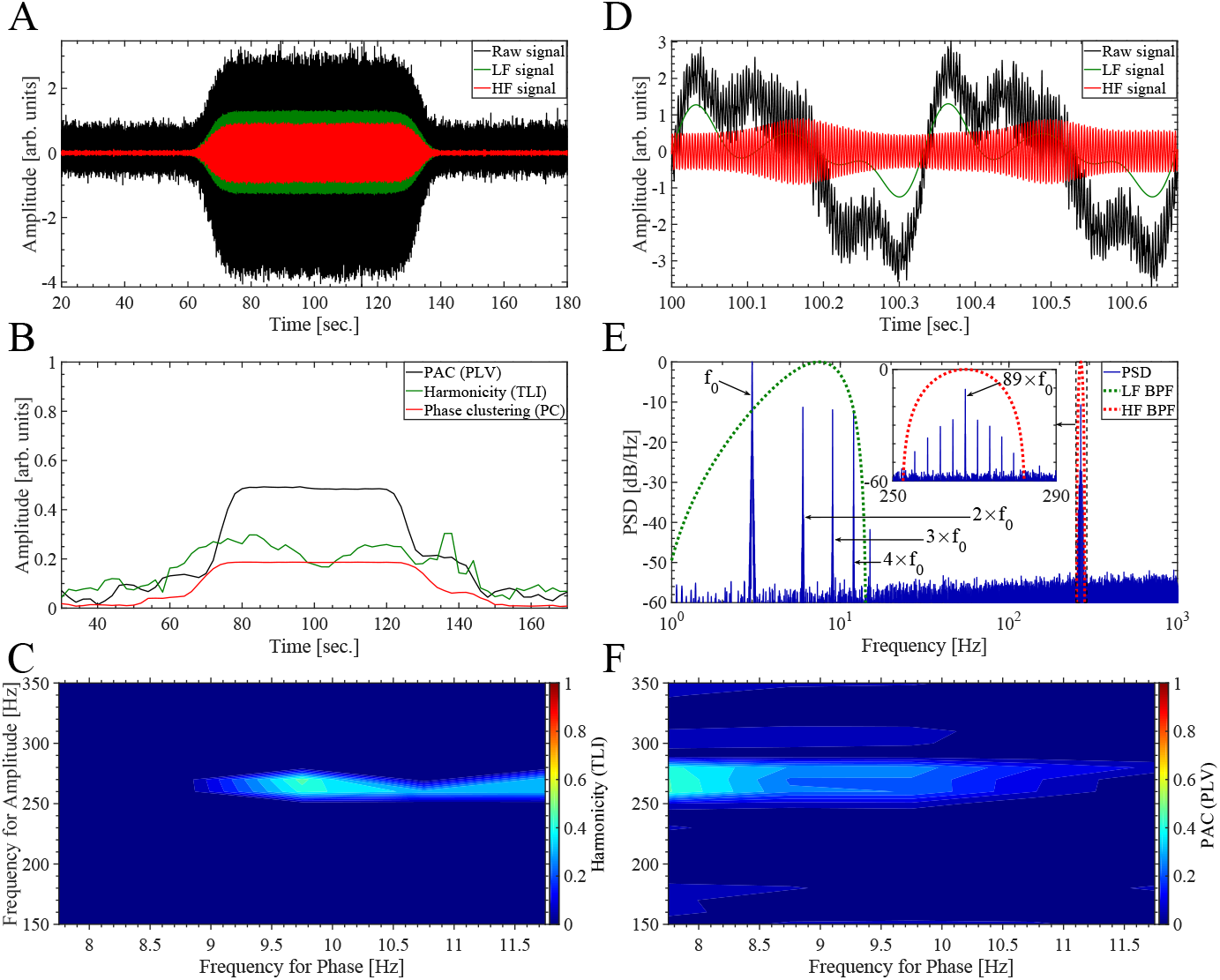
Temporal evolution of the PAC (*PLV_PAC_*), harmonicity (TLI) and phase clustering (*PC_LF_*) metrics during a synthetic dynamics presenting a transient harmonic PAC pattern. In this plot we use the same synthetic dynamics and the same set of hyperparameter values to compute the metrics than those described in the caption of Figure 9, except for the bandwidth of the BPF used to compute the LF component (Bw_LF_). In this case, the *PLV_PAC_*, TLI and *PC_LF_* metrics were computed using *Bw_LF_* = 13.5 Hz centered around 7.5 Hz (see the dotted green line in panel E). This wide BPF produces a non sinusoidal LF component (see solid green line in panel D), characterized by a non uniform distribution of phase values producing the increase of the phase clustering (*PC_LF_*) during the dynamics (see solid red line in panel B). Note the bias in the PAC (*PLV_PAC_*) and harmonicity (TLI) metrics due to the presence of phase clustering (*PC_LF_*). The description of the panels is the same than that given in Figure 9.

In Appendix B.2 we present the behavior of the harmonicity and PAC metrics during a variety of transient dynamics (e.g. non harmonic PAC), and also discuss the bias produced on the MVL metric (*MVL_PAC_*) by the phase clustering associated to a non sinusoidal low frequency component (*PC_LF_*).

It was found that the abrupt change of the raw signal amplitude associated to transient dynamics like those shown in Figures 9, 10, B.2 to B.5, is capable to produce spurious CFC values due to the interaction between the sliding epoch and the abrupt change of the amplitude envelope of the raw time series (see grey arrows in panels A and B of Figure 9). Importantly, these spurious CFC values at rising and falling edges of the transient dynamics are effectively detected by the *PC_LF_* metrics since they occurs concomitantly with an increase of the phase clustering. On the other hand, due to the fact that the TLI is an amplitude independent quantity sensitive only to PPC, it does not present these artifacts associated to changes in the amplitude of the analyzed dynamics (see the TLI time series in Figures 9, 10, B.2 to B.5).

We identify another confounding associated to the Algorithm 1 described in Table 2 for the computation of CFC time series. Specifically, Figures 11A and 11C show the P LVPAC and PCLF time series computed with the algorithms described in Table 2 together with the TLI metric for a transient harmonic PAC pattern similar to that shown in Figure 9. We found that using short sliding epochs of about 10 cycles of the slowest oscillation, Algorithm 1 produce time series of PAC metrics (e.g. *PLV_PAC_*) which are monotonically decreasing toward and away from the rising and falling edges of the transient dynamics (see grey arrows in Figure 11A). Besides, Figure 11B shows that this effect is also distinguishable in the case of a transient oscillatory dynamics without PAC similar to that shown in Figure B.2. We observed this behavior of the CFC time series computed via the Algorithm 1 in a variety of transient oscillatory dynamics, suggesting that is a confounding strongly related to the abrupt change of the amplitude envelope of the raw time series, irrespective of the type and intensity of the CFC present in the dynamics. We emphasize that this confounding is particularly dangerous since it seems not to be associated to an concomitantly increase of the phase clustering time series, and therefore it is difficult to detected (see red solid line in Figures 11A and 11B). On the other hand, Figures 11C and 11D show that the monotonically decreasing trend is absent in the time series of PAC metrics computed using the Algorithm 2. Moreover, when comparing Figures 11A and 11C it becomes evident the bias introduced by the Algorithm 1 in the maximum intensity of the P LVPAC time series. We identify the root cause of these confounding as the computation of features (e.g. phase, amplitude, frequency) via the Hilbert transform on the whole band-pass filtered time series including the abrupt changes of amplitude associated to the rising and falling edges of the transient dynamics, which affect the resulting features (see step 3 in Algorithm 1 of Table 2). On the other hand, these issues are avoided in the Algorithm 2 of Table 2 by first dividing the band-pass filtered signals in sliding epochs, the resulting epochs are Z-scored to make them independent of the absolute amplitude of the filtered signals and then the features are computed by applying the Hilbert transform on the Z-scored epochs (see step 4 and 5 in Algorithm 2 of Table 2).

**Figure 11:**
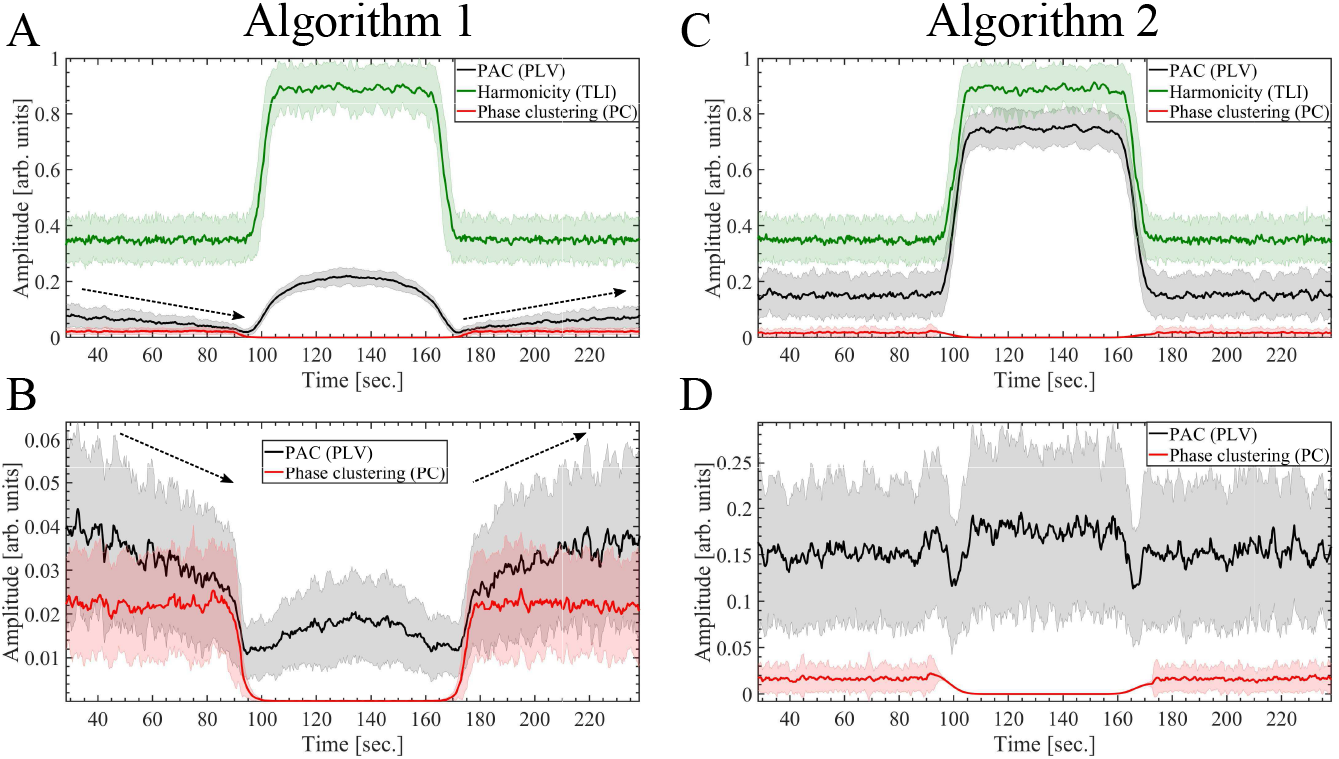
Comparison of the algorithms 1 and 2 are summarized in Table 2 aimed to compute time series of CFC metrics. The time series showing the temporal evolution of the *PLV_PAC_*, TLI and PCLF metrics were computed as described in Section 2.8 using a sliding window of 3.33 sec. in length, i.e. 10 cycles of the slowest oscillatory component at *f_0_* = *f_LF_* = 3 Hz. (A, C) The time series shown in panels A and C were computed from a synthetic dynamics presenting a transient harmonic PAC pattern, as described in Section 2.8 using the algorithms 1 and 2, respectively. The synthetic dynamics used in panels A and C was computed using the same set of hyperparameter values as those described in the caption of Figure 9, with the exception of the extrinsic noise *η(t)* which in this case was set to 30 percent of the maximum amplitude of the deterministic part of signal *x(t)* (i.e first term of the right-hand member of the Eq. A.1). (B, D) The time series shown in panels B and D were computed from a synthetic dynamics without PAC, as described in Section 2.8 using the algorithms 1 and 2, respectively. The synthetic dynamics used in panels B and D was computed using the same set of hyperparameter values as those described in the caption of Figure B.2, with the exception of the extrinsic noise *η(t)* which in this case was set to 30 percent of the maximum amplitude of the deterministic part of signal *x(t)* (i.e first term of the right-hand member of the Eq. A.1). In the panels A, B, C and D, the solid lines represent the mean values and the shaded error bars correspond to the standard deviation of 100 realizations at each point.

The behavior of time series of PAC metrics shown in Figures 11A and 11B associated to the confounding of Algorithm 1 has been also observed during the transition between the pre-ictal to ictal periods in intracerebral electroencephalography recordings (LFP: local field potential) obtained from the seizure onset zone of epilepsy patients [11] (data not shown). This result is relevant since several CFC types, in particular PAC, have been proposed as biomarkers for detecting the seizure onset in epilepsy patients. As a conclusion, our results suggest that Algorithm 1 should be avoided in analyzing oscillatory dynamics characterized by abrupt changes of amplitude, where the Algorithm 2 is recommended instead. In addition, the temporal evolution of CFC metrics around transient dynamics involving abrupt changes of amplitude (or any other feature), like those associated to epileptic seizures, should be analyzed and interpreted carefully. The TLI time series shown in Figure 11 present more dispersion and a higher baseline bias when compared to that of Figure 9B due to the fact that they were computed for a more noisy synthetic dynamics and using a shorter sliding epoch. Importantly, since the TLI metric is entirely computed in the time domain using band-pass filtered signals, it is not affected by the confounding produced by the Algorithm 1 associated to the computation of features using the Hilbert transform.

We characterized quantitatively the harmonic content of CFC patterns using the harmonicity-CFC plots which categorize the analyzed oscillatory dynamics in four quadrants, Q1: harmonic CFC, Q2: harmonic oscillations and No CFC, Q3: Non harmonic oscillations and No CFC, Q4: Non harmonic CFC. Figure 12 shows the harmonicity-PAC plot, computed as it was described in Section 2.6, for a variety of synthetic oscillatory dynamics and taking the amplitude modulation depth as a parameter.

**Figure 12:**
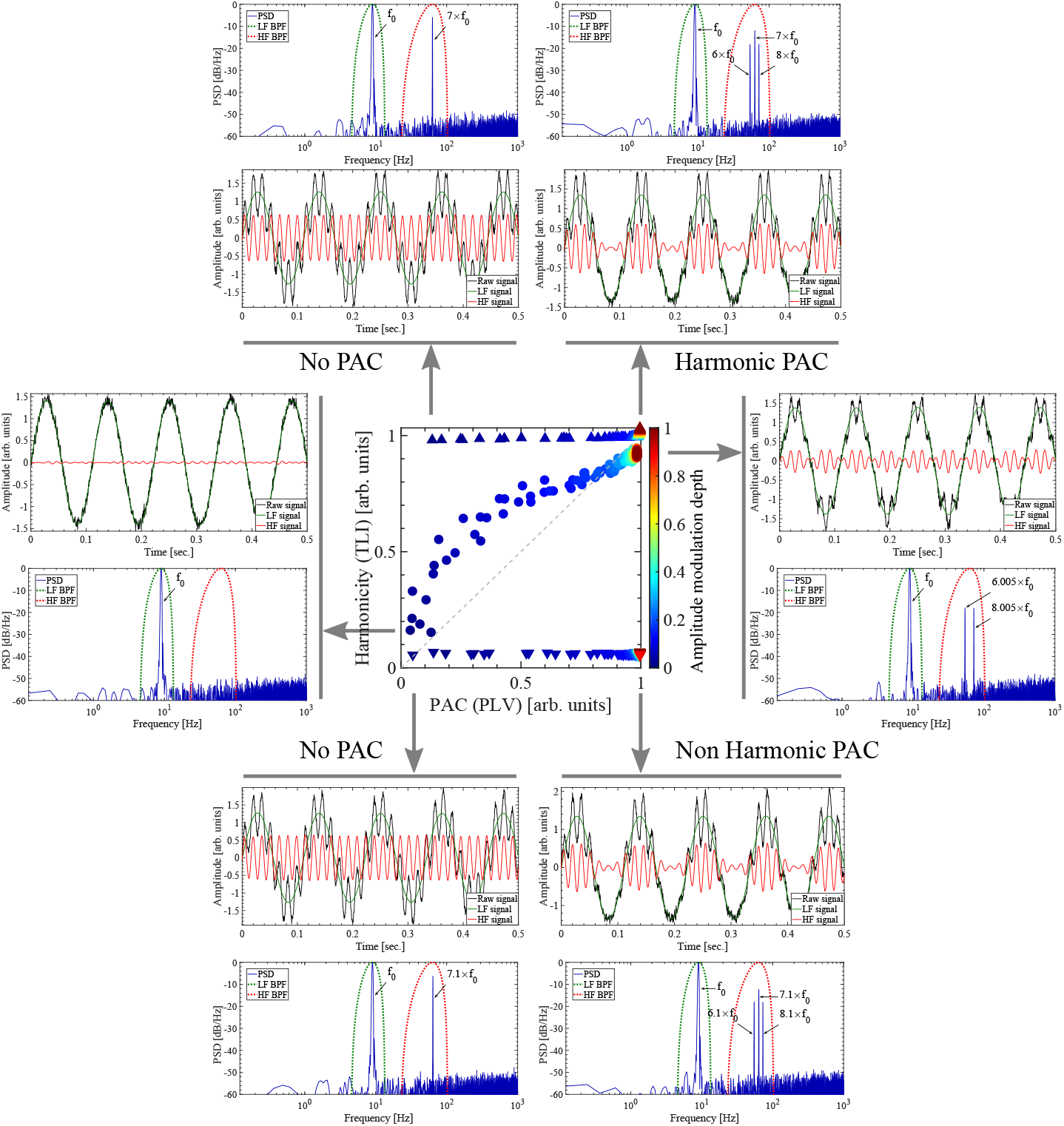
Harmonicity-PAC plot computed for a variety of synthetic oscillatory dynamics and taking the amplitude modulation depth as a parameter. The Harmonicity-PAC plot was computed as it is described in Section 2.6. The pseudocolor scale represents the 1 - m values ranging from the minimum 0 (blue) to the maximum 1 (red) modulation depth, with m defining the amplitude modulation depth as stated in Eq. A.4. The synthetic dynamics were computed as it is described in Appendix A.1 using sinusoidal modulating signals (Eq. A.6).

### 3.2. A single non sinusoidal oscillatory dynamics characterized by dependent frequencies

In this section we investigate the robustness of the proposed (TLI) and conventional (*PLV_PPC_*) harmonicity measures to quantify the harmonic content of the simulated dynamics of the Van der Pol oscillator (see Appendix A.2).

We shall show that, contrary to what is usually assumed in the literature, the spurious PAC elicited by a single non sinusoidal oscillatory dynamics like the one associated to the Van der Pol oscillator can produce both harmonic and non harmonic PAC patterns.

Figure 13 shows that, in the case of oscillatory dynamics containing multiharmonic HF components, the TLI is more robust than the *PLV_PPC_* against changes in the bandwidth of the BPF used to compute the high frequency component (HF BPF, see the dotted and solid red lines in panels A, B, C and D of Figure 13). The non sinusoidal oscillatory dynamics shown in Figure 13, is constituted by a fundamental component at *f_d_* = 5.56 Hz and odd harmonic components at *N* × *f*_0_ with *N* = 3, 5, 7, 9, 11, 13,· · ·. In Figure 13, the bandpass filters used to compute the harmonicity metrics were centered at *f_LF_* = 1×5.56 Hz (LF BPF) and *f_HF_*= 9×5.56 Hz (HF BPF), and consequently, the *PLV_PPC_* metric was computed using Eq. 4 with *M* = 1, *N* = 9. Figure 13E shows that the drop of the *PLV_PPC_* value occurs concurrently with the increase of the phase clustering *PC_HF_*, indicating that the former is produced by a non uniform distribution of the phase values associated to the HF component, i.e. non sinusoidal *x*_H F_(solid red line in Figure 13D). Worthy to note, Figure 13E also shows that increments of the HF bandwidth up to *Bw_HF_* ≈ 20 Hz degrade the signal-to-noise ratio in the band-pass filtered signal *x*HF producing a moderate drop of the TLI value. On the other hand, for *Bw_HF_* ≳ 20 Hz, the HF bandwidth is wide enough to include other harmonic components and thus improving the signal-to-noise ratio of *x_HF_* which translates in that the TLI values become closer to the unity again. Importantly, we found that the pairwise phase consistency measure [23, 24] is also affected by the phase clustering *PC_HF_* presenting a similar behavior to that shown by the *PLV_PPC_* metric in Figure 13E (data not shown). This results is not surprising since the pairwise phase consistency measure was algorithmically derived from the PLV metric [23].

**Figure 13:**
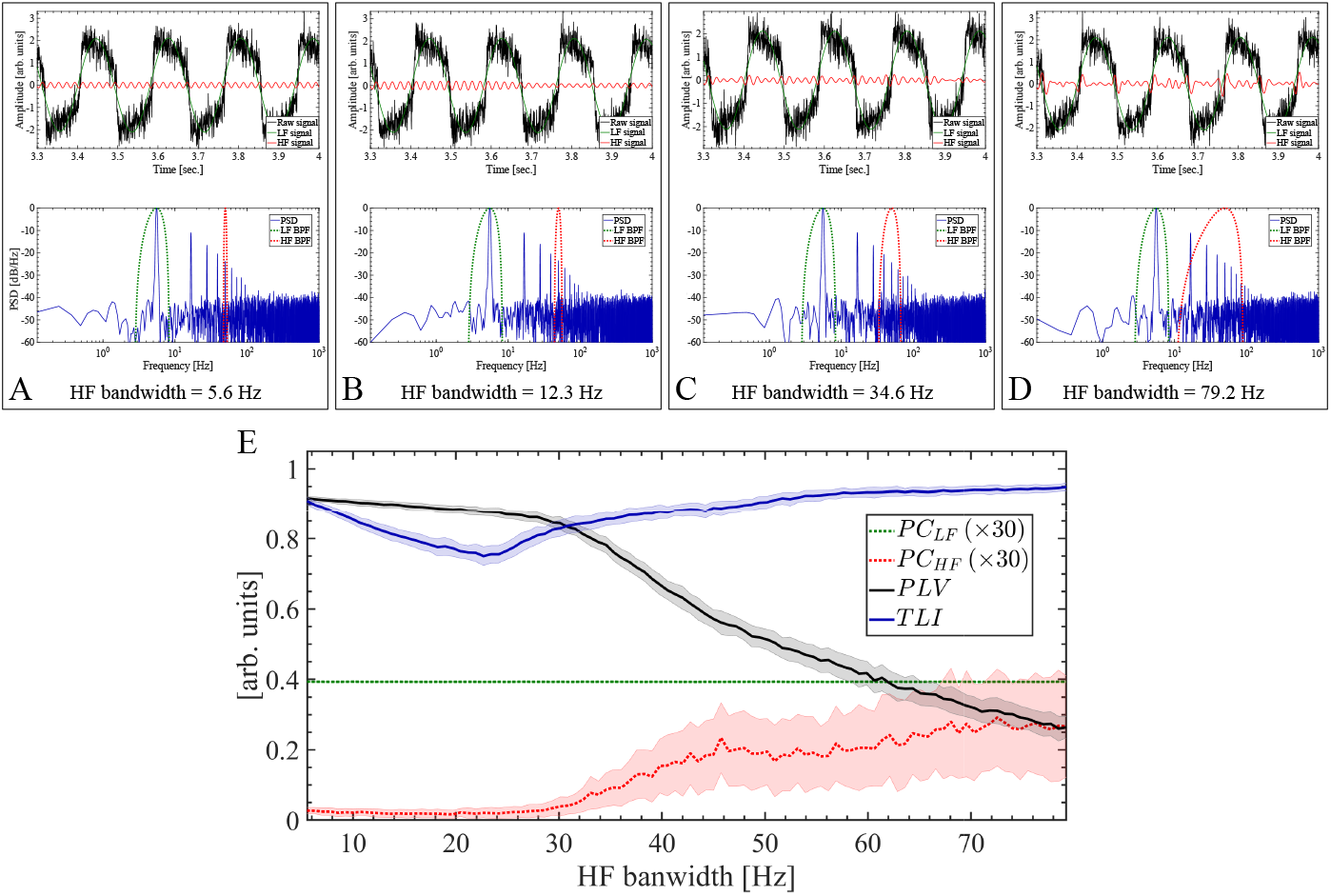
The TLI is more robust than the *PLV_PPC_* against changes in the bandwidth of the BPF used to compute the high frequency component. The Van der Pol oscillatory dynamics (solid black line) shown in panels A, B, C and D were simulated as described in Appendix A.2 using the following hyperpameter values: sampling rate *f_s_* = 2000 Hz, μ = 300, ω_0_ = 2π*f_0_*, *f_o_* = 10 Hz, *W_p_*= 0, *F_e_*= 0, initial conditions *x*(0) = 2, *ẋ*(0) = 1. With this configuration, we obtain a non sinusoidal oscillatory dynamics constituted by a fundamental component at *f_d_* = 5.56 Hz and odd harmonic components at *N* × *f_d_* with *N* = 3, 5, 7, 9, 11, 13, · · ·. The dynamics was computed without intrinsic noise (*g*_1_ = *g*_2_ = 0 in Eq. A.14) in order to obtain a non sinusoidal oscillatory dynamics with a constant fundamental period. Extrinsic noise *η(t)* was added as shown in Eq. A.14. In this case the noise level corresponds to the 20 percent of the maximum amplitude of the deterministic part of signal (i.e x1(t) in Eq. A.14), scaling the standard deviation σ of the additive white Gaussian noise 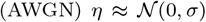. We computed the Van der Pol dynamics for 6 sec. time interval and then the first 0.1 sec. (200 samples) of the time series were discarded to remove the transient period of the numerical simulation. For this set of hyperparameter values, panels A, B, C, D show different realizations of the Van der Pol dynamics. To obtain all the band-pass filtered signals shown in this figure we use the BPF as described in Appendix A.5. The bandwidth of the BPF for the LF component (LF BPF) was kept fixed at *Bw_LF_* = *f_d_*= 5.56 Hz (see the dotted green lines superimposed to the power spectra shown in panels A, B, C, D). The band-pass filters used to compute the harmonicity metrics were centered at *f_LF_* = 1 × *f_d_*Hz (LF BPF) and *f_HF_* = 9 × *f_d_*Hz (HF BPF), and consequently, the *PLV_PPC_* metric was computed using Eq. 4 with *M* = 1, *N* = 9. In panels A, B, C, D we changed the bandwidth of the BPF for the HF component (HF BPF) whose power response (i.e. square magnitude) is shown as dotted red line superimposed to the power spectra. The resulting band-pass filtered HF signals are shown as solid red lines in the upper graph of panels A, B, C, D. The power spectra and the harmonicity metrics (TLI, *PLV_PPC_*) were computed using an epoch length of ≈ 33/*f_d_*≈ 6 sec. The panel (E) shows the magnitude of the harmonicity (TLI, *PLV_PPC_*) and phase clustering metrics (PCLF, PCHF) as a function of the bandwidth of the BPF for the HF component (HF BPF). In the panel E, the solid lines represent the mean values and the shaded error bars correspond to the standard deviation of 100 realizations at each HF bandwidth value.

We also investigate the effect of the bandwidth associated to the BPF used to compute the low frequency component (LF BPF, see the dotted and solid green lines in panels A, B and C of Figure 14). In this case the filters were centered at *f_LF_* = 5 × 5.56 Hz (LF BPF) and *f_HF_* = 15 × 5.56 Hz (HF BPF), and the *PLV_PPC_* metric was computed using Eq. 4 with *M* = 5, *N* = 15. Figure 14D shows that both harmonicity metrics are degraded by the increase of the phase clustering *PC_LF_* associated to a non sinusoidal low frequency component ċLF (see the solid green line in Figure 14C). That is, as the *Bw_LF_* is increased to include several harmonic components within its bandwidth, a non sinusoidal LF components is obtained at the output of the LF BPF filter (see the solid green line in Figure 14C). This in turns produces an increment in the phase clustering (*PC_LF_*), biasing the intensity of both metrics (TLI, *PLV_PPC_*) toward values close to zero. Un this condition, both harmonicity metrics (TLI, *PLV_PPC_*) fail to detect the presence of harmonic components in the oscillatory dynamics.

**Figure 14:**
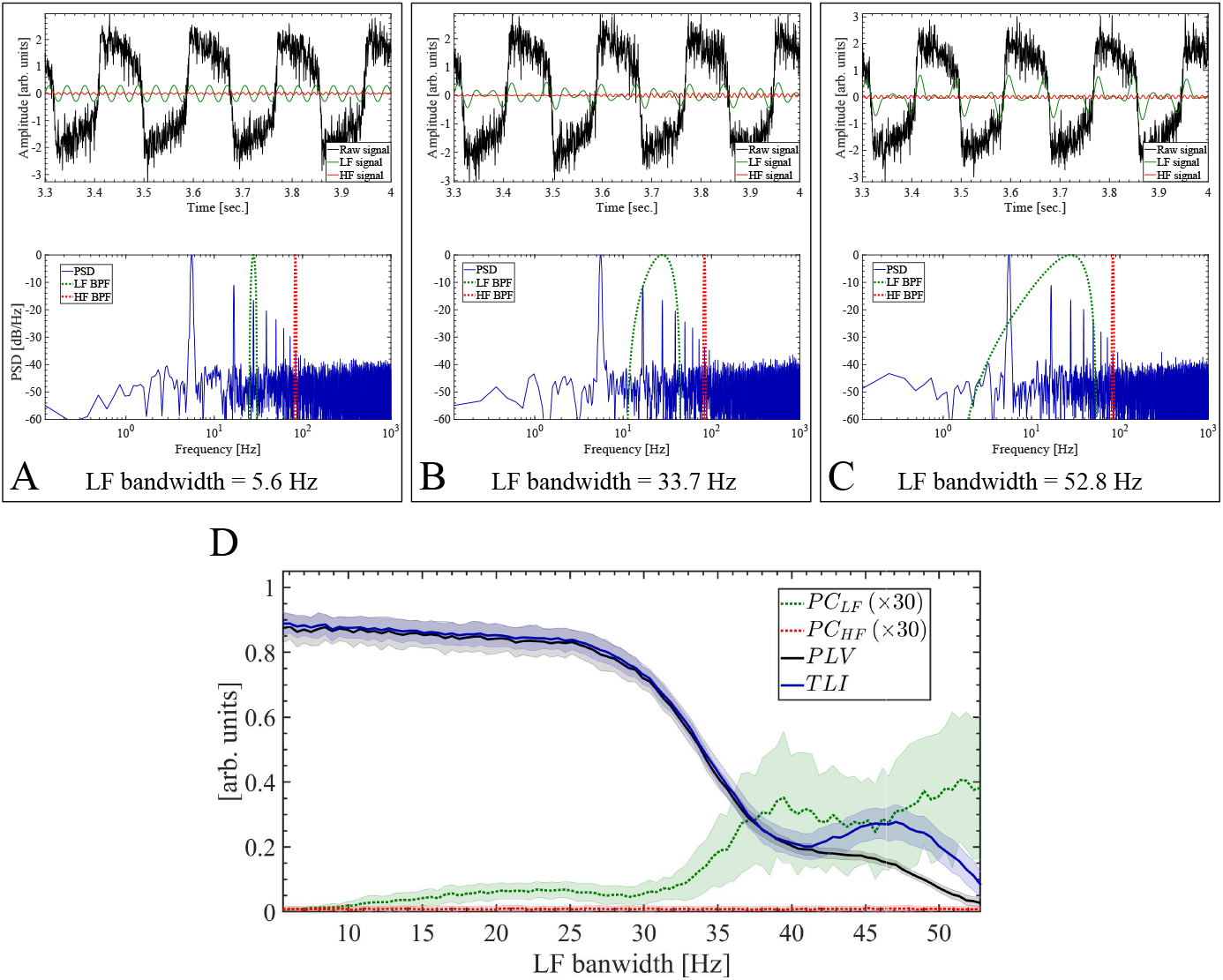
The phase clustering associated to the low frequency component (PCLF) produces a bias in both TLI and *PLV_PPC_* metrics. In this figure we use the same synthetic dynamics and the same set of hyperparameter values to compute the metrics than those described in the caption of Figure 13, except for the configuration of the BPFs used to compute the LF and HF components. In this case, the bandwidth of the BPF for the HF component (HF BPF) was kept fixed at *Bw_HF_* = *f_d_* = 5.56 Hz (see the dotted red lines superimposed to the power spectra shown in panels A, B, C). Besides, the band-pass filters used to compute the harmonicity metrics were centered at *f_LF_* = 5 × *f_d_*Hz (LF BPF) and *f_HF_* = 15 × *f_d_*Hz (HF BPF), and consequently, the *PLV_PPC_* metric was computed using Eq. 4 with *M* = 5, *N* = 15. In panels A, B, C we changed the bandwidth of the BPF for the LF component (LF BPF) whose power response (i.e. square magnitude) is shown as dotted green line superimposed to the power spectra. The resulting band-pass filtered LF signals are shown as solid green lines in the upper graph of panels A, B, C. The power spectra and the harmonicity metrics (TLI, *PLV_PPC_*) were computed using an epoch length of ≈ 33/*f_d_* ≈ 6 sec. The panel D shows the magnitude of the harmonicity (TLI, *PLV_PPC_*) and phase clustering metrics (PCLF, PCHF) as a function of the bandwidth of the BPF for the LF component (LF BPF). In the panel D, the solid lines represent the mean values and the shaded error bars correspond to the standard deviation of 100 realizations at each LF bandwidth value.

Any non linear oscillator can be used as a model that generates spurious PAC via separation of time scales due to non linear effects. Importantly, these emerging time scales elicited by non linearities of the system are not independent from each other, but harmonically related and dependent on the waveform shape of the resulting non sinusoidal oscillatory dynamics. Figure 15C shows the harmonicity-PAC plot associated to the single oscillatory dynamics of the Van der Pol oscillator (see Appendix A.2). Specifically, Figure 15C shows the evolution of the PAC (PLV) and harmonicity (TLI) metrics as the non linear parameter of the oscillator (μ/ω_0_) is increased from the sinusoidal oscillatory regime (μ/ω_0_ ≈ 0, see panels A and F in Figure 15) up to a high non sinusoidal regime (μ/ω_0_ ≈ 4.77, see panels D and K in Figure 15). In Figure 15, the sinusoidal oscillatory regime (μ/ω_0_ ≈ 0) shown in panels A and F, becomes evident by the single spectral component constituting the corresponding power spectra (see panels B and G), and by the phase portraits shown in Figures B.6A and B.6B. On the other hand, the non sinusoidal oscillatory regime (μ/ω_0_ ≈ 4.77) shown in panels D and K, becomes evident by the harmonic spectral components constituting the corresponding power spectra (see panels E and L), and by the phase portraits shown in Figures B.6C,D. In Figures 15A,B,C,D,E, the dynamics of the Van der Pol oscillator was computed by configuring the intrinsic noise of type AWGN applied only on the equation of *ẋ_2_(*g*_1_* = 0 and *g*_2_= 0.5 in Eq. A.14). In this scenario, as the non sinusoidal oscillatory regime emerges, the harmonicity metric (TLI) allows for a clear identification of the harmonic nature of the PAC pattern. Figure 15H shows the behavior of the PAC pattern in the case when AWGN is being applied on the equations of both *ẋ*_1_ and *ẋ_2_* (i.e. *g*_1_= *g*_2_= 0.5 in Eq. A.14). On the other hand, Figure 15H shows that the harmonicity of the PAC intensity increases up to a given value of the non linear paremeter of the oscillator (*μ/ω*_0_), after which subsequent increments of *μ/ω*_0_ produce a monotonic decrease of the harmonicity and keeping the PAC intensity unchanged. Figure 15 shows that the single oscillatory dynamics of the Van der Pol oscillator in presence of AWGN can elicit several PAC patterns depending on the value of *μ/ω*_0_: no PAC (Figures 15F,G), harmonic PAC (Figures 15I,J) and non harmonic PAC (Figures 15K,L). The results show in Figure 15H were computed using an epoch of 10 sec. which corresponds to approx. 50 cycles of the slowest oscillation at *f_LF_* ≈ 4.7 for *μ/ω*_0_ ≈ 4.8. Importantly, it was found that these results holds even in the case of using an epoch length of 1.5 sec. (≈ 7 cycles of the slowest oscillation), which is one order of magnitude shorter than that involved in the computation of Figure 15H. Moreover, we found that the results presented in Figure 15 hold for the dynamics of the Van der Pol oscillator simulated with intrinsic noise of the type non-additive white Gaussian noise (NAWGN). The Harmonicity-PAC plots computed for the simulated dynamics of the Van der Pol oscillator with NAWGN intrinsic noise are shown in Figure B.7 of Appendix B.3. In addition, we verified that these results also hold when PAC is assessed using different metrics (e.g. PLV, KLMI), hence, discarding the possibility of artifacts associated to a particular metric (compare panels A vs. B and C vs. D shown in Figure B.7 of Appendix B.3). These results suggest that the presence of intrinsic noise (AWGN or NAWGN) can change the period of the single oscillatory dynamics in almost a cycle-by-cycle manner significantly reducing the harmonic content in its power spectrum. This evidence supports the conclusion that ‘true’ and ‘spurious’ concepts applied to the CFC patterns are not intrinsically linked to the harmonic content of the underlying oscillatory dynamics. More specifically, the high harmonic content observed in a given oscillatory dynamics is neither sufficient nor necessary condition to interpret the associated CFC pattern as ‘spurious’ or epiphenomenal (i.e. a CFC pattern not representing a true interaction between two coupled oscillatory dynamics with independent fundamental frequencies). For instance, a single oscillatory dynamics characterized by a non constant oscillation period can produce ‘spurious’ CFC with low harmonic content (i.e. non harmonic CFC). This type of oscillatory dynamics is commonly observed in oscillators undergoing a chaotic regime or non linear oscillators under the effect of intrinsic noise (Figure 15H). On the other hand, in Sections 3.3 and 3.4 we shall present results supporting the hypothesis that two coupled oscillatory dynamics with independent fundamental frequencies can elicit ‘true’ CFC with high harmonic content via rectification mechanisms.

**Figure 15:**
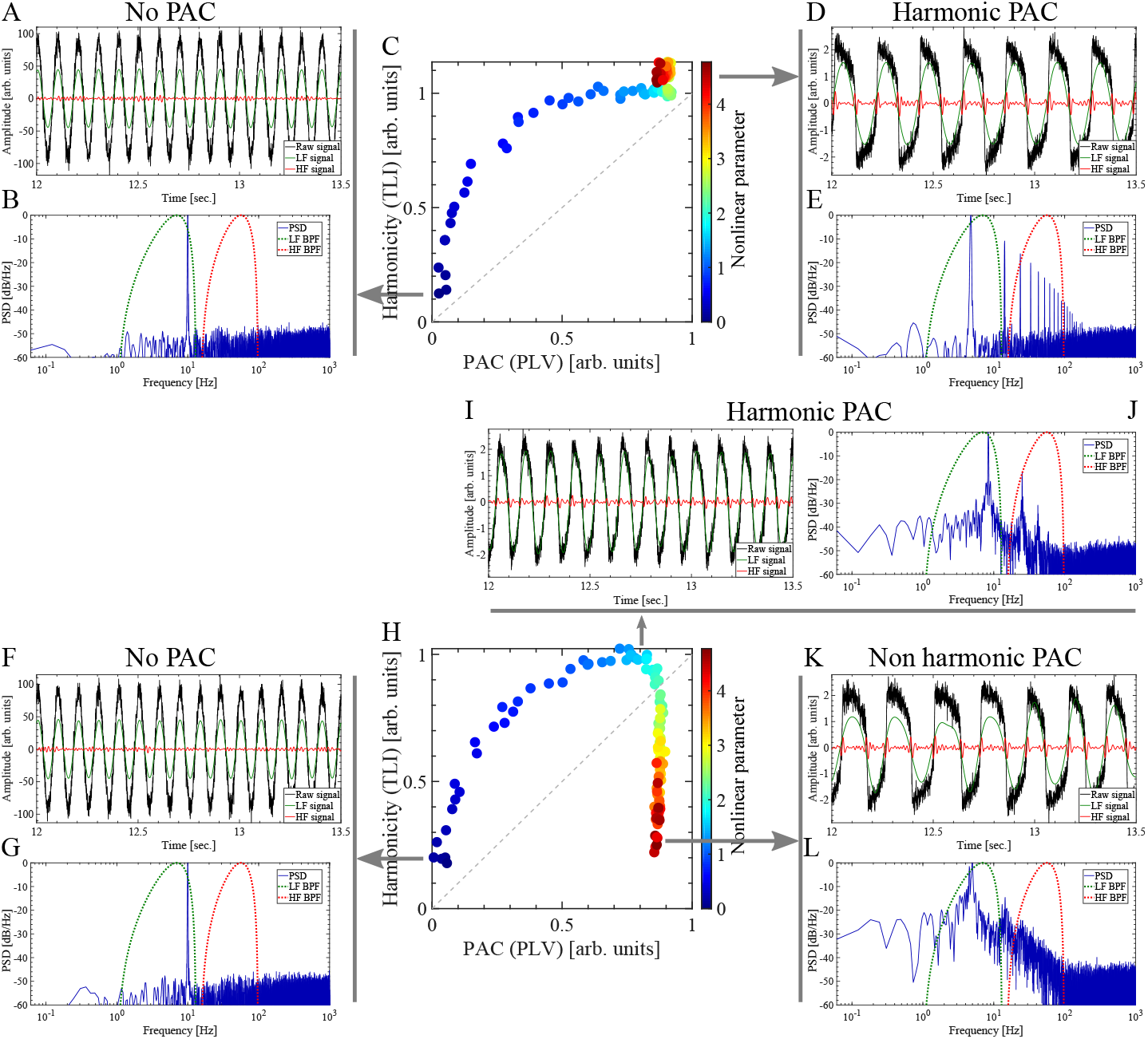
Harmonicity-PAC plot computed for the simulated dynamics of the Van der Pol oscillator with intrinsic noise of type additive white Gaussian noise (AWGN). Note that a single non sinusoidal oscillatory dynamics can produce both harmonic (panels D and E, I and J) and non harmonic (panels K and L) PAC patterns. The Van der Pol oscillatory dynamics (solid black line) shown in panels A, D, F, I and K were simulated as described in Appendix A.2 using the following hyperpameter values: sampling rate *f*_s_ = 2000 Hz, *ω*_0_ = 2*πf*_0_, *f*_0_ = 10 Hz, *W_p_* = 0, *F_e_* = 0, initial conditions x(0) = 2, *ẋ*(0) = 1. To compute the harmonicity-PAC plots shown in panels C and H, the parameter μ controlling the oscillator nonlinearity was increased from the sinusoidal oscillatory regime (μ/ω_0_ ≈ 0, see panels A, B, F, G) up to a high non sinusoidal regime (μ/ω_0_ ≈ 4.77, see panels D, E, I, J, K, L). In panels C and H, the pseudocolor scale represents the μ/ω_0_ values ranging from ≈ 0 (blue) to ≈ 4.8 (red). In panels A, B, C, D and E, the dynamics of the Van der Pol oscillator was simulated using intrinsic noise of type AWGN applied only on the equation of *ẋ_2_(*g*_1_* = 0 and *g*_2_= 0.5 in Eq. A.14). In panels F, G, H, I, J, K and L, the dynamics was simulated by applying the intrinsic noise of type AWGN on the equations of both *ẋ*_1_ and *ẋ*_2_ (i.e. *g*_1_= *g*_2_=0.5 in Eq. A.14). Therefore, in this case the intrinsic noise components (AWGN) in Eq. A.14 result 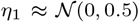 and 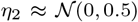. Extrinsic noise *η(t)* was added as shown in Eq. A.14. In this case the noise level corresponds to the 10 percent of the maximum amplitude of the dynamics *x*_1_ in Eq. A.14), scaling the standard deviation σ of the additive white Gaussian noise (AWGN) 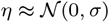. We computed the Van der Pol dynamics for 20 sec. time interval and then the first 10 sec. of the time series were discarded to remove the transient period of the numerical simulation. The power spectra (solid blue line in graphs B, E, G, J and L) were computed unsing a 10 sec. epoch from the synthetic dynamics shown in the corresponding graphs (solid black line in graphs A, D, F, I and K). In graphs B, E, G, J and L, the power responses (i.e. square magnitude) of the BPF used to compute the LF and HF signals are shown as dotted green and red lines, respectively. To obtain all the band-pass filtered signals shown in this figure we use the BPF as described in Appendix A.5. In all the cases shown in this figure, the bandwidth of the BPF for the LF (LF BPF) and HF (HF BPF) components were set at *Bw_LF_* = 12 Hz centered at 7 Hz and *Bw_HF_* = 82.65 Hz centered at 56.5 Hz, respectively. In graphs A, D, F, I and K, the resulting band-pass filtered LF and HF signals are shown as solid green and solid red lines, respectively. The harmonicity metric (TLI) was computed as it was described in Section 2.4. The PAC metric (*PLV_PAC_*) was computed using Eq. 4 with the configuration given by Eq. A.20 and *M* = *N* = 1.

It was found that the dependence of the harmonicity of the CFC pattern on the intrinsic noise (AWGN or NAWGN) is not exclusive of PAC but occurs in several CFC types. Figures 16C and 16H show this effect in the case of AAC and FFC patterns, respectively. To compute the Figure 16C we set Eqs. A.12, A.13 and A.14 with *W_p_* = 0, *f_e_* = 5.4 Hz and *f_m_* = 1.33 Hz. This configuration produces an amplitude-modulated dynamics due to the action of the external driving *F_e_*. This configuration produces an amplitude-modulated dynamics due to the action of the amplitude-modulated external driving *F_e_* in which the sinusoidal component at *f_m_* = 1.33 Hz modulates the amplitude of the oscillation at *f_e_* = 5.4 Hz (dotted grey line in Figures 16A and 16D). As a consequence, the slow rhythm at *f_m_* = 1.33 Hz effectively modulates the amplitude of the non sinusoidal oscillator dynamics (solid black line in Figures 16A and 16D). The amplitude-modulation of the resulting non sinusoidal dynamics becomes evident in the phase portraits shown in Figure B.8. Thus, two CFC patterns emerge from the resulting dynamics, (1) a PAC pattern in which the phase of the slow rhythm at *f_m_* = 1.33 Hz amplitude modulates the fundamental component of the non sinusoidal oscillator dynamics, and (2) an AAC pattern in which the amplitude of the harmonic components follow the changes of the fundamental component amplitude. In the PAC pattern we have a ‘true’ interaction between two oscillatory dynamics, i.e. *f_m_* = 1.33 Hz andfe = 5.4Hz. On the other hand, the AAC pattern can be thought as a ‘spurious’ or epiphenomenal coupling since it involves dependent frequencies related by the waveform shape of the single oscillatory dynamics. Figure 16C shows that the AAC intensity increases up to a given value of the external driving amplitude (Ae in Eq. A.13), after which subsequent increments of *A_e_* produce a significant drop in the harmonicity of the ‘spurious’ AAC pattern. To compute the Figure 16H we set Eqs. A.12, A.13 and A.14 with *F_e_* = 0, *f_0_* = 10 Hz, *f_p_* ≈ 1 Hz. As a result, we obtain an frequency-modulated dynamics due to the action of the time variant parameter *W_p_*. Specifically, the slow rhythm at *f_p_*≈ 1 Hz (dotted gray line in Figures 16F and 16I) effectively modulates the fundamental frequency of the non sinusoidal oscillator dynamics (solid black line in Figures 16F and 16I). As a consequence, two CFC patterns emerge from the resulting dynamics, (1) a PFC pattern in which the phase of the slow rhythm at *f_p_* ≈ 1 Hz frequency modulates the fundamental component of the non sinusoidal oscillator dynamics, and (2) an FFC pattern in which the instantaneous frequency of the harmonic components follow the changes of the fundamental component frequency. In the PFC pattern we have a ‘true’ interaction between two oscillatory dynamics, one associated to the time variant parameter *W_p_* and the other to the intrinsic dynamics of the oscillator. On the other hand, the FFC pattern can be thought as a ‘spurious’ or epiphenomenal coupling since it involves dependent frequencies related by the waveform shape of the single oscillatory dynamics. Figure 16H shows that the FFC intensity increases up to a given value of the *W_p_* intensity (i.e. *A_p_* in Eq. A.12), after which subsequent increments of *A_p_* produce a significant drop in the harmonicity of the ‘spurious’ FFC pattern.

**Figure 16:**
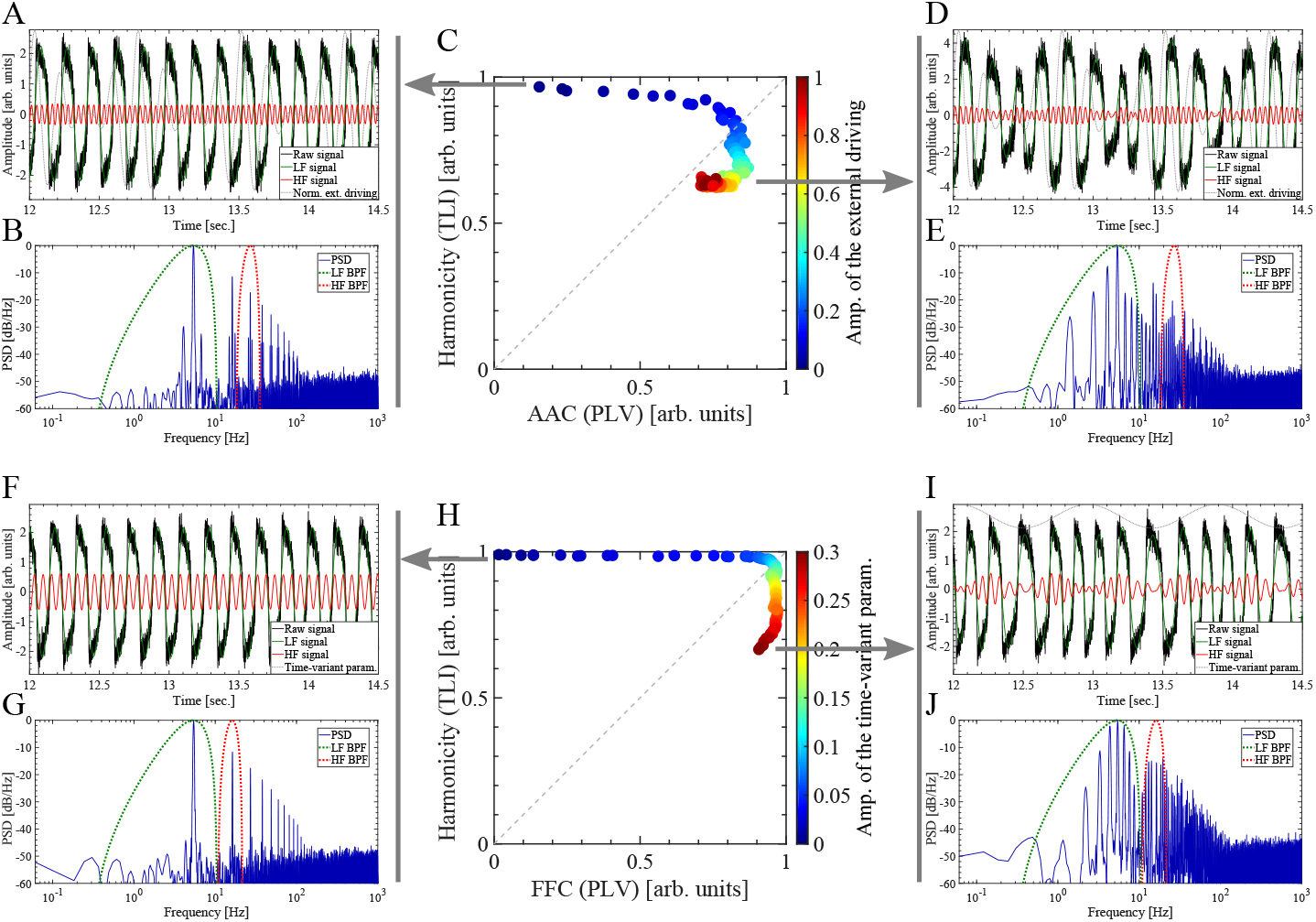
Harmonicity-AAC and Harmonicity-FFC plots computed for the simulated dynamics of the Van der Pol oscillator with intrinsic noise of type additive white Gaussian noise (AWGN). (A, B, C, D, E) To obtain the AAC pattern, the Van der Pol oscillatory dynamics (solid black line) shown in panels A and D were simulated as described in Appendix A.2 using Eqs. A.12, A.13 and A.14 with the following hyperpameter values: sampling rate *f_s_* = 2000 Hz, ω_0_ = 2π*f_0_*, *f_0_* = 10 Hz, no parametric driving *W_p_* = 0, *f_e_* = 5.4 Hz and *f_m_* = 1.33 Hz, Am = 1, c = 1 (i.e. DSB-C), maximum modulation depth m = 0, initial conditions *ẋ*(0) = 2, *ẋ*(0) = 1. To compute the harmonicity-AAC plot shown in panel C, the external driving amplitude (Ae in Eq. A.13) was increased from *A_e_* = 0 (no external driving) up to *A_e_* = 5 × 10^4^. In panel C, the pseudocolor scale represents the *A_e_*/(5 × 10^4^) values ranging from 0 (blue) to 1 (red). (F, G, H, I, J) To obtain the FFC pattern, the Van der Pol oscillatory dynamics (solid black line) shown in panels F and I were simulated as described in Appendix A.2 using Eqs. A.12, A.13 and A.14 with the following hyperpameter values: sampling rate *f_s_* = 2000 Hz, ω_0_ = 2π*f_0_*, *f_0_* = 10 Hz, no external driving *F_e_* = 0, *f_p_* ≈ 1 Hz. To compute the harmonicity-FFC plot shown in panel H, the intensity of the time variant parameter *W_p_* (*A_p_* in Eq. A.12) was increased from *A_p_* = 0 (no parametric driving) up to *A_p_* ≈ 10. In panel H, the pseudocolor scale represents the Ap/34 values ranging from 0 (blue) to 0.3 (red). For both panels C and H, the dynamics of the Van der Pol oscillator was simulated using intrinsic noise of type AWGN applied only on the equation of *ẋ*_2_*(g_1_* = 0 and *g*_2_= 0.5 in Eq. A.14). Therefore, the intrinsic noise components (AWGN) in Eq. A.14 result η1 = 0 and 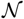(0, 0.5). Extrinsic noise *η(t)* was added as shown in Eq. A.14. In this case the noise level corresponds to the 10 percent of the maximum amplitude of the dynamics x1 in Eq. A.14), scaling the standard deviation σ of the additive white Gaussian noise (AWGN) η ≈ 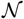(0, σ). The Van der Pol dynamics was computed for 20 sec. time interval and then the first 10 sec. of the time series were discarded to remove the transient period of the numerical simulation. The power spectra (solid blue line in graphs B, E, G and J) were computed unsing a 10 sec. epoch from the synthetic dynamics shown in the corresponding graphs (solid black line in graphs A, D, F and I). In graphs B, E, G and J, the power responses (i.e. square magnitude) of the BPF used to compute the LF and HF signals are shown as dotted green and red lines, respectively. To obtain all the band-pass filtered signals shown in this figure we use the BPF as described in Appendix A.5. In computing panel C (AAC), the bandwidth of the BPF for the LF (LF BPF) and HF (HF BPF) components were set at *Bw_LF_* ≈ 10.3 Hz centered at 5.4 Hz and *Bw_HF_* ≈ 17.6 Hz centered at 27 Hz, respectively. In computing panel H (FFC), the bandwidth of the BPF for the LF (LF BPF) and HF (HF BPF) components were set at *Bw_LF_* ≈ 10.3 Hz centered at 5.4 Hz and *Bw_HF_* ≈ 10.8 Hz centered at 16.2 Hz, respectively. In graphs A, D, F and I, the resulting band-pass filtered LF and HF signals are shown as solid green and solid red lines, respectively. The harmonicity metric (TLI) was computed as it was described in Section 2.4. For the panel C, the AAC metric (*PLV_AAC_*) was computed using Eq. 4 with the configuration given by Eq. A.21 and *M* = *N* = 1. For the panel H, the FFC metric (*PLV_FFC_*) was computed using Eq. 4 with the configuration given by Eq. A.24 and *M* = *N* = 1.

### 3.3. Two coupled oscillatory dynamics characterized by independent frequencies

In this section we present the results obtained with a 2nd order parametric oscillator showing that two coupled oscillatory dynamics with independent fundamental frequencies can elicit ‘true’ CFC with high harmonic content via the rectification mechanism. The equations describing the dynamics of the parametric oscillator are detailed in Section 3.3. Figure 17 shows the PFC patterns corresponding to the dynamics of the parametric oscillator generated by simultaneously applying an off-resonance external driving *F_e_* and a parametric driving *W_p_* tuned at the same frequency *f_e_* = *f_p_* = *f_0_* /12 ≈ 8.3 Hz and θe = 0 (see Eqs. A.16 and A.17), with *f_0_* being the natural resonance frequency of the undamped oscillator (μ = 0 in Eq. A.15). Figures 17C and 17J show the harmonicity-PFC plots for the cases when AWGN is applied only on the equation of *ẋ_2_(g_1_* = 0 and *g*_2_= 0.125 in Eq. A.18) and on the equations of both *ẋ*ι and *ẋ*_2_(i.e. g_1_= *g*_2_= 0.125 in Eq. A.18), respectively. In the latter case, the intrinsic noise is capable to drive the resonator at its natural frequency *f_0_* increasing the harmonicity of the oscillatory dynamics for low *A_p_* values (see Figures 17H, 17I, and 17J). This harmonicity of the oscillatory dynamics for low *A_p_* values is not present when the parametric oscillator is configured with non harmonic frequencies (e.g. *f_e_* = *f_p_* = *f_0_*/11.62 ≈ 8.6 Hz, see Figures B.10H, B.10I and B.10J). Figures 17C 17J show that the harmonicity of the PFC pattern increases as the parametric driving intensity *A_p_* increases. In Figure 17, the almost sinusoidal oscillatory regime (*A_p_*≈ 0) shown in panel A, becomes evident by the single spectral component constituting the corresponding power spectra (see panel B), and by the phase portrait shown in Figure B.9A. On the other hand, the non sinusoidal oscillatory regime (*A_p_* ≈ 0. 9) shown in panels D and F, becomes evident by the harmonic spectral components constituting the corresponding power spectra (see panels E and G), and by the phase portraits shown in Figures B.6E and B.6F. In particular, panels D and E in Figure 17 show that the phase of the slow rhythm (*f_LF_* = *f_e_* = *f_p_* ≈ 8.3 Hz) modulates both amplitude and frequency of the fast oscillation within the range 20 Hz < *f_HF_* < 140 Hz (see Figure 4A and 4B). We found that in the forced parametric oscillator, the fast oscillation constituting the oscillatory dynamics undergo a rectification process associated to the PAC pattern. That is, the amplitude of the HF component (*f_HF_*, solid red line in Figures 17D and 17F) goes to zero at some particular phase of the LF cycle (*f_e_* = *f_p_*, solid green line in Figures 17D and 17F). This periodic rectification process produces that the HF component resets its phase relative to the LF component in each LF cycle. As a consequence, the waveform shape of the resulting oscillatory dynamics is almost the same in each LF cycle even when the slow and fast rhythms have independent frequencies. This repetitive waveform shape (Figures 17D and 17F) is characterized by a high harmonic content in its power spectrum (Figures 17E and 17G) which accounts for the high harmonicity reported by the TLI metric for high *W_p_* values (Figures 17C and 17J). Importantly, we found that harmonic PFC patterns like those shown in Figures 17D and 17F are elicited for high values of the parametric driving *W_p_* irrespective of the ratio of the time scales involved in the parametric oscillator, i.e. harmonic PFC patterns occurs for harmonic (Figure 17) or non harmonic frequency ratios *f_e_*/*f_0_* with *f_e_* = *f_p_* (Figure B.10 in Appendix B.4). We also verified that these results also holds when PFC is assessed using different metrics (e.g. PLV, KLMI), hence, discarding the possibility of an artefact associated to a particular metric (compare Figures B.10 and B.11 in Appendix B.4). These results support the hypothesis that the har-monicity of the PFC pattern shown in Figures 17D, 17F, B.10D, B.10F, B.11D and B.11F are not related to a fine-tuning of the parameters fe, *f_p_* and *f_0_* of the parametric oscillator, but to an emerging rectification mechanism associated to the co-occurrence of PAC and PFC patterns which produce the phase resetting of the modulated HF component in each LF cycle.

**Figure 17:**
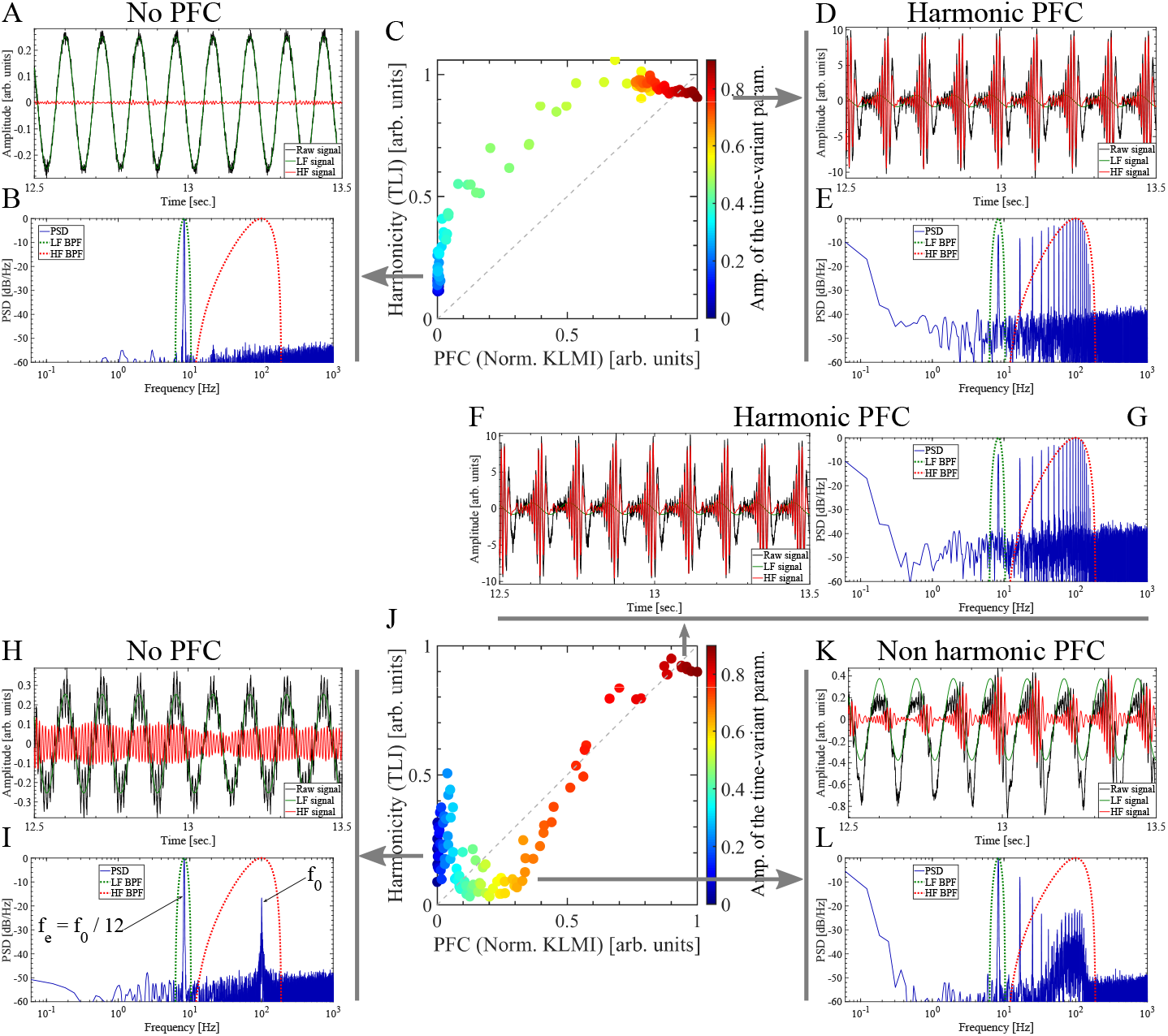
Harmonicity-PFC plot computed for the simulated dynamics of the 2nd order parametric oscillator with intrinsic noise of type additive white Gaussian noise (AWGN). Note that two oscillatory dynamics with independent frequencies can produce harmonic PFC patterns (panels D, E and F, G). The parametric oscillator dynamics (solid black line) shown in panels A, D, H, F and K were simulated as described in Appendix A.3 by simultaneously applying an off-resonance external driving *F_e_* and a parametric driving *W_p_* tuned at the same frequency and using the following hyperpameter values: sampling rate *f_s_* = 2000 Hz, μ = 200, ω_0_ = 2π*f_0_*, *f_0_* = 100 Hz, *f_p_* = *f_e_* = *f_0_*/12 ≈ 8.3 Hz, θe = 0, *A_e_* = 1 × 10^5^. To compute the harmonicity-PFC plots shown in panels C and J, the parameter *A_p_* controlling the parametric driving intensity was increased from the sinusoidal oscillatory regime (*A_p_* ≈ 0, see panels A, B, H, I) up to a high non sinusoidal regime (*A_p_* ≈ 0.9, see panels D, E, F, G). In panels C and J, the pseudocolor scale represents the *A_p_* values ranging from 0 (blue) to 0.9 (red). In panels A, B, C, D and E, the dynamics of the parametric oscillator was simulated using intrinsic noise of type AWGN applied only on the equation of *ẋ*_2_(g_1_= 0 and *g*_2_= 0.125 in Eq. A.18). In panels F, G, H, I, J, K and L, the dynamics was simulated by applying the intrinsic noise of type AWGN on the equations of both *ẋ*_1_ and *ẋ*_2_(i.e. *g*_1_= *g*_2_= 0.125 in Eq. A.18). Therefore, in this case the intrinsic noise components (AWGN) in Eq. A.18 result η1 ≈ 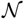(0, 0.125) and η2 ≈ 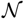(0, 0.125). Extrinsic noise *η(t)* was added as shown in Eq. A.14. In this case the noise level corresponds to the 5 percent of the maximum amplitude of the dynamics x1 in Eq. A.18), scaling the standard deviation σ of the additive white Gaussian noise (AWGN) η ≈ 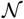(0, σ). We computed the dynamics of the parametric oscillator for 20 sec. time interval and then the first 10 sec. of the time series were discarded to remove the transient period of the numerical simulation. The power spectra (solid blue line in graphs B, E, G, I and L) were computed unsing a 10 sec. epoch from the synthetic dynamics shown in the corresponding graphs (solid black line in graphs A, D, F, H and K). In graphs B, E, G, I and L, the power responses (i.e. square magnitude) of the BPF used to compute the LF and HF signals are shown as dotted green and red lines, respectively. To obtain all the band-pass filtered signals shown in this figure we use t he BPF as described in Appendix A.5. In all the cases shown in this figure, the bandwidth of the BPF for the LF (LF BPF) and HF (HF BPF) components were set at *Bw_LF_* ≈ 4.2 Hz centered at *f_0_*/12 ≈ 8.3 Hz and *Bw_HF_* ≈ 179 Hz centered at *f_0_* = 100 Hz, respectively. In graphs A, D, F, H and K, the resulting band-pass filtered LF and HF signals are shown as solid green and solid red lines, respectively. The harmonicity metric (TLI) was computed as it was described in Section 2.4. The PAC metric (KLMIPFC) was computed using Eqs. 6 and 7 with the configuration given by Eq. A.22. Note that the *KLMI_PFC_* was normalized with its maximum value in each plot.

### 3.4. Biologically plausible neural network model

In this section we show that ‘true’ PAC patterns with high harmonic content (i.e, ‘true’ harmonic PAC) naturally emerge in the oscillatory dynamics of the biologically plausible neural network model shown in Figure 1. This model is considered as a canonical circuit for generating PAC [16], and it represents a network architecture that has been observed in a variety of sensory cortex areas in the form of a slow input stimuli (e.g. visual, auditory, olfactory) which entrain fast gamma oscillations underpinning local neural processing [13]. In addition, the model shown in Figure 1 has been analyzed in the context of the parkinsonian basal ganglia-thalamocortical circuit under dopamine depletion in connection with both the mechanism of action of the deep-brain stimulation therapy [14] and the exaggerated PAC between beta-gamma frequency bands. The latter, putatively associated to the pathological mechanism of motor symptoms in Parkinson’s disease [1, 33, 34].

In [1] we demonstrate that PAC phenomenon naturally emerges in mean-field models of biologically plausible networks, as a signature of specific bifurcation structures. In particular, for the model shown in Figure 1 we found that in the case of an oscillatory external driving without noise (i.e. *H_i_= h_i_* cos(*ω_i_t+ϕ_i_)+d_i_* and *η_i_* = 0 for *I_i_, i* ∈ {1, 2} in Eq. 1), the PAC patterns observed in the resulting dynamics were elicited by the periodic excitation/inhibition (PEI) of a network population producing intermittent fast oscillations (i.e. intermittent PAC). For a detailes discussion of the PEI mechanism associated to the model shown in Figure 1 the reader is referred to Section 3.1 and Appendix A of [1]. The threshold linear activation function *S(I_i_)* (Eq. 2) imposes certain conditions in the input space (*H_1_, H_2_*) for the activation of the two populations constituting the architecture shown in Figure 1. As a consequence, when the amplitude of the inputs are high enough to activate the two populations, the intrinsic fast rhythm (50 Hz) coexist with the external slow driving (ω_i_/(2π) = 3.33 Hz) in the resulting oscillatory dynamics. The fast rhythm cease if any of the two populations is deactivated. The locus in the (*H*_1_, *H*_2_) space defined by the activation conditions does not depends on the temporal evolution of the inputs *H*_1_,*H*_2_ (See Figure 13 in Appendix A of [1]). As a result, the trajectory of a periodic driving dynamics (*h_1_(t)*, *H*_2_(t)) crosses the locus of the activation condition in the same phase of the slow driving period (ω_i_/(2π) = 3.33 Hz). Thus, in the case of oscillatory inputs *H*_1_ and/or *H*_2_ capable to periodically activate and deactivate the populations of the intrinsic oscillator we obtain a PAC pattern associated to the intermittent occurrence of the fast rhythm phase locked to the slow external driving (i.e. PEI mechanism. See Figure 13 in Appendix A of [1]).

Importantly, due to the rectification process involved in the PEI mechanism in presence of threshold linear activation functions, the amplitude of the fast oscillation goes to zero at some particular phase of the slow cycle, hence, the fast oscillation resets its phase relative to the slow driving in each cycle (see Figure 18D). As a consequence, the waveform shape of the resulting oscillatory dynamics is almost the same in each slow cycle even when the slow and fast rhythms have independent frequencies. This repetitive waveform shape (Figure 18D) is characterized by a high harmonic content in its power spectrum (Figure 18E) which accounts for the high harmonicity reported by the TLI metric for high driving amplitude values (Figure 18C). We found that increasing levels of intrinsic noise *η_i_* (see Section 2.1) applied on the model constituted by threshold linear activation function *S(I_i_)* produce a drop in both the harmonicity and the intensity of the PAC as shown in Figures 18C, 18H and 18M. That is, it seems that the harmonic content and the PAC intensity are intrinsically linked by the rectification mechanism associated to threshold linear activation functions.

**Figure 18:**
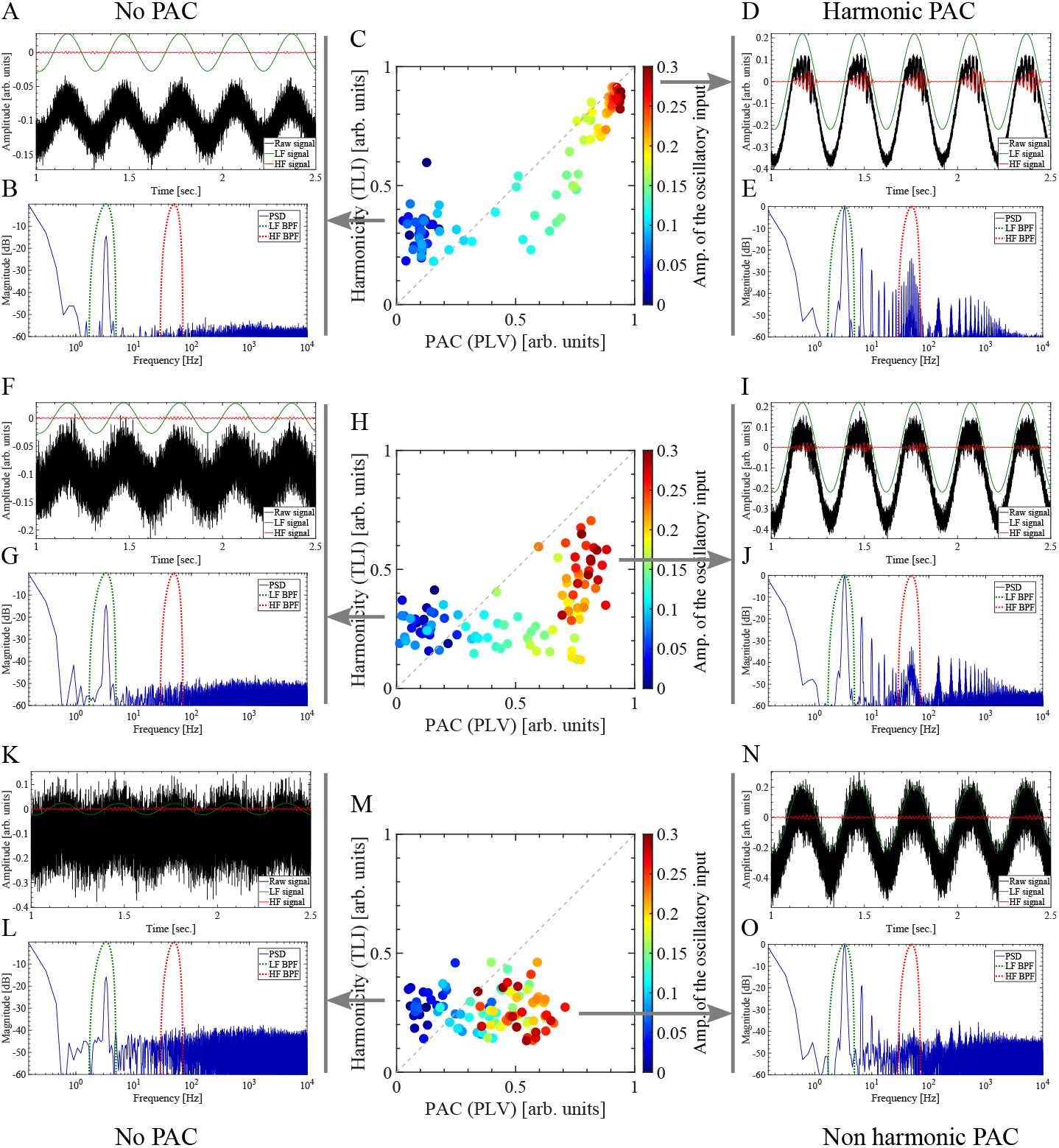
Harmonicity-PAC plot computed for the simulated dynamics of the biologically plausible neural network model shown in Figure 1 using the threshold linear activation function *S*(*I_i_*) (Eq. 2). Note that two coupled oscillatory dynamics with independent fundamental frequencies can produce ‘true’ PAC patterns with high harmonic content via rectification mechanisms (panels D and E). The neural network dynamics (solid black line) shown in panels A, D, F, I, K and N were simulated as described in 2.1 using the configuration detailed in Table 1, resulting in an oscillatory dynamics in the gamma band (50 Hz). Besides, we use the following set of hyperpameter values: sampling rate *f*_s_ = 20 kHz, *H*_1_ = 0, *H*_2_ = *A*_2_ cos(2*πf*_2_ *t*) with *f*_2_ ≈ 3.3 Hz. To compute the harmonicity-PAC plots shown in panels C, H and M, the parameter *A*_2_ controlling the amplitude of the oscillatory input *H*_2_ was increased from *A*_2_ = 0 (see panels A, B, F, G, K, L) up to *A*_2_ = 0.3 (see panels D, E, I, J, N, O). In panels C, H and M, the pseudocolor scale represents the *A*_2_ values ranging from ≈ 0 (blue) to ≈ 0.3 (red). The neural network dynamics was simulated using intrinsic noise *η_i_* of type AWGN to the node inputs *I*i (see Section 2.1). Panels C, H and M were computed with a noise level of 5, 10 and 20 percent of the external input maximum amplitude (*A*_2_ = 0.3) scaling the standard deviation *σ_i_* of the additive white Gaussian noise *η_i_* ≈ 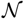(0*, σ*_i_), respectively. We computed theneural network dynamics for 10 sec. time interval and then the first 5 sec. of the time series were discarded to remove the transient period of the numerical simulation. The power spectra (solid blue line in graphs B, E, G, J, L and O) were computed unsing a 5 sec. epoch from the synthetic dynamics shown in the corresponding graphs (solid black line in graphs A, D, F, I, K and N). In graphs B, E, G, J, L and O, the power responses (i.e. square magnitude) of the BPF used to compute the LF and HF signals are shown as dotted green and red lines, respectively. To obtain all the band-pass filtered signals shown in this figure we use the BPF as described in Appendix A.5. In all the cases shown in this figure, the bandwidth of the BPF for the LF (LF BPF) and HF (HF BPF) components were set at *Bw_LF_*≈ 3.3 Hz centered at ≈ 3.3 Hz and *Bw_HF_* ≈ 43.2 Hz centered at 50 Hz, respectively. In graphs A, D, F, I, K and N, the resulting band-pass filtered LF and HF signals are shown as solid green and solid red lines, respectively. The harmonicity metric (TLI) was computed as it was described in Section 2.4. The PAC metric (*PLV_PAC_*) was computed using Eq. 4 with the configuration given by Eq. A.20 and *M* = *N* = 1.

We also investigate the characteristics of the PAC patterns observed in the oscillatory dynamics of the model shown in Figure 1 constituted by the infinitely differentiable softplus activation function defined in Eq. 3. It was found that in absence of noise, the PEI mechanism associated to softplus activation functions elicit PAC patterns with high harmonic content (i.e. harmonic PAC) in the resulting oscillatory dynamics (data not shown). However, in a more realistic scenario including a small level of intrinsic noise *η_i_* applied on the model constituted by softplus activation function *S_c_(I_i_)*, the harmonicity was significantly reduced and the PAC instensity was kept almost unchanged (see Figure B.12 in Appendix B.5). This result suggest that the harmonic content and the PAC intensity are not intrinsically coupled in presence of the softplus activation function and can be interpreted as follows. Due to the fact that *S_c_(I_i_)* >0, the amplitude of the intrinsic fast rhythm (50 Hz) is effectively modulated by the external driving (ω*_i_*/(2π) = 3.33 Hz) but it does not becomes strictly zero at any phase of the slow rhythm, hence, the fast oscillation never resets its phase relative to the slow driving. Thus, the two oscillations with incommensurable frequencies (50 Hz, 3.3 Hz) coupled via the PEI mechanism in absence of phase reseting, produce an oscillatory dynamics similar to that shown in the right panels of Figure 3 (non harmonic PAC).

## 4. DISCUSSION

In this work we provided an in-depth characterization of the Time Locked Index (TLI) as a novel tool aimed to efficiently quantify the harmonic content of noisy time series, and to assist the interpretation of CFC patterns observed in oscillatory dynamics of physical and biophysical systems. It was demonstrated that by operating in the time domain the TLI reliably assesses the degree of time-locking between the slow and fast rhythms, even in the case in which several (harmonic) spectral components are included within the bandwidth of the filter used to obtain the fast rhythm. In this aspect, the TLI measure outperforms the PLV and pairwise phase consistency metrics since the former is more robust against changes in the bandwidth or transition bands steepness of the BPF used to compute the HF component (see Figures 5, 8, 13 and 14 and related discussion).

We exploited the TLI metric together with other complementary signal processing tools to perform the harmonicity analysis on several types of CFC patterns using simulated and synthetic oscillatory dynamics under controlled levels of extrinsic (i.e. of observation) and intrinsic noise. To avoid the introduction of unnecessary timescales on the analyzed oscillatory dynamics, White Gaussian noise (AWGN) was used for this purpose. Since CFC is a rather ubiquitous phenomenon observed in a variety of physical systems, from physiological signals in the endocrine and cardiorespiratory systems, the neural activity of the human brain to the atmospheric variables, astronomical observations, earth seismic waves, nonlinear acoustics and stock market fluctuations (see [1] and references therein), our approach introduces a novel signal processing toolbox (and methodology) relevant to many physical and biophysical disciplines. CFC phenomenon observed in neural recordings has been proposed to be functionally involved in neuronal communication, memory formation and learning. In particular, experimental findings have shown that PAC and PPC patterns are important variants of CFC linked to physiological and pathological brain states like those observed in Parkinson’s disease and epilepsy [1, 11, 31, 34]. As it was discussed in Section 1, we recall that PPC is a signature related to the presence of harmonic spectral components in the underlying oscillatory dynamics. In this regard, the interpretation of the PAC patterns observed in local field potentials (LFP) recorded in humans and animal models remains challenging due to the fact that the brain activity is, in general, characterized by non sinusoidal oscillatory dynamics. The latter raises the question of whether PAC patterns are indicative of true interactions reflecting a mechanistic process between two independent neural oscillators, or whether it might be a more trivial consequence of spectral correlations due to the non sinusoidal waveform constituting the recorded time series [1, 20, 35]. The apparent PAC that arises from non sinusoidal dynamics with high harmonic content has been hypothesized to be informative about the underlying neural processes [7], and it was experimentally demonstrated for interacting non linear acoustic oscillators in [36]. However, the interpretation of the PAC phenomenon is completely different according to the mechanism that generates it. For instance, in [11] we demonstrated, through a harmonicity-PAC analysis using the TLI metric, that harmonic and non harmonic PAC patterns coexist during the seizure dynamics recorded with intracerebral macroelectrodes in epilepsy patients. We found that harmonic and non harmonic PAC patterns observed during the ictal activity can be interpreted as emerging features linked to the restrained and paroxysmal depolarizing shifts, which constitutes two essentially different neural mechanisms of seizure propagation. Importantly, the capability of the TLI metric to quantitatively distinguish the non harmonic PAC pattern, is clinically relevant since this specific pattern has been previously associated with the ictal core through the paroxysmal depolarizing shifts mechanism of seizure propagation.

The evidence discussed above highlights the relevance to unravel the complex interplay between spectral harmonicity and different types of CFC. Several approaches and controls have been previously proposed to address the true/spurious dichotomy in connection with PAC. In [37] it was argued that an increase in PAC intensity associated with a decrease in power of the modulating LF component would be an indication of the existence of ‘true’ coupling. Conversely, the presence of concomitant AAC and PAC patterns could be a proxy for ‘spurious’ PAC, in which the thigh correlation between the amplitude of the putative modulating LF and the modulated HF rhythms giving rise the AAC pattern, are produced by harmonically related spectral components constituting an underlying non sinusoidal oscillatory dynamics [38]. Another approach suggested in [37] refers to the use of multimodal recordings (e.g. LFP, single and multi unit activity). Specifically, analysis of spike-triggered LFP recordings can be used to confirm that spike timing is clocked by the phase of ongoing HF component (e.g. gamma oscillations), hence, revealing that the modulated fast rhythm is not an HF harmonic of the slow modulating rhythm but associated to genuine HF oscillatory activity. Multi site recordings allow measures of inter area PAC in which slow and fast rhythms are extracted from time series recorded in different neural populations. Importantly, measures of inter area PAC in which the slow and fast rhythms are generated in distinct oscillators reduce concerns on spurious coupling [37, 39]. In this regard, it has been noted that one-to-one mapping between electrode measurement (i.e. time series) and neural source of oscillations (e.g. LFP) does not hold in real data, as there are multiple neural networks that generate fields measured by a single electrode [17, 40]. Thus, the electrode time series is the result of a mix that could have very non sinusoidal waveform shape that is not present in any of the individual sources [17, 40]. Multichannel recordings in combination with tools for CFC source identification have been proposed as a way to disambiguates this issue [17, 40, 39].

Here we noted that in all these previous works it has been implicitly assumed that ‘spurious’ CFC patterns are intrinsically linked to an underlying non sinusoidal oscillatory dynamics characterized by a high harmonic content in its power spectrum. However, our results suggest that this assumption does not hold in realistic scenarios. In Velarde et al. [1] we analytically demonstrated that PAC phenomenon naturally emerges in mean-field models of biologically plausible networks, as a signature of specific bifurcation structures. Importantly, Velarde et al. [1] found that the mechanisms producing ‘true’ PAC (i.e. secondary Hopf bifurcation and PEI mechanism), in general elicit two coupled non sinusoidal oscillatory dynamics with independent fundamental frequencies. These results suggest that the resulting oscillatory dynamics underlying ‘true’ PAC is in general characterized by a high harmonic content in its power spectrum. In this work we quantitatively analyzed the role of the spectral harmonicity in different types of CFC patterns not restricted only to PAC and thus, providing a broader vision on this open issue in comparison to that addressed in previous reports. The results obtained using biologically plausible neural network models and more generic non linear and parametric oscillators reveal that harmonicity-CFC interplay is more complex than previously thought.

In line with the discussion given above about co-occurring AAC and PAC pa-terns [37, 38], we found that special care should be taken to interpret CFC patterns involving the same properties in the LF and HF frequency bands (e.g. PPC, AAC, FFC) since they might be epiphenomenal patterns elicited by a single non sinusoidal oscillatory dynamics constituted by harmonically related frequency components, in which the harmonic components within the HF band follows the changes of the fundamental frequency component in the LF band (see Section 3.2). However, in Sections 3.3 and 3.4 we show that concomitant PAC and PFC patterns were related to the presence of ‘true’ PFC with high harmonic content via the rectification mechanisms elicited by the PAC pattern. As a conclusion, the co-occurrence of multiple CFC patterns should not be taken as a straightforward indicator of spurious coupling per se. In Section 3.2 we show that a single oscillatory dynamics characterized by a non constant oscillation period can produce ‘spurious’ CFC with low harmonic content (i.e. non harmonic CFC). This type of oscillatory dynamics is commonly observed in oscillators undergoing a chaotic regime or non linear oscillators under the effect of intrinsic noise (Figure 15H). On the other hand, in Sections 3.3 and 3.4 we show that two coupled oscillatory dynamics with independent fundamental frequencies can elicit ‘true’ CFC with high harmonic content via rectification mechanisms (or other post-interaction nonlinear processing mechanisms). In Table 3 we resume the evidence supporting the conclusion that ‘true’ and ‘spurious’ concepts applied to the CFC patterns are not intrinsically linked to the harmonic content of the underlying oscillatory dynamics. Based on this results, we claim that the high harmonic content observed in a given oscillatory dynamics is neither sufficient nor necessary condition to interpret the associated CFC pattern as ‘spurious’ or epiphenomenal, i.e. not representing a true interaction between two coupled oscillatory dynamics with independent fundamental frequencies.

**Table 3:** Summary of the harmonic and nonharmonic cross-frequency couplings observed in simulated and experimental oscillatory dynamics.

|  |  | Nature of the CFC |  |
| --- | --- | --- | --- |
|  |  | True CFC* | Spurious CFC** |
| Harmonicity | Non harmonic | **PAC:**<br>- PEI mechanism without phase-resetting (Figure B.12I,J,N,O).<br>- Secondary Hopf bifurcation [1].<br>- Epileptiform local field potentials, e.g. spike-wave discharges associated to the paroxysmal depolarizing shifts [11].<br>**PFC:**<br>- Forced parametric oscillator with additive noise (Figures 17K,L, B.10K,L and B.11K,L). | **PAC:**<br>- Non linear oscillator with intrinsic noise (Figure 15K,L).<br>**AAC:**<br>- Forced non linear oscillator with intrinsic noise (Figures 16D,E).<br>**FFC:**<br>- Forced non linear oscillator with intrinsic noise (Figures 16I,J). |
|  | Harmonic | **PAC:**<br>- PEI mechanism with phase-resetting (Figure 18D,E and [1]).<br>- Epileptiform local field potentials, e.g. non sinusoidal repetitive wave-form shapes associated to restrained depolarising shifts [11].<br>**PFC:**<br>- Forced parametric oscillator with additive noise (Figures 17D,E,F,G, B.10D,E,F,G and B.11D,E,F,G).<br>**AAC:**<br>- Coupled non linear acoustic oscillators [36]. | **PAC:**<br>- Non linear oscillator (Figure 15D,E,I,J, [1] and [36]).<br>**AAC:**<br>- Forced non linear oscillator (Points in Figure 16C located in between panels A and D).<br>**FFC:**<br>- Forced non linear oscillator (Points in Figure 16H, located in between panels F and I). |
| * **True CFC:** Two (or more) coupled oscillatory dynamics characterized by independent fundamental frequencies.<br>** **Spurious CFC:** A single non sinusoidal oscillatory dynamics characterized by dependent, i.e. harmonically related, frequencies. |  |  |  |

## 5. CONCLUSION

We found that harmonic and non harmonic patterns associated to a variety of CFC types (e.g. PAC, PFC) naturally emerges in the dynamics characterizing biologically plausible neural network models and more generic non linear and parametric oscillators. Substantial evidence was presented supporting the conclusion that ‘true’ and ‘spurious’ concepts applied to the CFC patterns are not intrinsically linked to the harmonic content of the underlying oscillatory dynamics. More specifically, the high harmonic content observed in a given oscillatory dynamics is neither sufficient nor necessary condition to interpret the associated CFC pattern as ‘spurious’ or epiphenomenal, i.e. not representing a true interaction between two coupled oscillatory dynamics with independent fundamental frequencies. In addition, the proposed signal processing techniques provide an extension of the traditional analytic toolkit used to quantify and interpret CFC patterns observed in oscillatory dynamics elicited by physical and biophysical systems. There is mounting evidence suggesting that the combination of multimodal recordings, specialized signal processing techniques and theoretical modeling is becoming a required step to completely understand CFC patterns observed in oscillatory rich dynamics of physical and biophysical systems.

## ACKNOWLEDGEMENTS

This work was partially supported by UNCuyo-SeCTyP 2019 (80020180100576UN, 80020180100653UN) and Consejo Nacional de Investigaciones Científicas y Tecnicas (CONICET) PIP 112 201301 00 256, Argentina.

## Appendix A. Supplementary methods

### Appendix A.1. Synthetic signals

Synthetic dynamics associated to the analysis of various PAC and PPC patterns were computed as follows,

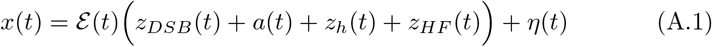

In Eq. A.1, 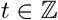 is the discrete time index, *z_DSB_*(*t*) is the amplitude modulated (double side band) signal with a sinusoidal carrier of frequency *f_HF_*, *a*(*t*) is the modulating signal, *z_h_*(*t*) is a sum of harmonic oscillations of the fundamental frequency *f_LF_, z_HF_*(*t*) is a sinusoidal component with frequency *f_HF_, η*(*t*) represent extrinsic (i.e. of observation) additive white Gaussian noise (AWGN). The amplitude envelope of the entire time series 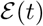 was included to emulate CFC and harmonicity transients in the synthetic dynamics and it was defined in terms of the sigmoid function,

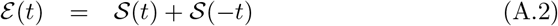

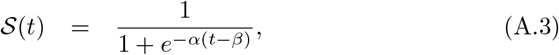

where *α* and *β* are parameters controlling the edge steepness and the time shift of the time series envelope, respectively. For synthetic oscillatory dynamics in permanent regime with no transients we use 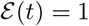. The amplitude modulated signal *z_DSB_*(*t*) was defined as,

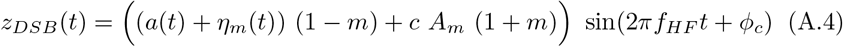

In Eq. A.4, *a*(*t*) defines the shape of the amplitude envelope of the sinusoidal carrier with frequency *f_HF_, A_m_* is the maximum value of the modulating *a*(*t*), *η_m_*(*t*) is additive white Gaussian noise (AWGN) intrinsic to the modulation process, *m* define the modulation depth (*m* = 0 imply maximum modulation depth and *m* = 1 for no modulation) and *c* is the carrier factor controlling the type of amplitude modulation (AM),

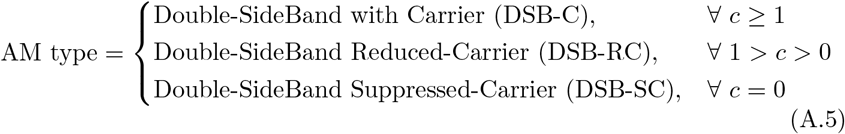

For the sake of consistency with modulation depth (*m*) and carrier factor (*c*) parameters in Eq. A.4, the modulating signal *a*(*t*) must satisfy the condition *min*(*a*(*t*)) = −*max*(*a*(*t*)). We explored two types of waveform shapes for the periodic modulating signals *a*(*t*). The sinusoidal modulating signal was defined as,

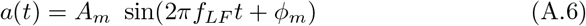

On the other hand, the periodic Gaussian modulating was defined as,

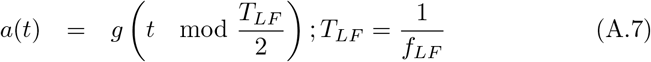

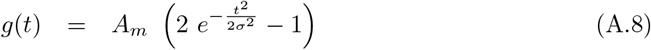

In Eq. A.8, *g*(*t*) define a single period with the shape of a Gaussian probability density function with standard deviation *σ*. In Eqs. A.7, *g*(*t*) is repeated with a period of *T_LF_* samples to obtain a periodic modulating signal *a*(*t*) with a Gaussian waveform shape.

The *z_h_*(*t*) and *z_HF_*(*t*) signals were defined as follows,

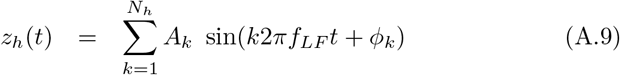

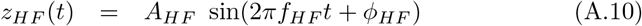

### Appendix A.2. Van der Pol oscillator

The Van der Pol oscillator is a non linear and time invariant system whose dynamics is defined by the following differential equation,

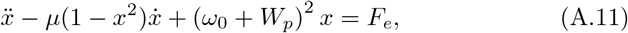

where the over-dot represents time derivative, *μ* is a scalar parameter controlling the nonlinearity, *ω*_0_ = 2*πf*_0_ is the angular frequency of oscillation when *μ* = 0, *W_p_* = 0 and *F_e_* = 0. The time variant parameter *W_p_* and the external driving *F_e_* were defined as,

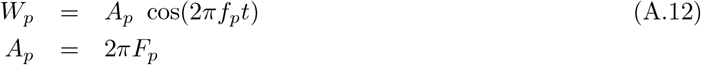

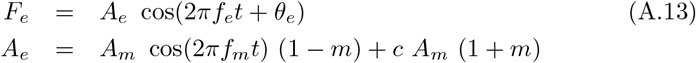

where *F_p_* and *f_p_* have units of Hz, *F_e_* is defined as an amplitude-modulated external driving with frequency *f_e_* and constant phase *θ_e_*, being *f_m_* the frequency of the sinusoidal modulating, *A_m_* the maximum value of the modulating, *m* the modulation depth and *c* the carrier factor controlling the type of amplitude modulation (see Eq. A.5). In presence of noise, the dynamics of the Van der Pol oscillator is described by the following system of stochastic differential equations,

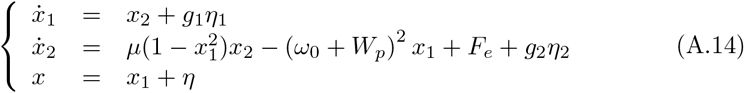

In Eq. A.14, *η*_1_ and *η*_2_ are independent and identically distributed random variables representing intrinsic noise, and *η* represent extrinsic (i.e. of observation) noise. For the simulations computed with the Eq. A.14, we use independent and normally distributed random variables for both intrinsic (*η*_1_, *η*_2_) and extrinsic (*η*) noise (i.e. white Gaussian noise). Therefore, the intrinsic noise components in Eq. A.14 result 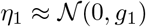 and 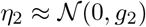, where *g*_1_ and *g*_2_ represent the standard deviation of the zero-mean normal distribution 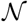. Unless otherwise specified, we use Additive White Gaussian Noise (AWGN), that is, the parameters defining the noise intensity *g*_1_ and *g*_2_ do not depend on the state variables *x*_1_ and *x*_2_. In the case of *g*_1_ = *g*_2_ = 0, Eqs. A.14 and A.11 are equivalent. In addition, we fixed *f*_0_ = 10 Hz and *θ_e_* = 0. Regarding the numerical integration of the stochastic differential equation A.14 we use an explicit solver based on the Euler-Heun method [41].

### Appendix A.3. Parametric oscil lator

To analyse the PFC patterns we use a linear and time variant system based on a 2nd order parametric oscillator whose dynamics is defined by the following differential equation,

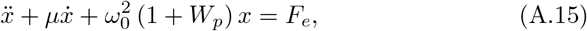

where the over-dot represents time derivative, *μ* is the parameter defining the intensity of the dissipative term, *ω*_0_ = 2*πf*_0_ is the angular frequency of oscillation when *μ* = 0, *W_p_* = 0 and *F_e_* = 0. The time variant parameter *W_p_* and the external driving *F_e_* were defined as,

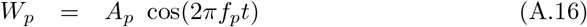

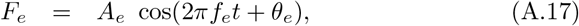

where *f_p_* and *f_e_* have units of Hz, *θ_e_* is a constant phase in rads., the parameters *A_p_* and *A_e_* defines the intensity of parametric and external driving, respectively. In presence of noise, the dynamics of the parametric oscillator is described by the following system of stochastic differential equations,

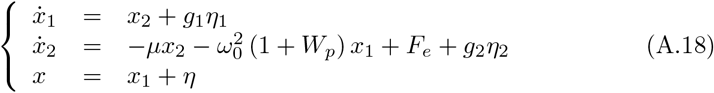

In Eq. A.18, *η*_1_ and *η*_2_ are independent and identically distributed random variables representing intrinsic noise, and *η* represent extrinsic (i.e. of observation) noise. For the simulations computed with the Eq. A.18, we use independent and normally distributed random variables for both intrinsic (*η*_1_, *η*_2_) and extrinsic (*η*) noise (i.e. white Gaussian noise). Therefore, the intrinsic noise components in Eq. A.18 result 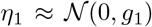 and 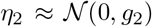, where *g*_1_ and *g*_2_ represent the standard deviation of the zero-mean normal distribution 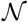. Unless otherwise specified, we use Additive White Gaussian Noise (AWGN), that is, the parameters defining the noise intensity *g*_1_ and *g*_2_ do not depend on the state variables *x*_1_ and *x*_2_. In the case of *g*_1_ = *g*_2_ = 0, Eqs. A.18 and A.15 are equivalent. In addition, we fixed *f*_0_ = 100 Hz, *μ* = 200, *f_p_* = *f_e_*. Regarding the numerical integration of the stochastic differential equation A.18 we use an explicit solver based on the Euler-Heun method [41].

### Appendix A.4. Cross frequency coupling metrics

Eqs. A.19 to A.24 show the proper configuration of the *y_HF_*(*t*), *ϕ_HF_*(*t*) and *ϕ_LF_*(*t*) time series to quantify PPC, PAC, AAC, PFC, AFC and FFC by means of the PLV, MVL and KLMI metrics using Eqs. 4 to 7.

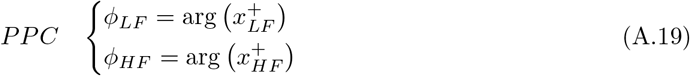

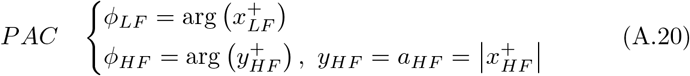

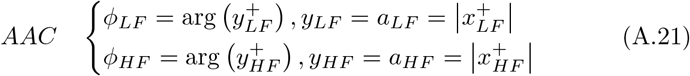

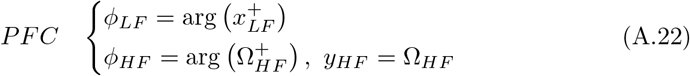

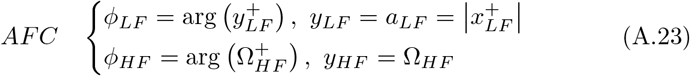

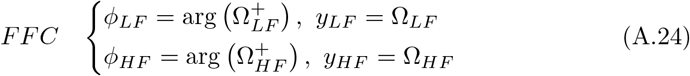

The time series configuration to assess PC using Eq. 8 is given by Eq. A.25.

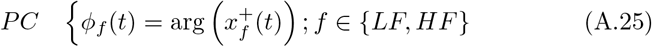

The *y_HF_*(*t*), *ϕ_HF_*(*t*) and *ϕ_LF_*(*t*) time series required to assess the six types of CFC were computed using the Filter-Hilbert method (see Chapter 14 in [17]). In brief, the raw time series *x*(*t*) was band-pass filtered around the frequency band of interest *f* ∈ {*LF,HF*}, then, the analytic signal 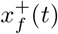 corresponding to the filtered time series *x_f_*(*t*) was computed in the frequency domain using the following equations [36, 42],

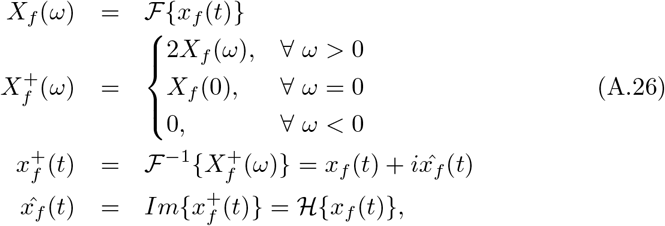

In Eq. A.26, 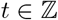 is the discrete time index, *ω* is the non dimensional angular frequency (see Appendix A.6), *X_f_*(*ω*) is the discrete Fourier transform, *i* is the imaginary unit, 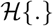 denotes the Hilbert transform, 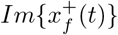 stands for the imaginary part of 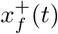 and the operators 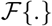 and 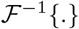 denote the discrete Fourier transformation and its inverse respectively, which were computed via the fast Fourier transform algorithm.

The amplitude envelope *a_f_*(*t*) and phase *ϕ_f_*(*t*) time series for that particular frequency band *f* ∈ {*LF, HF*} were obtained by computing the absolute value and argument of the analytic signal 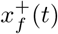, respectively:

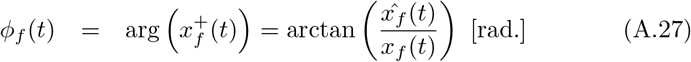

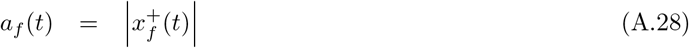

In Eqs. A.22, A.23 and A.24, the instantaneous frequency time series *Ω_LF_*(*t*) and *Ω_HF_*(*t*) were computed following the procedures described in the Section 2.5.

### Appendix A.5. Band-pass filtering

The band-pass filters (BPF) involved in the computation of *x_f_*(*t*) from the raw time series *x*(*t*) were implemented in the frequency domain by multiplying the Fourier transform of the input signal by a Hann window and then, applying the inverse Fourier transform to get the band-pass filtered signal back in the time domain (i.e. circular convolution in the discrete time domain) [1, 11, 36]. Note that this filtering approach was used to effectively isolate the desired frequency bands (i.e. null-to-null bandwidth), which is not guaranteed when other linear filters are used (e.g. low order IIR filters) [31]. We verified that our BPF implementation do not produce neither phase distortions nor significant artificial oscillations in the output signal capable to generate spurious CFC [32], showing a performance comparable to that of the FIR filters implemented in the EEGLAB (eegfilt function, data not shown) [43]. In order to mitigate edge artifacts due to the transient response of the BPFs and the computation of the analytic signals, we implemented the time series reflection procedure described in [17]. Briefly, time series are reversed in time, concatenated to both ends of the real-data time series, analyses were performed, and then, the reflected portion of the data were trimmed.

### Appendix A.6. Fourier transform of the discrete time derivator

In this section we shall obtain the expression for the discrete Fourier transform of the discrete time derivator defined as,

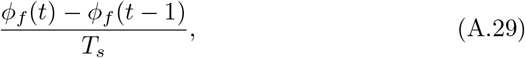

where *T_s_* = 1/*f_s_* is the sampling time interval corresponding to the sampling rate *f_s_*. Taking into account the analysis and synthesis equations of the discrete time Fourier transform,

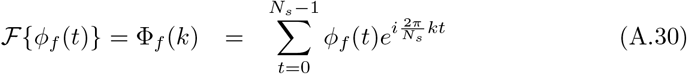

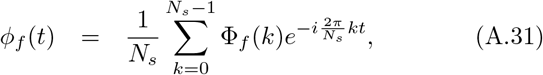

where *i* is the imaginary unit, *t*, 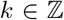 are the discrete time and frequency indices, respectively and *N_s_* is the number of samples of the time series. Applying Eq. A.30 to Eq. A.29 and introducing the non dimensional angular frequency *ω* = *k* 2*π/N_s_*, we obtain,

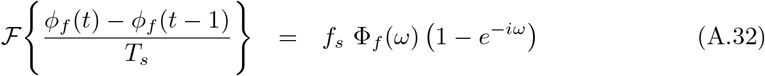

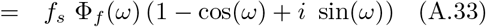

where we have applied the time shifting property of the Fourier transform (in this case for a time shift *t* = −1). The Eq. A.32 is the discrete Fourier transform of the discrete time derivator in Eq. A.29.

Let us now consider that the oversampling condition given by *f_s_* ≫ *f*: *f* ∈ {*LF, HF*} is satisfied. As a consequence, in the discrete frequency domain this condition implies *k* ≪ *N_s_*, or equivalently, *ω* ≈ 0. Under this condition, the Eq. A.33 can be well described by a first order approximation in the non dimensional angular frequency *ω* which can be written as,

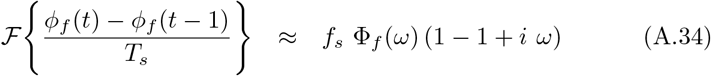

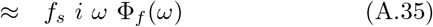

## Appendix B. Supplementary results

### Appendix B.1. Bias of the TLI

Figure B.1 shows that the TLI and *PLV_PPC_* metrics present a comparable bias when computed on non harmonically related oscillations. Figure B.1 shows that the bias of the TLI and *PLV_PPC_* metrics rapidly increases for epoch lengths shorter that ≈ 10 cycles of the slow rhythm, being this bias rather independent of the noise level (AWGN) and the non harmonic ratio (*R* = *f_HF_/f_LF_*) between the slow and fast oscillations.

**Figure B1:**
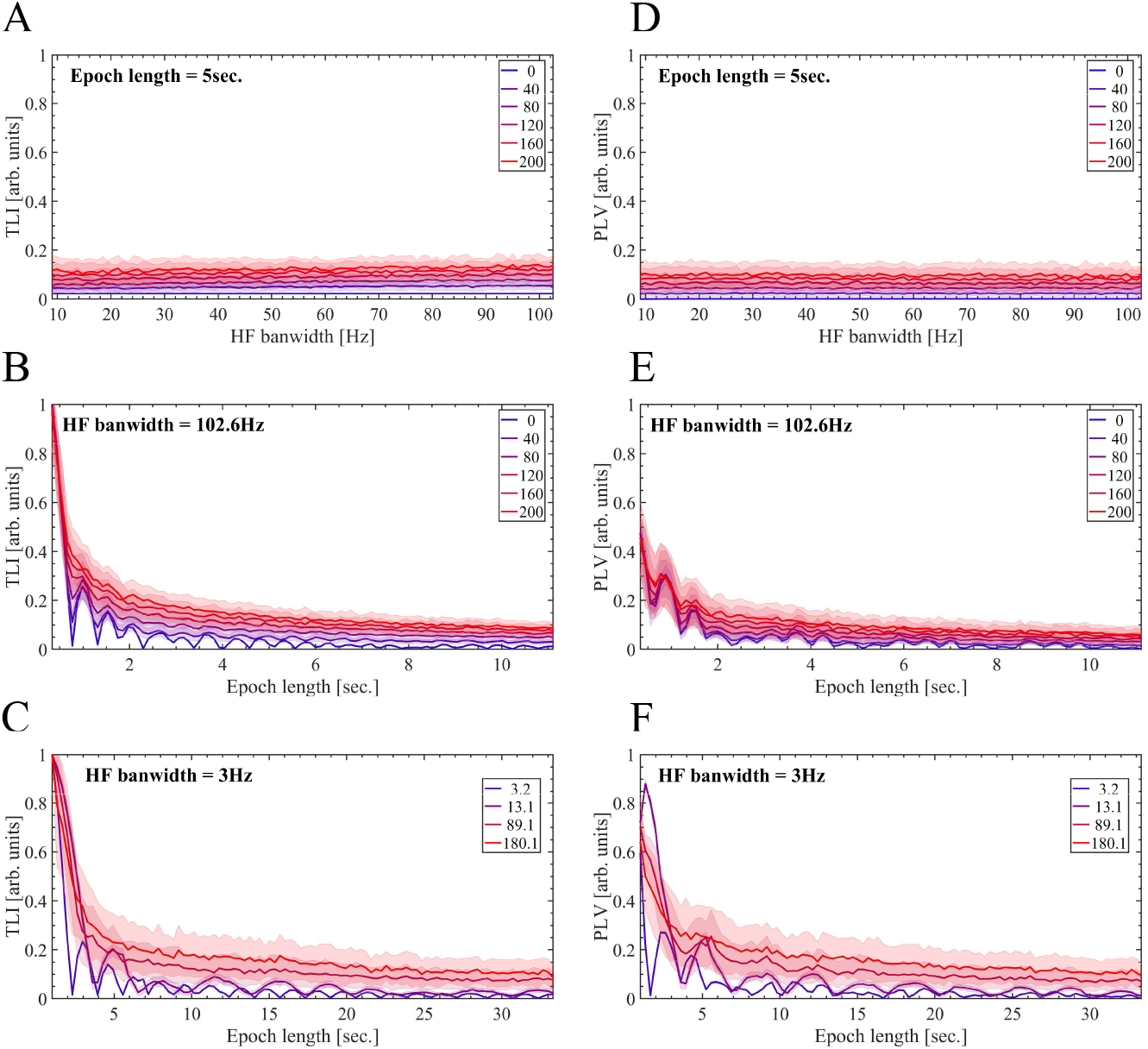
The TLI and *PLV_PPC_* metrics present a comparable bias when computed on non harmonically related oscillations. In all the cases shown in this figure, we used a sampling rate of *f_s_* = 2000 Hz and the bandwidth of the BPF for the LF component (LF BPF) was kept fixed at *Bw_LF_* = *f_LF_*. We use the BPF as described in Appendix A.5. Besides, in all the cases shown in this figure the noise level is expressed as the percent of the amplitude of the LF component at *f_LF_* Hz scaling the standard deviation *σ* of the additive white Gaussian noise 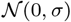. Panels A, B, D and E, were computed using a synthetic dynamics similar to that used in Figure 5, but in this case it is constituted by two non harmonic oscillations at *f_LF_* = 9 Hz and *f_HF_* = 7.2 *f_LF_* = 64.8 Hz. For panels A, B, D and E, the *PLV_PPC_* was computed using Eq. 4 with the configuration given by Eq. A.19 and *M* = 1, *N* = 7. (A, D) TLI and *PLV_PPC_* metrics as a function of the HF bandwidth (*Bw_HF_*) corresponding to the filter HF BPF used to obtain the HF signal (*x_HF_*(*t*)), and taking the level AWGN as a parameter. The minimum and maximum *Bw_HF_* values used to compute the graphs B and E were 9 Hz and 102.6 Hz, respectively. To compute these graphs, the epoch length was kept unchanged in 5 sec. (B, E) TLI and P LVPPC metrics as a function of the epoch length and taking the level of additive white Gaussian noise (AWGN) as a parameter. To compute graphs B and E, the bandwidth of the filter HF BPF was kept unchanged at *Bw_HF_* = 102.6 Hz. Our implementation of the TLI algorithm (Section 2.4) requires at least 3 cycles of the low frequency oscillation (*f_LF_* = 9 Hz), which determines the minimum epoch length shown in graphs A and D (3/*f_LF_* ≈ 0.3 sec.). The maximum epoch length used to compute graphs A and D was 100/*f_LF_* ≈ 11.1 sec. (C, F) The TLI and P LVPPC metrics as a function of the epoch length and taking the non harmonic ratio *R* = *f_HF_/f_LF_* as a parameter. Panels C and F were computed using a synthetic dynamics similar to that used in Figure 5, but in this case it is constituted by two non harmonic oscillations at *f_LF_* = 3 Hz and *f_HF_* = *R* × *f_LF_* with *R* = 3.2, 13.1, 89.1, 180.1. To compute graphs C and F, the bandwidth of the filter HF BPF was kept unchanged at *Bw_HF_* = *Bw_LF_* = *f_LF_* = 3 Hz. The noise level was set to 20 percent of the amplitude of the LF component at *f_LF_* = 3 Hz. The minimum and maximum epoch length shown in graphs C and F are 3/*f_LF_* ≈ 1 sec. and 100/*f_LF_* ≈ 33.3 sec., respectively. In all the panels, the solid lines represent the mean values and the shaded error bars correspond to the standard deviation of 100 realizations at each point.

### Appendix B.2. CFC time series and the bias produced by phase clustering

Figures B.2 and B.3 show the temporal evolution of the PAC (*PLV_PAC_*), harmonicity (TLI) and phase clustering (*PC_LF_*) metrics for synthetic dynamics presenting non PAC and a transient pattern of non harmonic PAC, respectively. Figures B.2 and B.3 should be compared with the results for a synthetic dynamics presenting a transient pattern of harmonic PAC (Figure 9).

**Figure B2:**
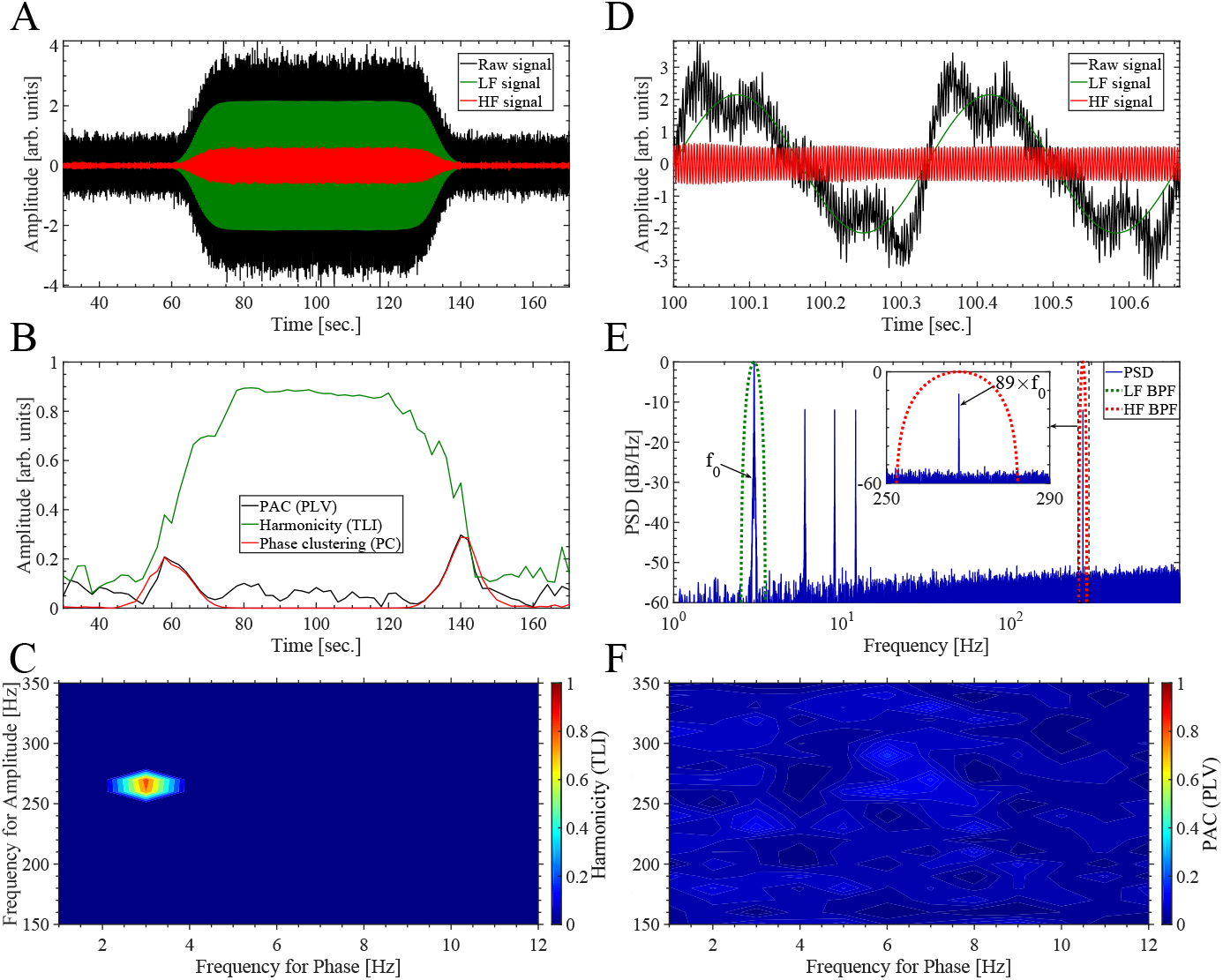
Temporal evolution of the PAC (*PLV_PAC_*), harmonicity (TLI) and phase clustering (*PC_LF_*) metrics during a synthetic dynamics without PAC. To obtain all the band-pass filtered signals shown in this figure we use the BPF as described in Appendix A.5. (A) Synthetic dynamics (solid black line) together with the HF and LF signals shown as solid red and green lines, respectively. The dynamics (solid black line) was synthesized using Eqs. A.1 and A.4 with the following hyperpameter values: sampling rate *f_s_* = 2000 Hz, *c* = 1 (i.e. DSB-C), zero modulation depth *m* = 0, *η_m_* = 0, we used a sinusoidal modulating *a*(*t*) with the fundamental frequency at *f*_0_ = *f_LF_* = 3 Hz as given by Eq. A.6 with *A_m_* = 1, *z_DBS_* was set with *f_HF_* = 89 × *f_LF_* = 267 Hz, *ϕ_c_* = 0, *z_HF_* = 0, for *z_h_* we use *A*_1_ = 4, *A_k_* = 1 ∀ 2 ≤ *k* ≤ 4, *A_k_* = 0 ∀*k* ≥ 5 and *ϕ_k_* = 0 ∀*k*. The transient pattern was implemented through the time series envelope 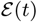 as defined in Eqs. A.2 and A.3, with *α* = 0.5 and *β* equals to one third of the time series length. Extrinsic noise *η* (*t*) was added as shown in Eq. A.1. In this case the noise level corresponds to the 10 percent of the maximum amplitude of the deterministic part of signal *x*(*t*) (i.e first term of the right-hand member of the Eq. A.1), scaling the standard deviation *σ* of the additive white Gaussian noise (AWGN) 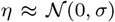. The LF (solid green line) and HF (solid red line) signals where obtained by filtering the raw signal (solid black line) with the band-pass filters whose power responses are shown as dotted green (*Bw_LF_* = 1 Hz) and red (*Bw_HF_* = 30 Hz) lines in graph E, respectively. (B) Time series showing the temporal evolution of the *PLV_PAC_*, TLI and *PC_LF_* metrics. These time series were computed as described in Section 2.8 using the algorithm 2 summarized in Table 2, with a sliding window of 20 sec. in length, i.e. 60 cycles of the slowest oscillatory component at *f*_0_ = *f_LF_* = 3 Hz. (C) TLI harmonicity map computed as described in Section 2.7 using a 20 sec. epoch extracted from the center (*Time* ≈ 100 sec.) of the synthetic dynamics shown in panel A. In computing the map, all the TLI values below the significance threshold were set to zero (see Section 2.7). The pseudocolor scale represents the TLI values ranging from 0 (blue) to 1 (red). (D) Zoom showing two cycles of the synthetic dynamics (solid black line) together with the HF and LF signals shown as solid red and green lines, respectively. The two cycle epoch corresponds to the center (*Time* ≈ 100 sec.) of the synthetic dynamics shown in panel A. (E) Power spectrum (solid blue line) computed from the synthetic dynamics (solid black line in graph A). The power responses (i.e. square magnitude) of the BPF used to compute the LF and HF signals are shown as dotted green and red lines, respectively. (F) Comodulogram computed as described in Section 2.7 computed from the same epoch used to obtain the harmonicity map (panel C). In computing the comodulogram, all the |*PLV_PAC_*| values below the significance threshold were set to zero (see Section 2.7). The pseudocolor scale represents the |*PLV_PAC_*| values ranging from 0 (blue) to 1 (red). The harmonicity map (panel C) and comodulogram (panel F) were computed using the same BPF (see Appendix A.5).

**Figure B3:**
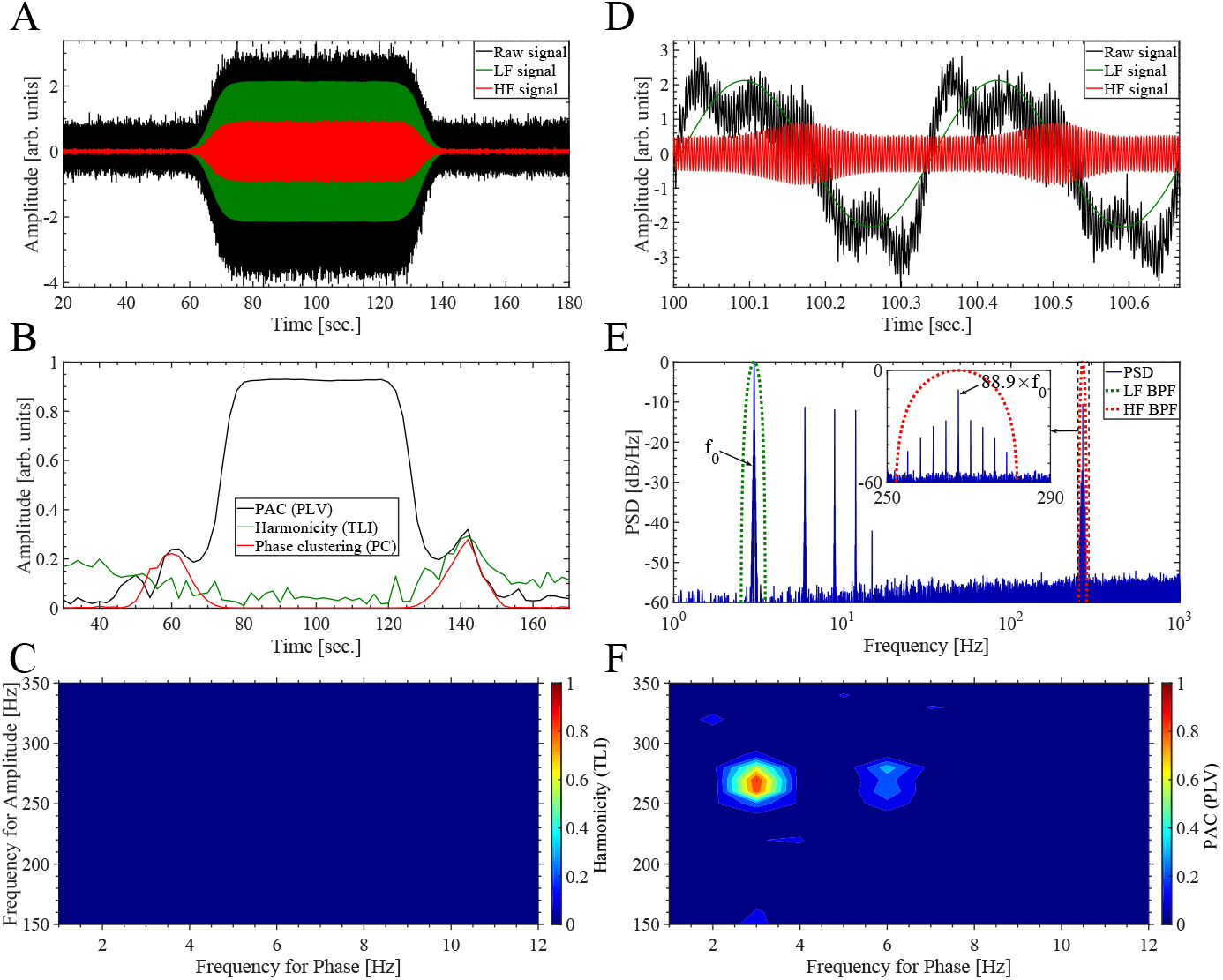
Temporal evolution of the PAC (*PLV_PAC_*), harmonicity (TLI) and phase clustering (*PC_LF_*) metrics during a synthetic dynamics presenting a transient non harmonic PAC pattern. The synthetic dynamics was synthesized using the same parameter values than those used in Figure 9, except for the frequency of the carrier in the *z_DBS_* signal which in this case was set to *f_HF_* = 88.9 × *f_LF_* = 88.9 × 3 = 266.7 Hz. The *PLV_PAC_*, TLI and *PC_LF_* metrics were computed using the same set of hyperparameter values than those used in Figure 9. The description of the panels is the same than that given in Figure 9.

Figures 9 and 10 in the main text show that the presence of phase clustering (*PC_LF_*) produces a bias which reduces the magnitude of the PAC metric (*PLV_PAC_*) in presence of a harmonic PAC pattern. On the other hand, Figures B.4 and B.5 illustrate the complementary situation in which the magnitude of the PAC metric (*MVL_PAC_*) in absence of PAC is biased from closed to zero (see Figure B.5B) toward higher magnitude values (|*MVL_PAC_*| ≈ 0.6 in Figure B.5B), as a consequence of the presence of phase clustering (*PC_LF_*). In Figures B.4 and B.5 is also shown that the presence phase clustering (*PC_LF_*) introduces a bias that reduces the magnitude of the harmonicity metric (TLI) in presence of harmonically related oscillations (*f*_0_ = *f_LF_* = 3 Hz and *f_HF_* = 89 × *f_LF_* = 267 Hz).

**Figure B4:**
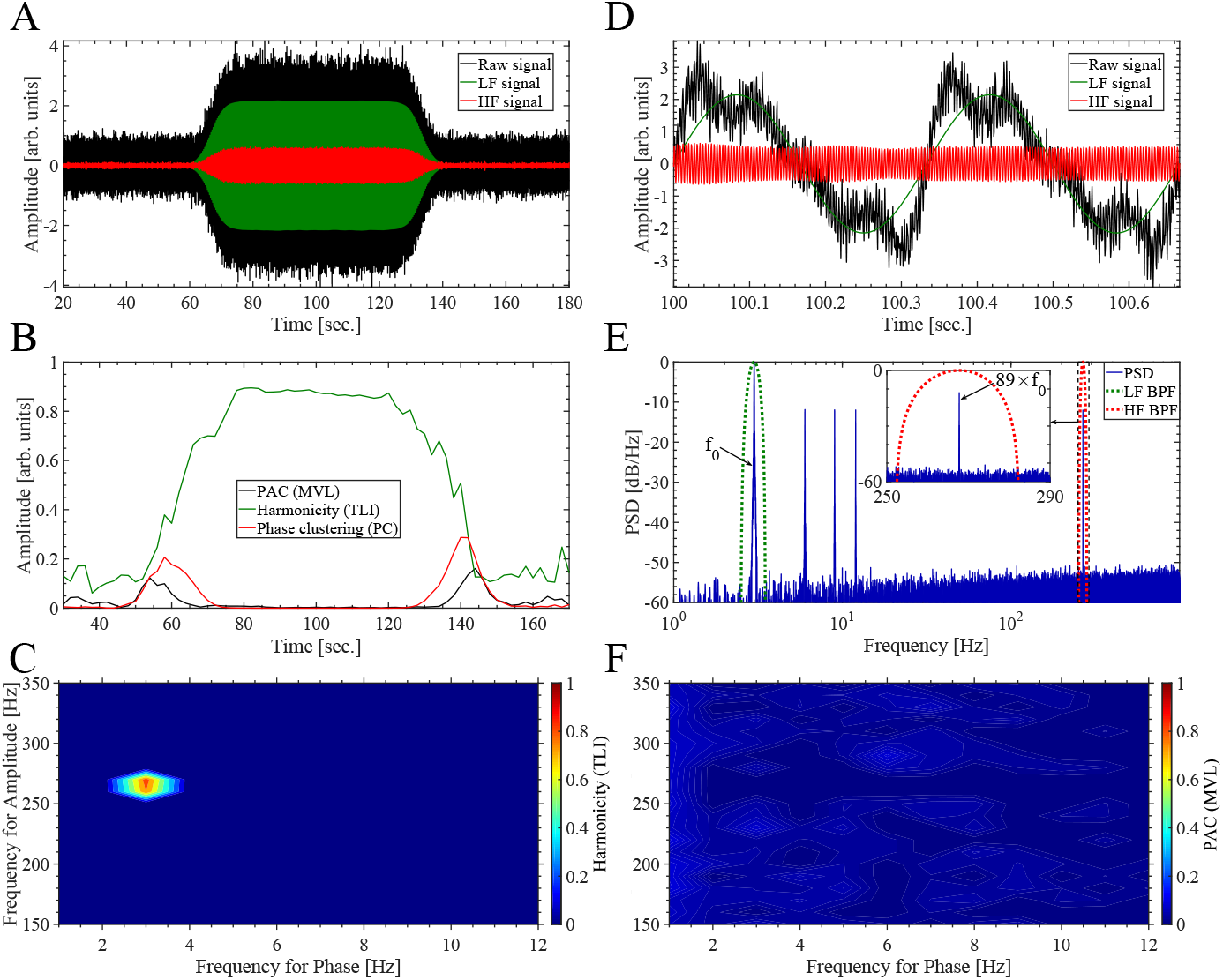
Temporal evolution of the PAC (*MVL_PAC_*), harmonicity (TLI) and phase clustering (*PC_LF_*) metrics during a synthetic oscillatory dynamics constituted by harmonically related rhythms with no PAC. The synthetic dynamics was synthesized using the same parameter values than those used in Figure B.2. The *MVL_PAC_* (see Eq. 5), TLI and *PC_LF_* metrics were computed using the same set of band pass-filters and hyperparameter values than those used in Figure B.2. The description of the panels is the same than that given in Figure B.2.

**Figure B5:**
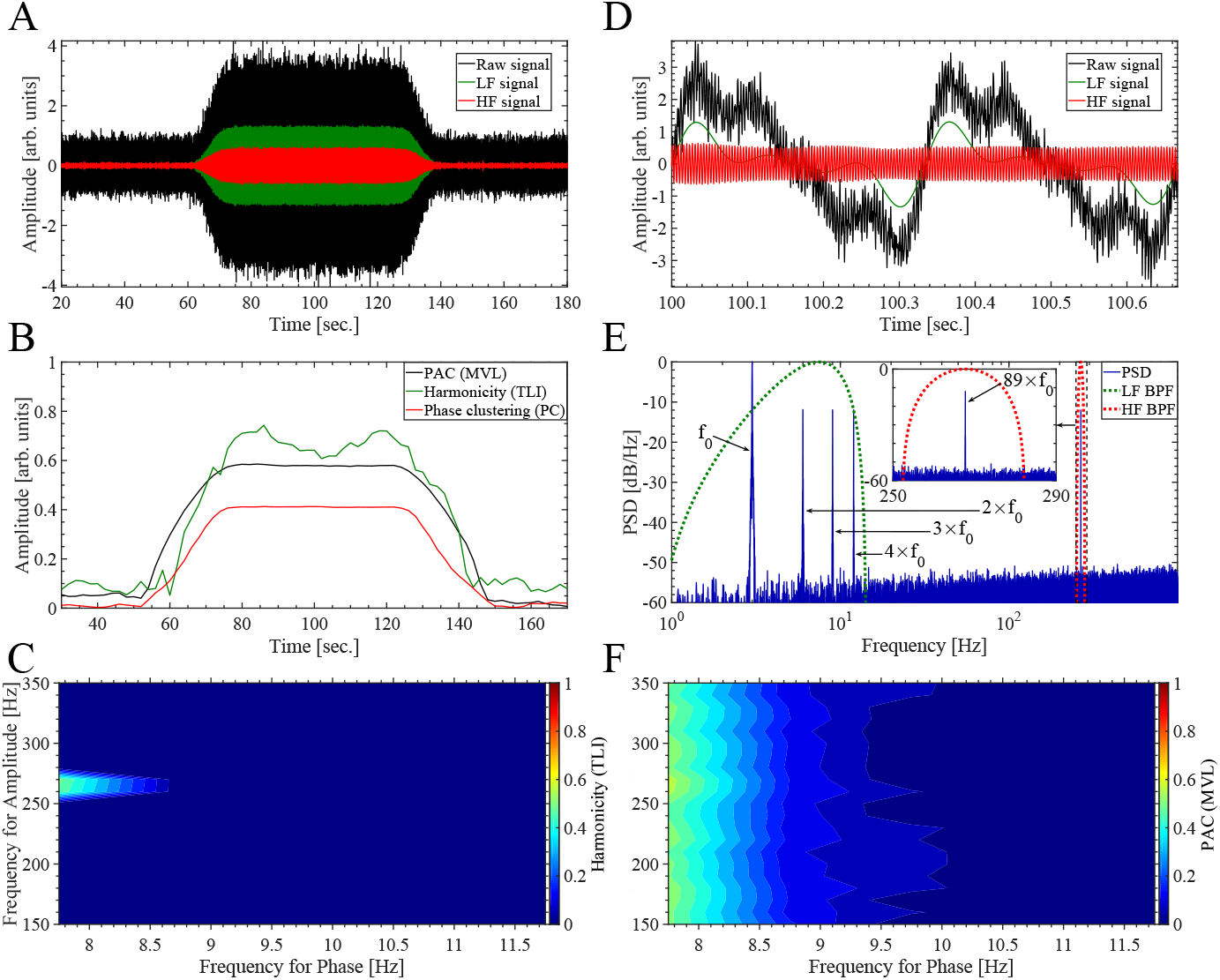
Temporal evolution of the PAC (*MVL_PAC_*), harmonicity (TLI) and phase clustering (*PC_LF_*) metrics during a synthetic oscillatory dynamics constituted by harmonically related rhythms with no PAC. In this plot we use the same synthetic dynamics and the same set of hyperparameter values to compute the metrics than those described in the caption of Figure B.2, except for the bandwidth of the BPF used to compute the LF component (*Bw_LF_*). In this case, the *MVL_PAC_* (see Eq. 5), TLI and *PC_LF_* metrics were computed using *Bw_LF_* = 13.5 Hz centered around 7.5 Hz (see the dotted green line in panel E). This wide BPF produces a non sinusoidal LF component (see solid green line in panel D), characterized by a non uniform distribution of phase values producing the increase of the phase clustering (*PC_LF_*) during the dynamics (see solid red line in panel B). Note the bias in the PAC (*MVL_PAC_*) and harmonicity (TLI) metrics due to the presence of phase clustering (*PC_LF_*). The description of the panels is the same than that given in Figure B.2.

### Appendix B.3. A single oscillatory dynamics characterized by dependent frequencies

Figure B.6 shows the phase portraits for the simulated dynamics of the Van der Pol oscillator, complementing the results shown in Figure 15 of the main text.

**Figure B6:**
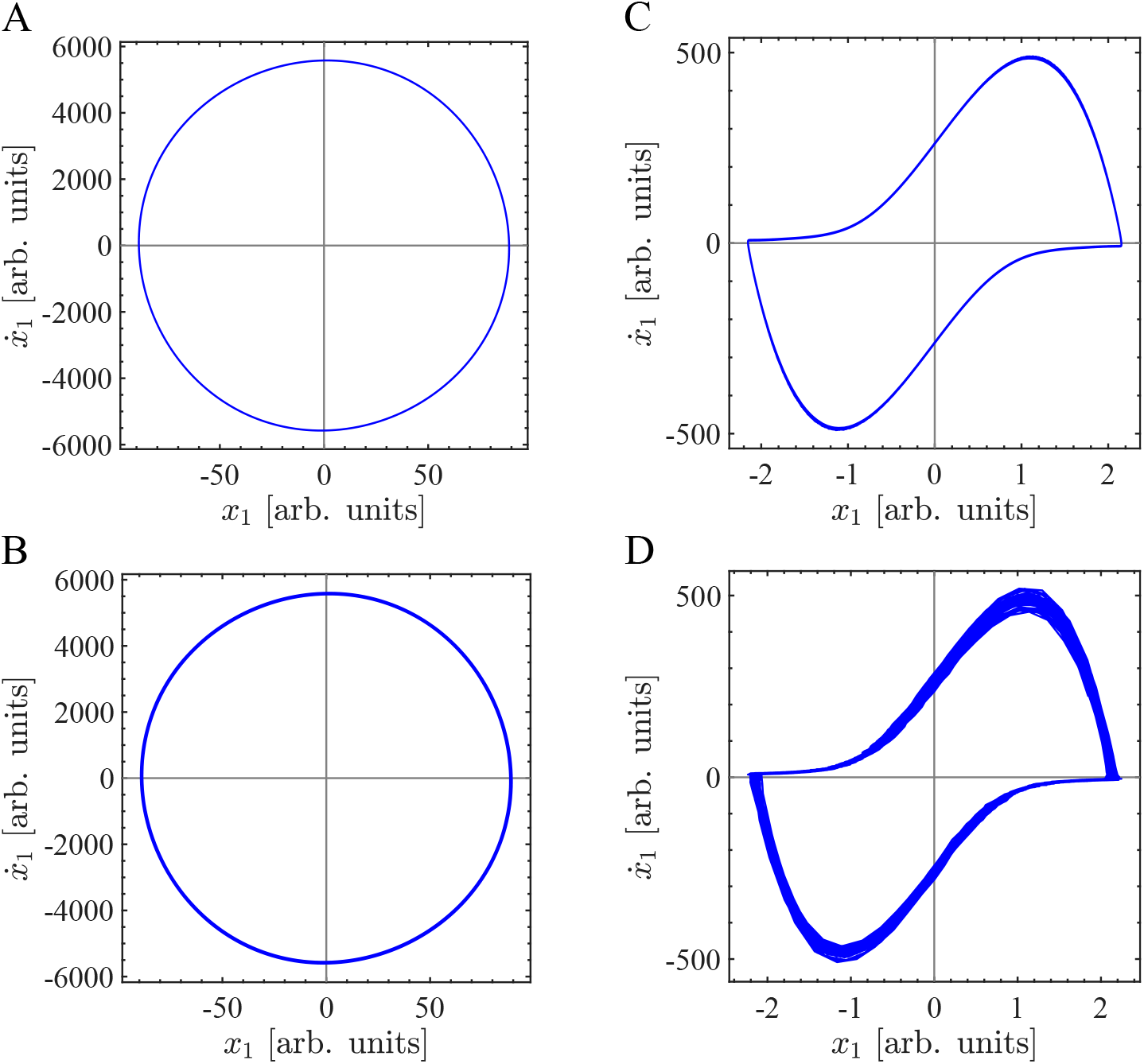
Phase portraits for the simulated dynamics of the Van der Pol oscillator. In this figure we use the same synthetic dynamics and the same set of hyperparameter values than those described in the caption of Figure 15. In particular, the phase portraits were computed using the dynamics *x*_1_ in Eq. A.14 which only takes into account the effect of the intrinsic noise, that is, without including the extrinsic (i.e. of observation) noise *η*. (A) Phase portrait corresponding to the dynamics shown in Figure 15A (No PAC). (B) Phase portrait corresponding to the dynamics shown in Figure 15F (No PAC). (C) Phase portrait corresponding to the dynamics shown in Figure 15D (Harmonic PAC). (D) Phase portrait corresponding to the dynamics shown in Figure 15K (Non harmonic PAC).

Figure B.7 shows the harmonicity-PAC plot usnig the TLI, P LVP AC and KLM IPAC metrics computed for the simulated dynamics of the Van der Pol oscillator with intrinsic noise of type non-additive white Gaussian noise (NAWGN).

**Figure B7:**
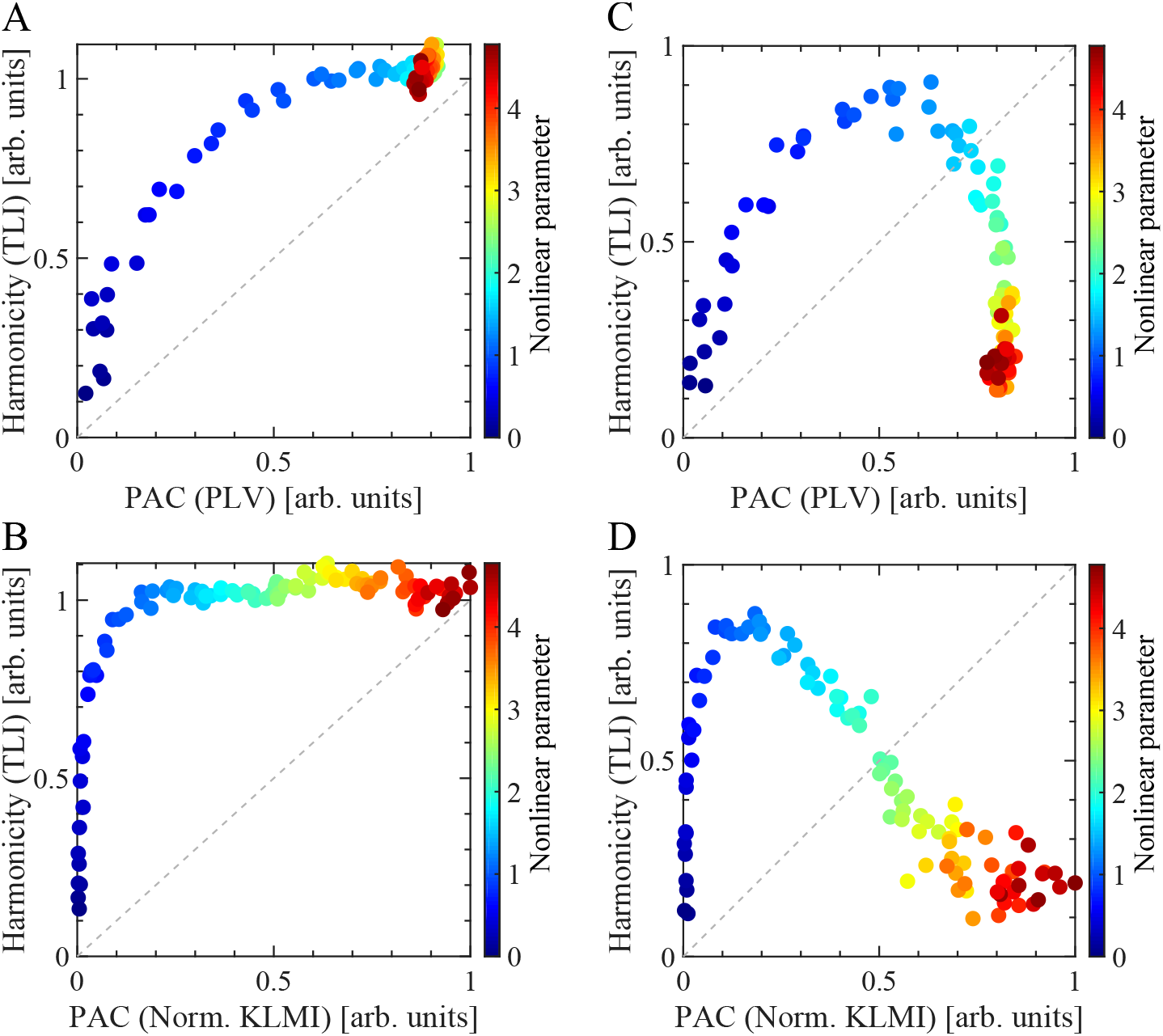
Harmonicity-PAC plot computed for the simulated dynamics of the Van der Pol oscillator with intrinsic noise of type non-additive white Gaussian noise (NAWGN). In this figure we use the same synthetic dynamics and the same set of hyperparameter values to compute the metrics than those described in the caption of Figure 15, except for the configuration of the intrinsic noise. In this case, we use non-additive white Gaussian noise (NAWGN). For the numerical integration of the stochastic differential equation A.14 we use an explicit solver based on the Euler-Heun method [41] using the Stratonovich integral formulation. Importantly, we verified that the harmonicity-PAC plots shown in this figure do not change when computed using the Itö integral formulation. For the panels A and B, the dynamics of the Van der Pol oscillator was simulated using intrinsic noise of type NAWGN applied only on the equation of *ẋ*_2_(*g*_1_ = 0 and *g*_2_ = 0.5*x*_2_ in Eq. A.14). For the panels C and D, the dynamics was simulated by applying the intrinsic noise of type non-additive white Gaussian noise (NAWGN) on the equations of both *ẋ*_1_ and *ẋ*_2_ (i.e. *g*_1_ = 0.5*x*_1_ and *g*_2_ = 0.5*x*_2_ in Eq. A.14). Therefore, in this case the intrinsic noise components (NAWGN) in Eq. A.14 result 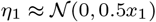 and 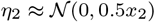. Extrinsic noise *η*(*t*) was added as shown in Eq. A.14. In this case the noise level corresponds to the 10 percent of the maximum amplitude of the dynamics *x*_1_ in Eq. A.14), scaling the standard deviation *σ* of the additive white Gaussian noise (AWGN) 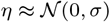. The harmonicity metric (TLI) was computed as it was described in Section 2.4. For the panels A and C, the PAC metric (*PLV_PAC_*) was computed using Eq. 4 with the configuration given by Eq. A.20 and *M* = *N* = 1. For the panels B and D, we compute the *KLMI_PAC_* using Eqs. 6 and 7 with the configuration given by Eq. A.20. Note that the *KLMI_PAC_* was normalized with its maximum value in each plot.

Figure B.8 shows the phase portraits for the simulated dynamics of the Vander Pol oscillator, complementing the results shown in Figure 16C of the main text.

**Figure B8:**
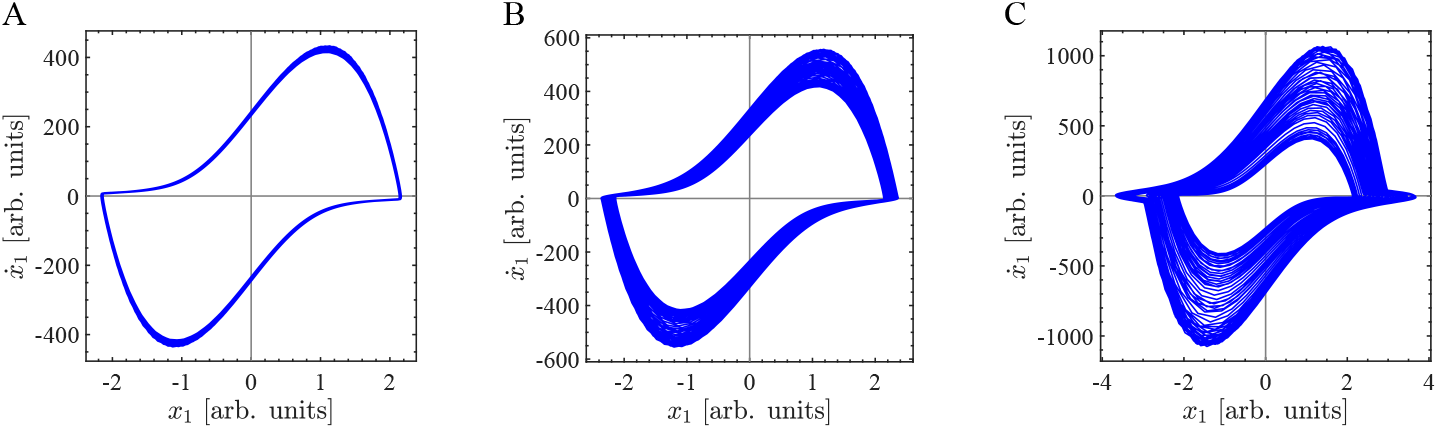
Phase portraits illustrating the simulated dynamics of the Van der Pol oscillator under an amplitude-modulated external driving. In this figure we use the same synthetic dynamics and the same set of hyperparameter values than those used to compute Figure 16C. In particular, the phase portraits were computed using the dynamics x1 in Eq. A.14 which only takes into account the effect of the intrinsic noise, that is, without including the extrinsic (i.e. of observation) noise *η*. (A) Phase portrait corresponding to the dynamics shown in Figure 16C for *A_e_*/(5 × 10^4^) ≈ 0.01. (B) Phase portrait corresponding to the dynamics shown in Figure 16C for *A_e_* /(5 × 10^4^) ≈ 0.1. (C) Phase portrait corresponding to the dynamics shown in Figure 16C for *A_e_* /(5 × 10^4^) ≈ 1.

### Appendix B.4. Two coupled oscillatory dynamics characterized by independent frequencies

Figure B.9 shows the phase portraits for the simulated dynamics of the 2nd order parametric oscillator, complementing the results shown in Figure 17 of the main text.

**Figure B9:**
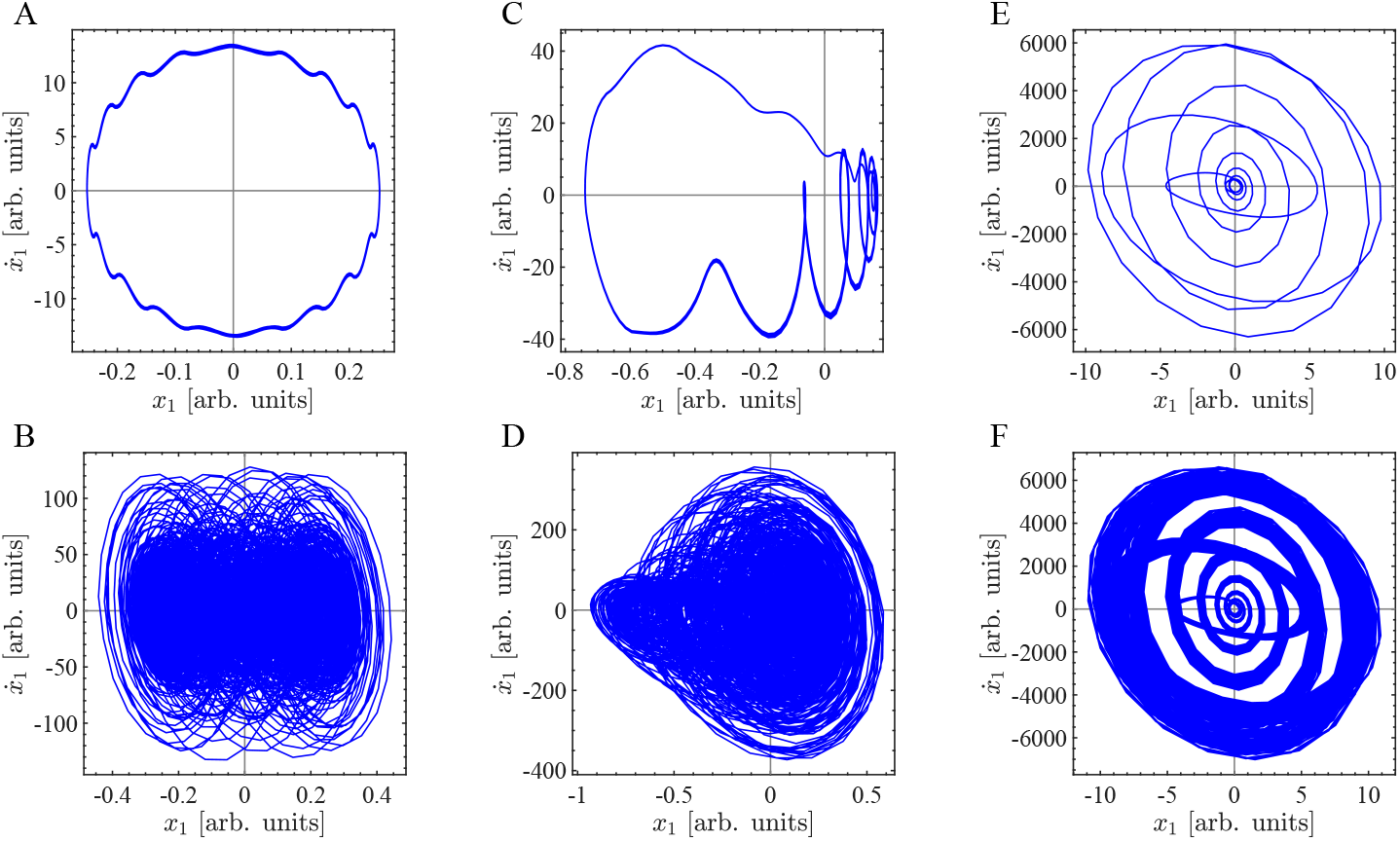
Phase portraits for the simulated dynamics of the 2nd order parametric oscillator. In this figure we use the same synthetic dynamics and the same set of hyperparameter values than those described in the caption of Figure 17. In particular, the phase portraits were computed using the dynamics *x*_1_ in Eq. A.18 which only takes into account the effect of the intrinsic noise, that is, without including the extrinsic (i.e. of observation) noise *η*. (A) Phase portrait corresponding to the dynamics shown in Figure 17A,B (No PFC). (B) Phase portrait corresponding to the dynamics shown in Figure 17H,I (No PFC). (C) Phase portrait corresponding to an intermediate dynamics in between the cases shown in Figure 17A,B and Figure 17D,E. (D) Phase portrait corresponding to the dynamics shown in Figure 17K,L (Non harmonic PFC). (E) Phase portrait corresponding to the dynamics shown in Figure 17D,E (Harmonic PFC). (F) Phase portrait corresponding to the dynamics shown in Figure 17F,G (Harmonic PFC).

Figures B.10 and B.11 show the PFC patterns corresponding to the oscillator dynamics generated by simultaneously applying an off-resonance external driving Fe with the parametric driving *W_p_* tuned at the same frequency *f_e_* = *f_p_* = *f*_0_ /11.62 ≈ 8.61 Hz and *θ_e_* = *π*/2 (see Eqs. A.16 and A.17). For the configuration used to compute the Figures B.10J and B.11J, the intrinsic noise is capable to drive the resonator at its natural frequency *f*_0_ for low *A_p_* values (see panels H and I in Figures B.10 and B.11). However, no harmonicity is observed in Figures B.10J and B.11J for low *A_p_* values (see blue filled circles in Figures B.10J and B.11J), due to the fact that we configured the external (*f_e_*) and parametric (*f_p_*) driving frequencies having a non harmonic ratio with the natural resonance frequency (*f*_0_) of the undamped oscillator (*μ* = 0), i.e. *f_e_* = *f_p_* = *f*_0_/11.62 ≈ 8.6 Hz. In this regard, compare the harmonicity for low Ap values (blue filled circles) in Figures 17J, B.10J and B.11J.

**Figure B10:**
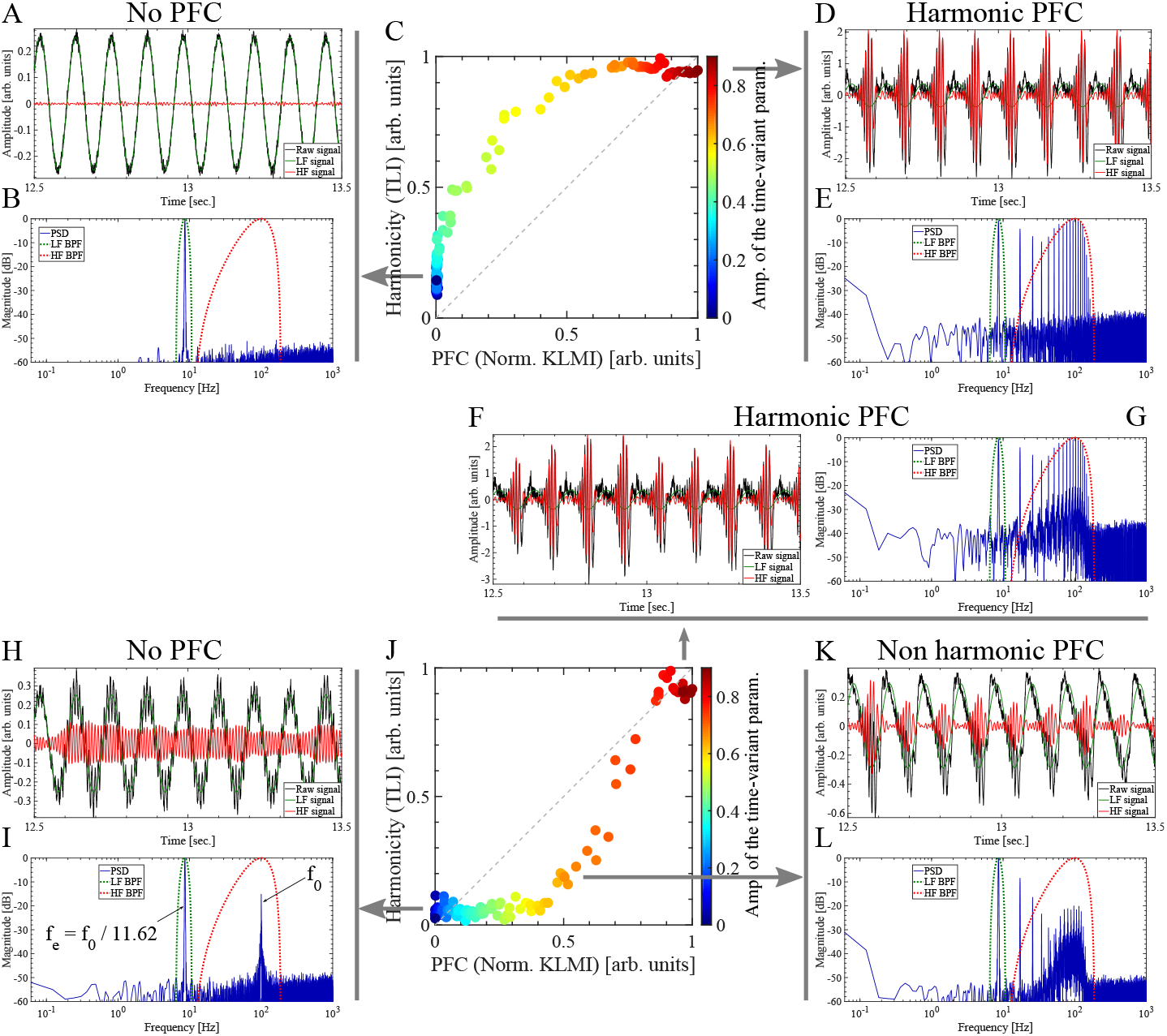
Harmonicity-PFC plot computed for the simulated dynamics of the 2nd order parametric oscillator with intrinsic noise of type additive white Gaussian noise (AWGN). Note that two oscillatory dynamics with independent frequencies can produce harmonic PFC patterns (panels D, E and F, G). In this figure we use the same synthetic dynamics and the same set of hyperparameter values to compute the metrics than those described in the caption of Figure 17, except for the phase of the external driving *θ_e_* = *π*/2 and the frequency of the parametric and external driving, which were configured as *f_p_* = *f_e_* = *f*_0_ /11.62 ≈ 8.3 Hz (i.e. *f_p_* and *f_e_* are non harmonics of *f*_0_). The harmonicity metric (TLI) was computed as it was described in Section 2.4. The PFC metric (*KLMI_PFC_*) was computed using Eqs. 6 and 7 with the configuration given by Eq. A.22. Note that the *KLMI_PFC_* was normalized with its maximum value in each plot.

**Figure B11:**
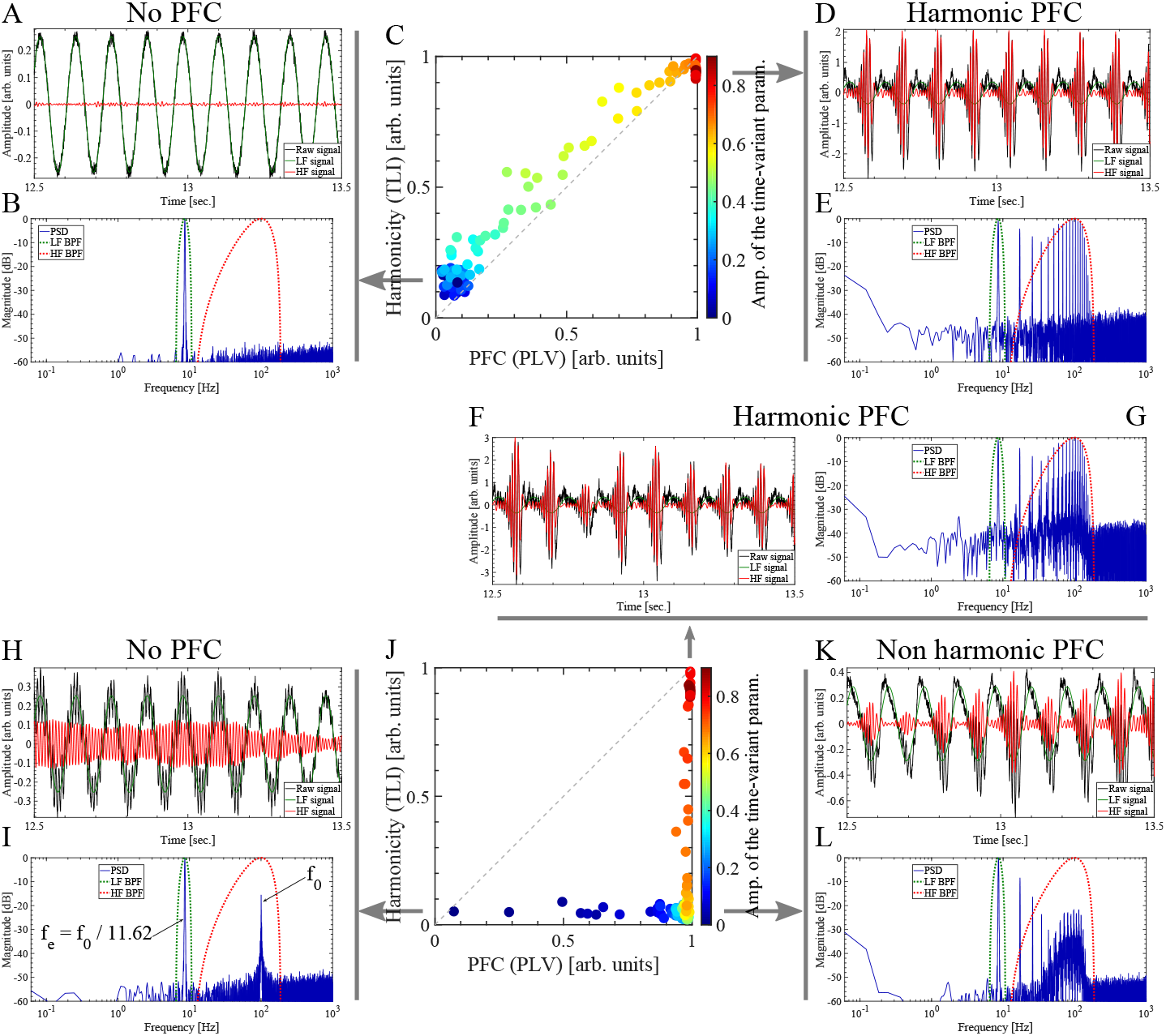
Harmonicity-PFC plot computed for the simulated dynamics of the 2nd order parametric oscillator with intrinsic noise of type additive white Gaussian noise (AWGN). Note that two oscillatory dynamics with independent frequencies can produce harmonic PFC patterns (panels D, E and F, G). In this figure we use the same synthetic dynamics and the same set of hyperparameter values to compute the filtering and harmonicity metric (TLI) than those described in the caption of Figure 17, except for the phase of the external driving *θ_e_* = *π*/2 and the frequency of the parametric and external driving, which were configured as *f_p_* = *f_e_* = *f*_0_ /11.62 ≈ 8.3 Hz (i.e. *f_p_* and *f_e_* are non harmonics of *f*_0_). The harmonicity metric (TLI) was computed as it was described in Section 2.4. In this case, the PFC metric (*PLV_PFC_*) was computed using Eq. 4 with the configuration given by Eq. A.22 and *M* = 1, *N* = 1.

### Appendix B.5. Biologically plausible neural network model

Figure B.12 shows the harmonicity-PAC plots computed for the for the simulated dynamics of the biologically plausible neural network model shown in Figure 1 using the softplus activation function *S*(*I_i_*) (Eq. 3). The results shown in Figure B.12 should be compared with those shown in Figure 18 of the main text.

**Figure B12:**
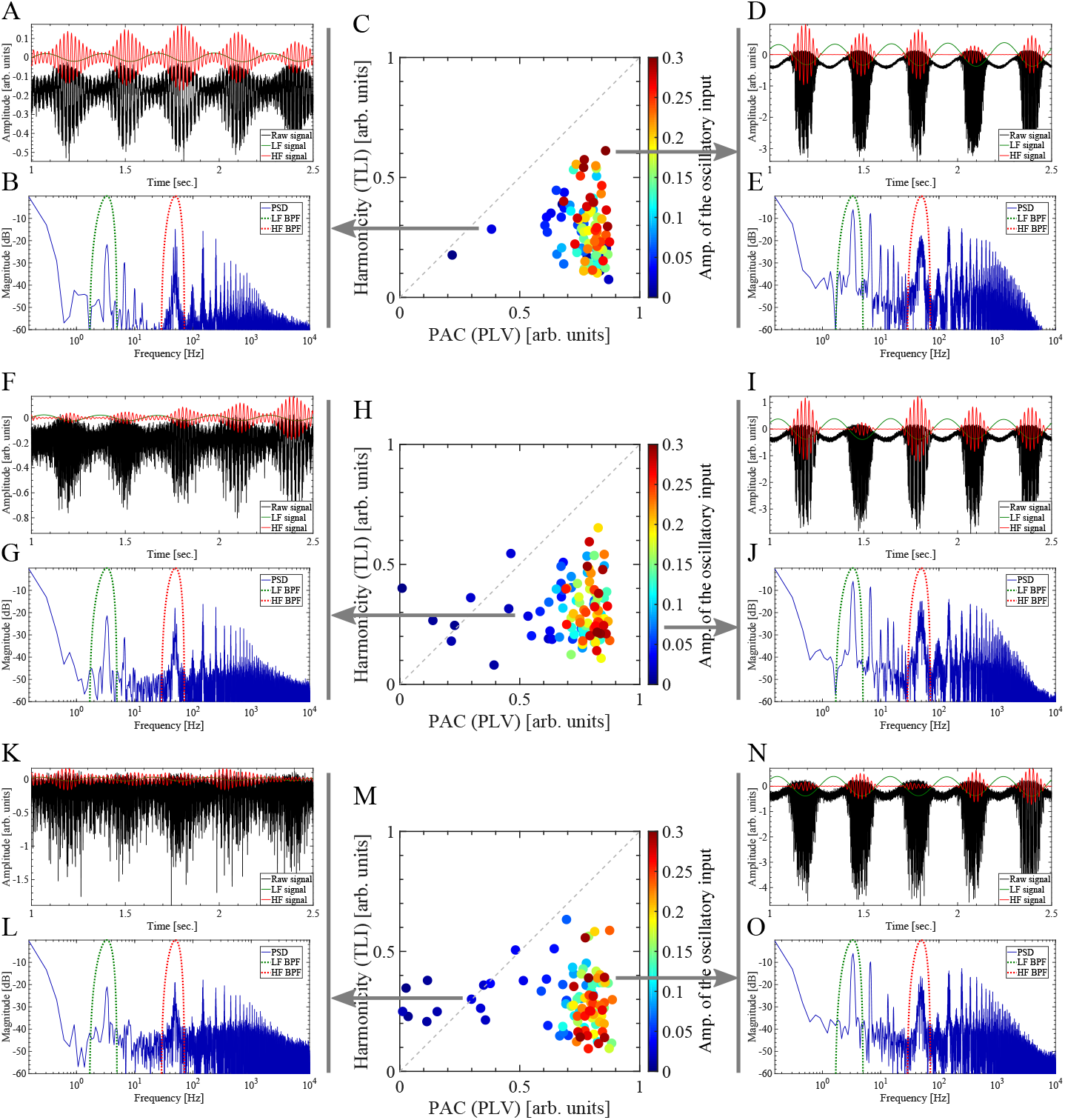
Harmonicity-PAC plot computed for the simulated dynamics of the biologically plausible neural network model shown in Figure 1 using the softplus activation function *S*(*I_i_*) (Eq. 3). In this figure we use the same synthetic dynamics and the same set of hyperparameter values to compute the metrics than those described in the caption of Figure 18, except for the activation function *S*(*I_i_*) which in this case was computed using the Eq. 3 with *c* = 20.

## Notes

https://github.com/damian-dellavale/Time-Locked-Index/

### Competing Interest Statement

The authors have declared no competing interest.

